# Prediction of Stroke Outcome in Mice Based on Non-Invasive MRI and Behavioral Testing

**DOI:** 10.1101/2022.05.13.491869

**Authors:** Felix Knab, Stefan Paul Koch, Sebastian Major, Tracy D. Farr, Susanne Mueller, Philipp Euskirchen, Moritz Eggers, Melanie T.C. Kuffner, Josefine Walter, Daniel Berchtold, Samuel Knauss, Jens P. Dreier, Andreas Meisel, Matthias Endres, Ulrich Dirnagl, Nikolaus Wenger, Christian J. Hoffmann, Philipp Boehm-Sturm, Christoph Harms

## Abstract

**Background:** Prediction of post-stroke outcome using the degree of subacute deficit or magnetic resonance imaging is well studied in humans. While mice are frequently used animals in preclinical stroke research, systematic analysis of outcome predictors is lacking.

**Methods:** We introduced heterogeneity into our study to broaden the applicability of our prediction tools. We analyzed the effect of 30, 45 and 60 minutes of arterial occlusion on the variance of stroke volumes. Next, we built a heterogeneous cohort of 215 mice using data from 15 studies that included 45 minutes of middle cerebral artery occlusion and various genotypes. Motor function was measured using the staircase test of skilled reaching. Phases of subacute and residual deficit were defined. Magnetic resonance images of stroke lesions were co-registered on the Allen Mouse Brain Atlas to characterize stroke topology. Different random forest prediction models that either used motor-functional deficit or imaging parameters were generated for the subacute and residual deficits.

**Results:** Variance of stroke volumes was increased by 45 minutes of arterial occlusion compared to 60 minutes and including various genotypes. We detected both a subacute and residual motor-functional deficit after stroke and different recovery trajectories. In mice with small cortical lesions, lesion volume was the best predictor of the subacute deficit. The residual deficit was most accurately predicted by the degree of the subacute deficit. When using imaging parameters for the prediction of the residual deficit, including information about the lesion topology increased prediction accuracy. A subset of anatomical regions within the ischemic lesion had particular impact on the prediction of long-term outcome.

**Conclusions:** We developed and validated a robust tool for the prediction of functional outcome after stroke in mice using a large heterogeneous cohort. Study design and imaging limitations are discussed. In the future, using outcome prediction can improve the design of preclinical studies and guide intervention decisions.

## INTRODUCTION

Stroke remains a major public health challenge, as its occurrence rises in aging societies.^1^ With decreasing mortality rates due to progress in acute stroke therapy, the need to develop treatments that improve functional outcome has become more pressing.^2^ Understanding predictors of functional outcome after stroke both in humans and rodents is an essential step towards defining adequate treatment decisions and improving the design of preclinical studies.^3^ Prediction of expected motor-functional outcomes in mice could potentially guide researchers in their treatment decisions in preclinical stroke intervention studies and support outcome-dependent stratifications. In humans, magnetic resonance imaging (MRI) is used to assess stroke volume and topology.^4^ Lesion volume is most commonly used to analyze stroke severity and predict functional outcome, but the prognostic value has been shown to be better when including lesion topology.^5–11^ This was also shown in an exploratory preclinical study using a porcine stroke model.^12^ Additionally, the degree of the subacute deficit has been shown to be a reliable predictor of long-term outcome, especially for mild functional deficits.^13,14^ In contrast to humans, little is known about predictors of functional outcome after stroke in mice, although they are commonly used in preclinical research. Detecting motor-functional deficits in rodents following experimental stroke is difficult as they show impressive recovery and have subtle, if any, long-term motor deficits in common behavioral tests.^15^ The stroke preclinical assessment network (SPAN) recently completed the largest preclinical stroke study on mice and provided examples of how the middle cerebral artery occlusion (MCAO) model can be used to conduct studies that are similar to those performed in humans.^16^ Various efforts to standardize preclinical stroke models have been made, including standardization of operation procedures and monitoring of cerebral blood flow.^17,18^ Today, heterogeneity of the MCAO model has been reduced but still largely depends on factors such as the experience of the surgeon or filament choice.^19^ Additionally, other factors such as the natural microbiome, comorbidities, age or sex are likely contributing to the observable heterogeneity in preclinical stroke studies. In their most recent publication, the SPAN consortium has suggested to embrace and systematically integrate this heterogeneity of outcomes into preclinical research, to more accurately simulate the reality of clinical studies.^19^ In a previous study, we have investigated accuracy of early perfusion/diffusion MRI for final infarct prediction.^20^ The main goal of the present study was to identify and compare different predictors of functional outcome following brain ischemia in mice and to provide researchers with a tool, that can help to cope with heterogeneity of stroke volumes. In order to incorporate increased levels of heterogeneity, we first tested the effect of arterial occlusion on the heterogeneity of stroke volumes by comparing 30 minutes of MCAO to 45 and 60 minutes. We further tested the effect of genotype on the stroke volume. Next, we aggregated imaging and behavioral data from 15 MCAO studies and developed a prediction tool that uses early MR imaging and behavioral testing to predict the expectable functional outcome. Heterogeneity of stroke volumes was incorporated by choosing the 45 minutes of MCAO and including 15 different genotypes. Using a newly modified protocol for the staircase test of skilled reaching, we tested forepaw function before and after stroke over a period of 41 days. We defined two phases of post-stroke recovery and tried to predict functional outcome after one week (subacute outcome) or three weeks (long-term outcome) using early MRI. We hypothesized that incorporating lesion topology and behavioral data would improve the predictability of post-stroke outcomes compared to using only the lesion volume. Therefore, we compared the prognostic value of lesion volume and topology for predicting the *subacute deficit* and additionally included the *subacute deficit* for predicting the *residual deficit* using machine learning on a large cohort of 148 mice. We replicated our findings in an independent cohort. Our work opens new avenues towards a better understanding of post-stroke recovery and impairment in mice and paves the way to using prediction of functional outcome for the optimization of preclinical stroke studies. The provided prediction tool gives laboratories, in particular those that are just beginning to gradually increase standardization of the MCAO model in an iterative process, the option to cope with heterogeneity of stroke volumes from various sources.

## METHODS

### Heterogeneity of Stroke Volumes

The effect of arterial occlusion time on the heterogeneity of stroke volumes was assessed using data from a total of 420 mice undergoing either 30 (n=114 C57Bl/6 wildtype), 45 (n=56 C57Bl/6 wildtype, n=129 C57Bl/6 transgenic) or 60 (n=121 C57Bl/6 wildtype) minutes of MCAO. Mean lesion volumes and deviation from group mean in percent was assessed. As the 45 minutes MCAO group included 15 different genotypes, we further analyzed the effect of the co-factors genotype, age and sex on the stroke volume using mixed effects ANOVA.

### Experimental Design

Data from 13 studies performed between 2016 to 2018 were pooled for behavioral and imaging analyses. Mice were trained in the staircase test for three weeks before and after 45 min of MCAO surgery followed by MRI after 24 hours (Figure 2A, details in Supplemental Materials & Methods). 360 mice were considered for inclusion into this study. Four exclusion criteria were pre-specified (Figure S1): **1)** In the third week of testing, mice performed below 20% of the average in the staircase test (n=30); **2)** no visible infarct in MRI (n=18); **3)** Surgical death or humane euthanasia before study end (n=121) and **4)** incomplete data sets for behavioral data (n=25). Following these criteria, 166 mice (MCAO: n=148, sham: n=18) were included in the cohort referred to as *prediction cohort* (Table S1). Finally, to validate our prediction models, an additional cohort was built, termed *replication cohort*, using two studies from 2015 and 2019 with n=49 mice (MCAO: n=37, sham: n=12) finally included (Figure S2). Basic characteristics of all animals are provided in Table S2 and Figure S3.

### Staircase Test of Skilled Reaching

Behavioral testers were blinded to the intervention and MRI results. Mice were housed, food restricted and familiarized with the staircase test (details in Supplemental Materials & Methods).^21^ We chose one experimental variable, the percentage of collected pellets per side. Performance was expressed as a percentage of average pre-stroke performance (*baseline*). Mice were randomized and allocated to either MCAO or sham surgery. Based on the mean performance of the entire cohort after stroke, two phases were defined: a ‘*subacute deficit*’ phase (days 2–6), during which performance appeared dynamic and increasing, and a ‘*residual deficit*’ phase (days 12–21), throughout which performance appeared stable. Average performance was calculated for each mouse for the respective period. A gap of five days was chosen between the *subacute* and *residual deficit* to ensure the dynamic phase had ended.

### Classification of Severity Grades for Subgroup Analysis

Alluding to the modified Rankin scale, a score regularly used in stroke patients, five different functional subgroups were defined.^22^ Subgroups were based on the severity of the motor-functional deficit of the paretic paw: mild (performance >80%), moderate-mild (60–79%), moderate (40–59%), moderate-severe (20–39%) and severe deficit (0–19%). Both *subacute* and *residual deficit* were used to define subgroups, depending on the analysis context (Figure 2B–D).

### Selection of Predictors

Two factors were analyzed for their predictive value for the degree of the *subacute deficit* after MCAO: **1)** stroke *lesion volume* measured within the coordinates of the Allen Mouse Brain Atlas (AMBA), which makes it identical to the volume of remaining brain tissue,^23^ and **2)** MRI lesion percentages measured in the 736 AMBA regions. Throughout the manuscript, this predictor is referred to as *segmented MRI*. The dimensionality of the *segmented MRI* (736 initial regions) was reduced by discarding all regions that did not show a lesion in at least one animal. This left the *segmented MRI* with 536 regions. In addition to those two imaging-based parameters, for the prediction of the *residual deficit*, the degree of *subacute deficit* was included as a third factor.

### Development of Prediction Models with Machine Learning

For development of machine learning based prediction models, we divided the *prediction cohort* into a training (2/3 of the animals) and a testing (1/3 of the animals) cohort (details in Supplemental Materials & Methods). For each predictor, we calculated the median absolute error (MedAE) and report the average as *prediction error* (*PE*). Only *PEs* from the test cohort were used for accuracy analysis. *PE* measures the difference between predicted and actual performance. Small errors indicate high accuracy. First and third quartiles and interquartile ranges (IQR) of absolute error were calculated to estimate *PE* variation within a cohort. To determine which anatomical regions affect MRI-based *residual deficit* prediction accuracy, the *out-of-bag* predictor importance for each anatomical region in the *segmented MRI* was calculated (details in Supplemental Materials & Methods).^24^ The prediction models were uploaded and are available at https://github.com/major-s/mouse-mcao-outcome-predictor/releases/tag/v1.0.0.

### Statistics, Data Availability & Ethics Approval

Data are reported according to the ARRIVE guidelines. Methods for statistics, data display and availability including ethics approval are described in the Supplemental Materials & Methods. Data is openly accessible on zenodo (https://doi.org/10.5281/zenodo.6534690).

## RESULTS

### Occlusion Time and Genotype Influences Heterogeneity of Stroke Volumes

Edema corrected lesion volumes were calculated and stroke incidence maps were generated for the three occlusion times (Figure 1A through 1C). Mean lesion volume after 30 minutes of MCAO was 14.62mm^3^ (95% CI: 11.63-17.60), 31.94mm^3^ (26.01-37.87) after 45 minutes in wildtype C57Bl/6 and 23.54 mm^3^ (19.86-27.23) in the transgenic C57Bl6 mice and 50.21mm^3^ (45.19-55.23) after 60 minutes of MCAO (Figure 1D). Volumes were significantly different between the occlusion times according to Kruskal-Wallis test (chi^2^=131.0, p<0.0001). Deviation from group mean in percent was 72.71% (57.43-87.99) in the 30 minutes MCAO group, 58.26% (48.44-68.98) after 45 minutes in the C57Bl/6 and 71.11% (61.64-80.59) in the Non-C57Bl/6 mice and 44.73% (38.86-50.60) in the 60 minutes MCAO group (Figure 1E). Deviation from group mean was significantly different between the three groups (chi^2^=26.65, p<0.0001). After correction for multiple comparisons using Dunn’s test, difference of deviation from group mean between the 30 and 60 (Z=3.12, p=0.011) and between the 45 and the 60 minutes MCAO (wildtype C57Bl/6: Z=2.77, p=0.034; transgenic C57Bl/6: Z=5.09, p<0.0001) was statistically significant. Next, we assessed the effect of genotype, sex and age on the stroke volume in the 45 minutes MCAO group. Mixed effects ANOVA showed that genotype had a statistically significant effect on the stroke volume (F(17, 165)=1.909, p=0.020), while sex (F(1, 165)=0.139, p=0.710) and age (F(1, 165)=0.060, p=0.806) did not.

**Figure 1.**
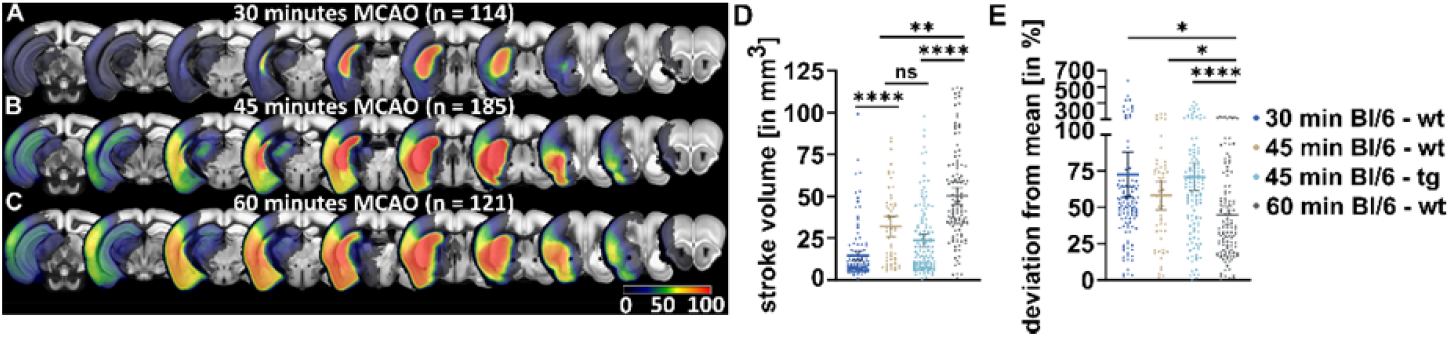
Heterogeneity of Stroke Volumes By Occlusion Time. **A**, Lesion incidence maps of mice with 30, **B**, 45 or **C**, 60 minutes of MCAO. Notably, an increasing percentage of the mice show affection of cortical regions and therefore of the entire MCA territory. **D**, Stroke volumes resulting from the three different arterial occlusion times. The 45 minutes MCAO group included both C57Bl/6 wildtype (wt) and 14 different transgenic (tg) mice and were therefore analyzed separately. Lesion volumes became significantly larger with longer occlusion times. Error bars indicate 95% confidence interval, black line indicates mean. **E**, Individual deviation from group mean was expressed in percent. Interestingly, the 60 minutes of MCAO provided significantly smaller deviations, indicating more homogeneity of stroke volumes in longer occlusion times.

**Figure 2.**
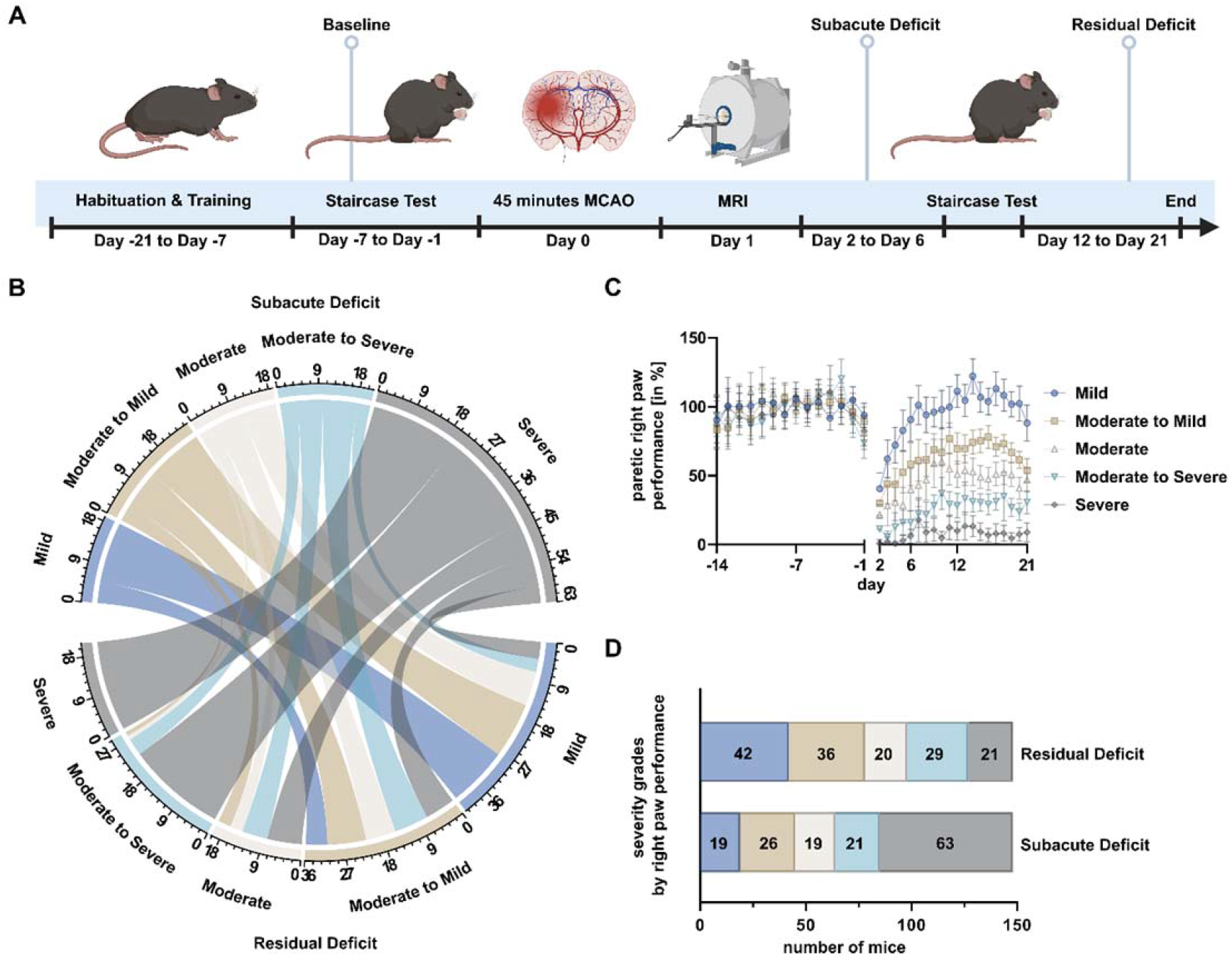
Experimental Design and Assessment of Functional Subgroups. **A**, Mice were trained in pellet reaching for 21 days. Average percentage of retrieved pellets during the last week prior to MCAO was used to normalize performance (baseline). Mice received either 45 min of MCAO or sham surgery. MRI was performed on day 1 and mice were then tested for another 21 days. **B**, Chord diagram displaying the different trajectories of recovery. Groups on top display mice stratified by degree of *subacute deficit*, while groups on bottom display mice stratified by *residual deficit*. Numbers indicate group sizes. Interconnections visualize how many mice with a given *subacute deficit* recovered to have a given *residual deficit*, i.e. roughly a third of the mice with a severe *subacute deficit* displayed no recovery and showed a severe *residual deficit*. **C**, Performance of the paretic right paw over time for each severity grade. Error bars indicate 95% CI. Severity grades here were based on the degree of the *residual deficit*. **D**, Overview over sizes of functional subgroups. Groups are either based on the degree of *subacute* or *residual deficit*. While the majority of mice displayed a severe *subacute deficit*, due to their impressive capability for spontaneous recovery, the majority of mice displayed only mild *residual deficits*.

### MRI Characterization and Lesion Topology in Prediction and Replication Cohort

T2-weighted MR-images were co-registered on the AMBA to evaluate lesion topology (*segmented MRI*) and incidence maps were generated (Figure S4A and S4B). Quantitative evaluation of the hyperintense lesion was performed on 536 regions and lesion volume in each region was expressed in percent of the total region (Table S3). Lesion volumes and topological characteristics varied slightly between the *prediction* and *replication cohort* (Figure S4C and S4D). Further, incidence maps and lesion volumes were calculated for the functional subgroups (Figure S5) and more details can be found in the Supplemental Materials & Methods.

### A Modified Staircase Test Can Assess Early and Long-Term Functional Outcome in Mice after Stroke

Performance peaked after approximately 7 days (Figure 3A–B). The average percentage of retrieved pellets during *baseline* testing was 49.26% (47.22–51.31) on the left (later non-paretic) side and 46.29% (44.16–48.42) on the right (later paretic) side in the MCAO group and 45.81% (37.38–54.23) and 47.11% (38.95–55.26) on the left and right sides in the sham group (Figure S6). Mixed effects analysis showed no side-group interaction (F(1,164)=1.464, p=0.2281). After MCAO, mean subacute performance was 61.60% (56.12–67.08) on the non-paretic side and 37.59% (31.92–43.26) on the paretic side (Figure 3C). In sham animals, subacute performance was 78.40% (63.46–93.34) on the left and 72.23 (57.30–87.15) on the right side. Mixed-effects analysis found a significant interaction between side and group (F(1,164)=4.189, p=0.0423). After correcting for multiple comparisons using Šidák’s test, performance of the paretic paw was significantly lower in MCAO animals when compared to sham (LSMD=34.63pp, t=4.092, DF=328, p=0.0001). Non-paretic sides did not differ between groups (LSMD=16.80pp, t=1.985, DF=328, p=0.0937). Paired t-test was performed to test for a difference in between paretic and non-paretic paws of the MCAO mice to show side-specificity of the motor-functional deficit. Performance in the paretic paw was significantly lower than in the non-paretic paw (MD: 24.01 %, 18.34–29.68, t=8.365, df=147, p<0.0001) (Figure S7A).

**Figure 3.**
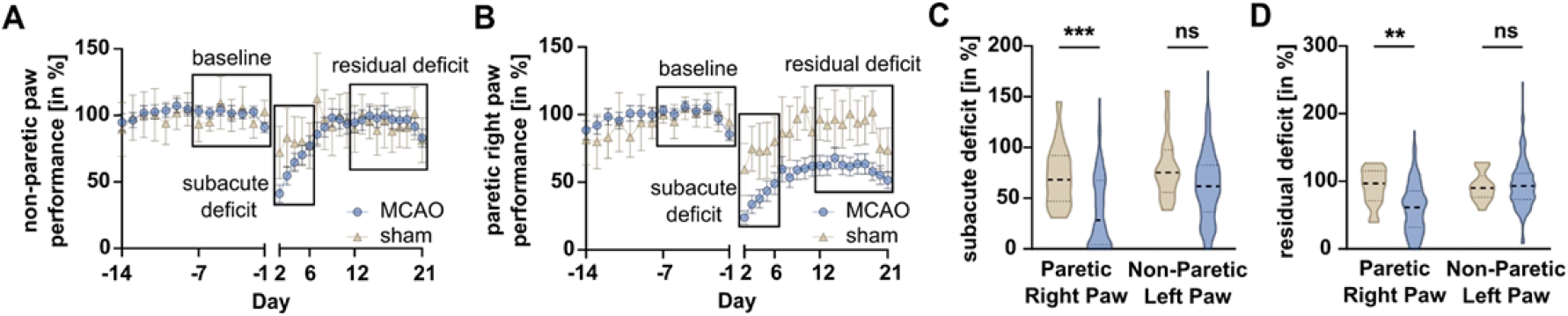
Assessing the Sensorimotor Deficit after Stroke. **A**, Non-paretic left and **B**, paretic right paw performance over time, bars depict 95% confidence interval. Performance on days -7 to -1 were averaged and taken as a baseline (first black rectangle). Data from days 2 to 6 were summarized and referred to as *subacute deficit* (second rectangle) while data from days 12 to 21 are referred to as *residual deficit* (third rectangle). **C**, *Subacute* and **D**, *residual deficit* of the non-paretic and paretic paw in sham (brown plot) and MCAO (blue plot) animals. Dashed lines indicate medians, dotted line indicate quartiles. Mixed-effects model followed by Šidák’s post-hoc tests for multiple comparison were used to test effect of group and side.

In the period of residual deficit, non-paretic performance was 95.35 % (89.54–101.17) and paretic performance was 60.88 % (54.94–66.81; Figure 3D). During the phase of residual deficit, sham animals performed 92.39% (81.80–102.98) on the non-paretic side and 91.56% (78.45–104.66) on the paretic side. Mixed-effects analysis showed a significant interaction between side and group (F(1,164)=9.384, p=0.0026). Using Šidák’s test, paretic paw performance was significantly lower in MCAO animals compared to sham (LSMD=30.68pp, t=3.501, DF=328, p=0.0011). Non-paretic sides showed no statistically significant difference between groups (LSMD=2.96pp, t=0.3377, DF=328, p=0.9302). Finally, we analyzed the effects of the general well-being of the animals on functional outcome. We used the Modified DeSimoni Neuroscore for the assessment of general deficits and the weight loss on day 2 as proxies for general well-being and found poor correlation to the degree of either subacute or residual deficit (Figure S8 and S9).^25^ In addition, *PEs* of weight loss or neuroscore were significantly higher compared to imaging-or performance-based prediction (Figure S10 and S11).

### Prediction of Subacute Deficit from Acute MR Imaging

Using random forest and early non-invasive MRI data, *subacute deficit* prediction models were created. *PE* was 13.30pp (13.00–13.59, Q_1_: 5.98, Q_3_: 31.48, IQR: 25.51) for the *lesion volume* and 16.23pp (16.03–16.42, Q_1_: 8.04, Q_3_: 29.18, IQR: 21.14) for the *segmented MRI* (Figure 4A–B). Unpaired, two-tailed t-test revealed, that *PE* was significantly smaller when using the *lesion volume* as a predictor compared to the *segmented MRI* (MD=2.93pp, 2.58– 3.28, t=16.60, df=98, p<0.0001).

**Figure 4.**
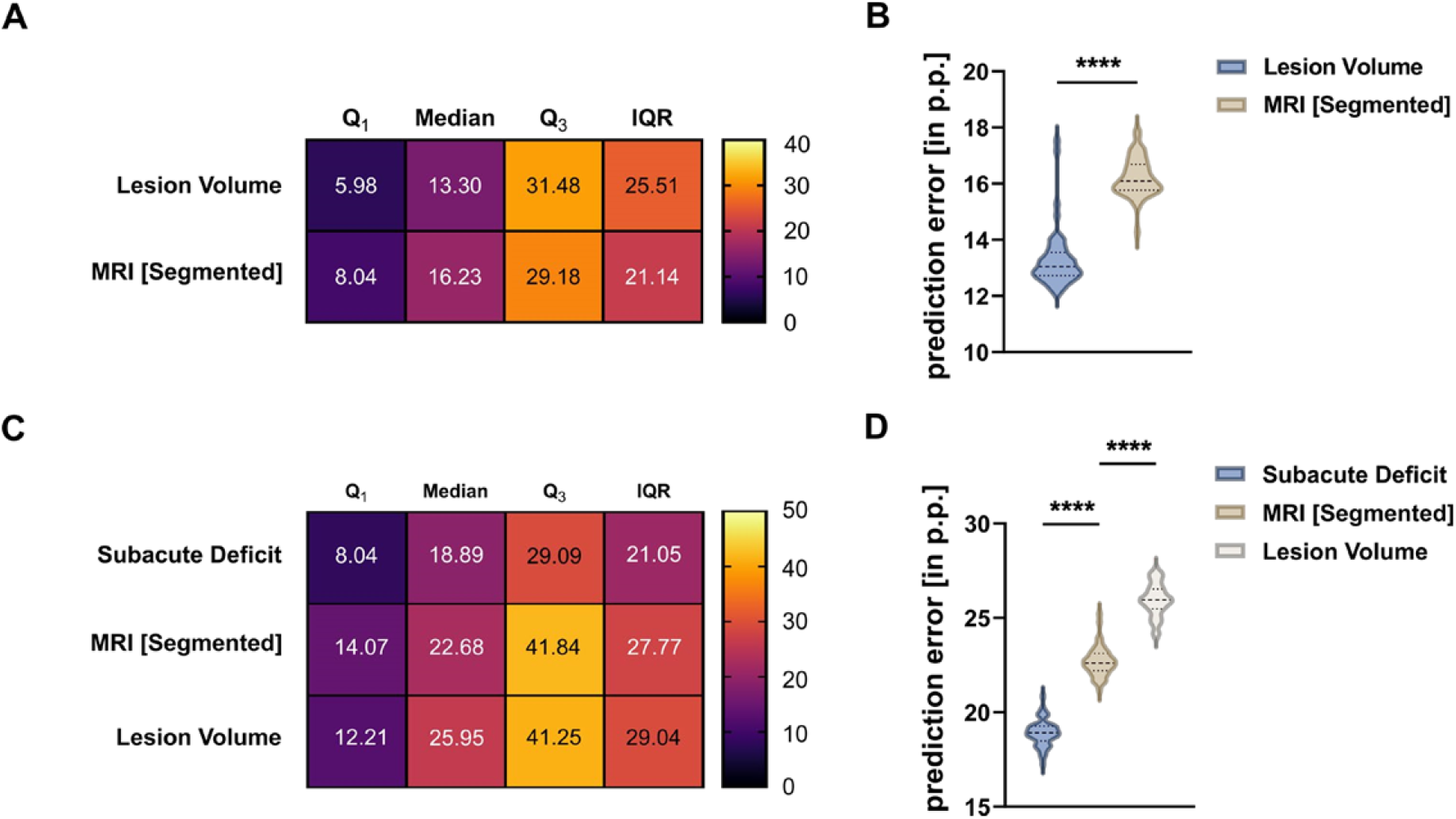
Comparison of Predictors of the Subacute and Residual Deficit. **A** through **B**, Comparison of the prediction accuracy of the two predictors of the *subacute deficit*. Averaged Q_1_, median, Q_3_ and IQR were calculated using the 50 independent models for each predictor. Dashed lines indicate median, dotted lines indicate quartiles. Unpaired t-test was used to compare *PEs*. Small *PE* indicates small deviation between predicted and achieved performance. **C** through **D**, Comparison of prediction accuracy of the two imaging based parameters and the *subacute deficit* for the prediction of the *residual deficit*. Again, one-way ANOVA followed by Tukey’s test for multiple comparison was used.

### Prediction of Residual Deficit from Subacute Deficit and MRI

*PE* was 18.90pp (18.70–19.09, Q_1_: 8.04, Q_3_: 29.09, IQR: 21.05) for the *subacute deficit*, 22.68pp (22.46–22.90, Q_1_: 14.07, Q_3_: 41.84, IQR: 27.77) for the *segmented MRI* and 25.95pp (25.71–26.29, Q_1_: 12.21, Q_3_: 41.25, IQR: 29.04) for the *lesion volume* (Figure 4C–D). One-way ANOVA showed predictors differed significantly (F(2,147)=1032, p<0.0001). After multiple comparisons were corrected using Tukey’s test, the *PE* of the *subacute deficit* was significantly smaller than *PE* of the *segmented MRI* (MD=3.79pp, 4.15–3.42, p<0.0001) and the *lesion volume* (MD=7.06pp, 7.43–6.69, p<0.0001). *Segmented MRI* outperformed the prediction accuracy of the *lesion volume* (MD=3.28pp, 3.65–2.91, p<0.0001).

### Identification of Anatomical Regions Associated with Long-Term Outcome

To better understand which anatomical regions within the lesion best predict long-term outcome, the contribution of each region measured by MRI was analyzed by quantifying its importance within the random forest algorithm. We found 81 regions with above-zero out-of-bag predictor importance, meaning they contribute more than random data (Figure 5A–B, Table S4). Among those regions were the caudoputamen, the primary somatosensory area, the external capsule and the corticospinal tract. In the next step, random forest models were trained using increasing number of MRI regions, starting with the most important region. The *PE* of those models gradually decreased with increasing number of MRI regions (Figure 5C). *PE* quickly reached a plateau, and adding more regions only slightly improved accuracy. A relevant local minimum of *PE* was observed with the first 14 anatomical regions (*PE*=22.91pp).

**Figure 5.**
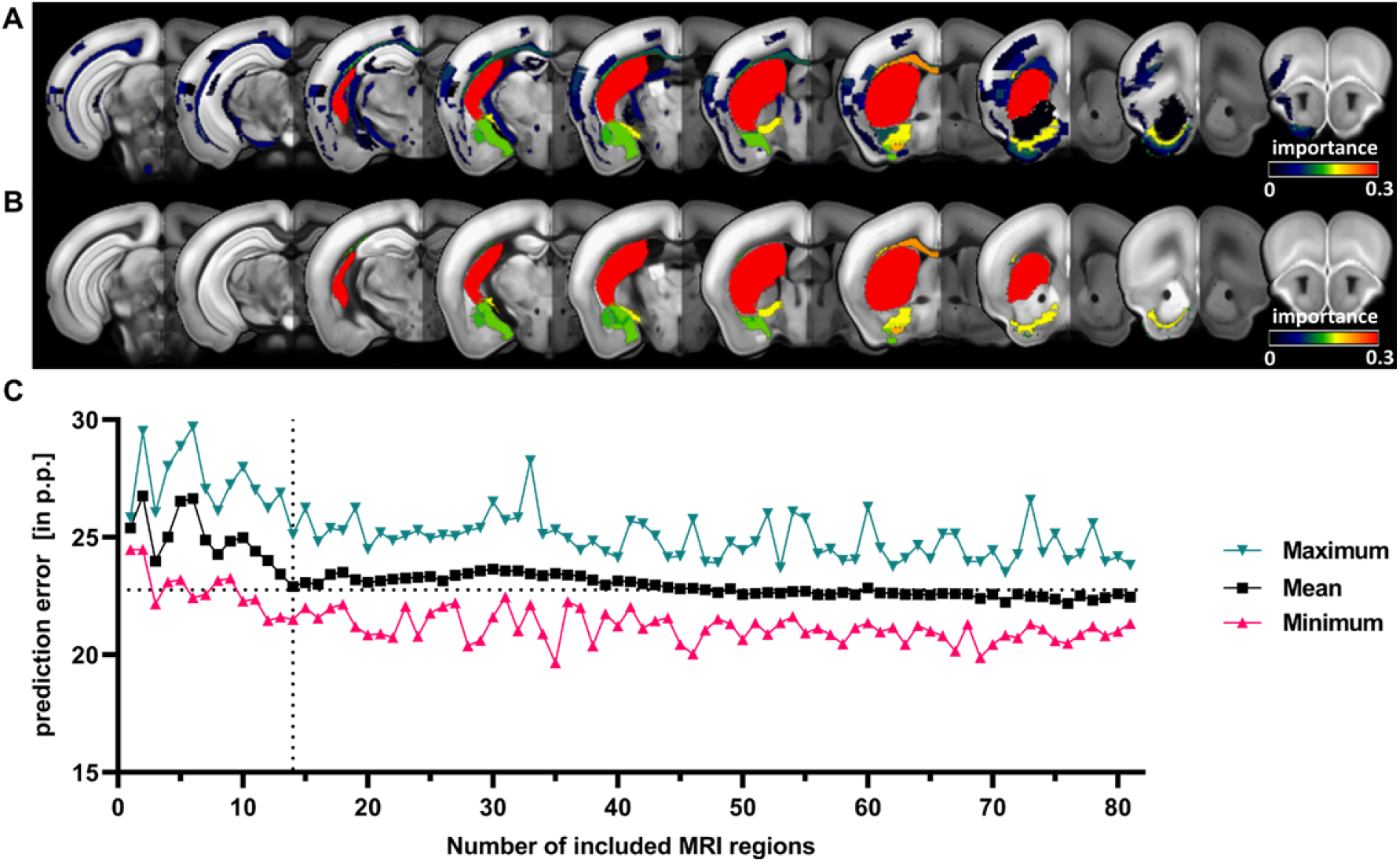
Identification of Anatomical Regions that Drive Imaging-Based Prediction of Functional Outcome. **A**, Coronal slices depicting the importance of the 81 regions calculated from our random forest models (RF). Only regions with an importance level >0 are shown **B**, Visualization of the importance level of the 14 anatomical regions that were included in the prediction model that resulted in a local minimum of *PE*. **C**, Prediction of *residual deficit* was performed using an increasing number of regions in the *segmented MRI. PE* as well as maximum and minimum *PE* are depicted. Dashed lines at x=14 and y=21.5 denote the first local minimum of *PE* when including only 14 regions.

### Validation in an Independent Cohort

As a final step, we tested our prediction models on the independent *replication cohort*. Imaging and behavioral analyses can be found in the Supplemental Materials & Methods. For the prediction of the *subacute deficit*, mean *PE* when using the *lesion volume* was 23.19pp (22.85–23.53) and 20.00pp (19.74–20.27) for the *segmented MRI* (Figure 6A–B). In contrast to our first cohort, now the *segmented MRI* provided significantly smaller *PEs* compared to the *lesion volume* (MD=3.19pp, 3.61–2.76, t=14.89, df=98, p<0.0001). The *PE* for the *residual deficit* using the *subacute deficit* was 10.76pp (10.31–11.22), 22.96pp (22.15–23.77) for the *segmented MRI* and 23.94pp (23.41–24.46) for the *lesion volume* (Figure 6C–D). One-way ANOVA revealed, that there was a statistically significant difference between predictors (F(2,147)=18.98, p<0.0001). After correction for multiple comparisons using Tukey’s test, the *PE* of the *subacute deficit* was significantly smaller than the *PE* of the *segmented MRI* (MD=12.20pp, 13.23–11.17, p<0.0001) and the *lesion volume* (MD=13.18pp, 14.20–12.15, p<0.0001). The difference between s*egmented MRI* and the *lesion volume* was less clear than in our first cohort, but trended towards significance, with the *segmented MRI* providing slightly smaller PEs compared to the *lesion volume* (MD=0.98pp, 2.01–0.05, p=0.0668). Lastly, we tested the prediction accuracy when including an increasing number of anatomical regions. Only regions we identified to have an impact on the prediction of the *residual deficit* in our *testing cohort* were used. Surprisingly, in our *replication cohort*, using the lesion percentage in the caudoputamen provided smaller *PE*s compared to using the entire *segmented MRI* or the *lesion volume*. When adding an increasing number of regions, we first observed a sharp spike in *PEs*, followed by a slow decrease. Excitingly, after including fourteen regions, we again observed a local minimum of *PE* (Figure 6E), replicating our finding from the *testing cohort*.

**Figure 6.**
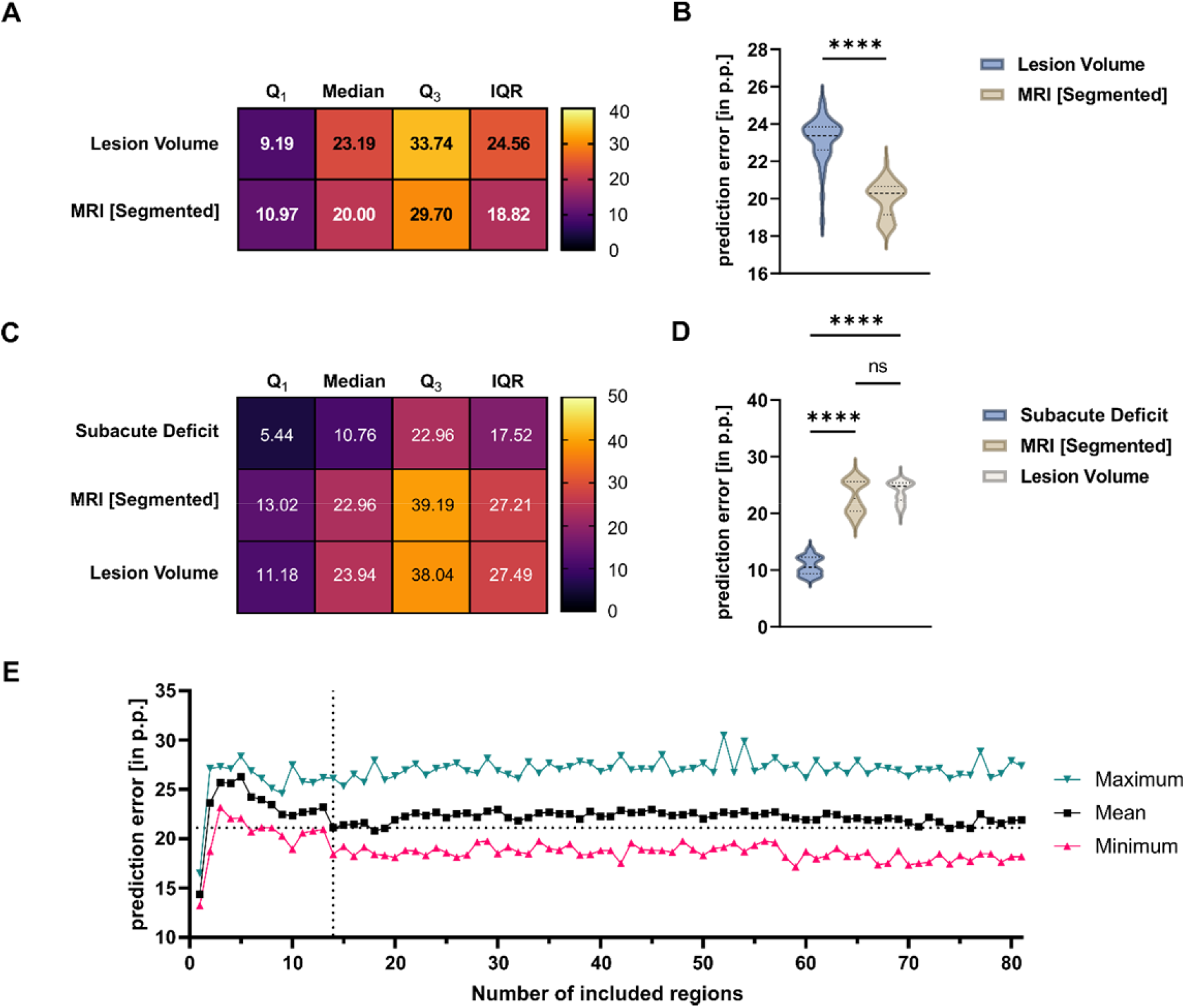
Validation of Prediction Models in an Independent Cohort. **A**, through **B**, Comparison of the prediction accuracy of the two predictors of the *subacute deficit*. Dashed lines indicate median, dotted lines indicate quartiles. Unpaired t-test was used to compare *PEs*. While *PEs* were comparable to the ones seen in our testing cohort, subacute deficit was best predicted when using the segmented MRI **C** through **D**, Comparison of prediction accuracy of the two imaging based parameters and the *subacute deficit* for the prediction of the *residual deficit*. Again, the *subacute deficit* was the best predictor of the *residual deficit*, followed by the *segmented MRI* and the *lesion volume*. **E**, Analogous to the testing cohort, prediction of *residual deficit* was performed using an increasing number of regions in the *segmented MRI*. Dashed lines at x=14 and y=21.5 denote the local minimum of *PE* that had previously been observed in the testing cohort.

## DISCUSSION

In this study, we systematically compared different predictors of subacute and long-term functional outcomes after stroke in a large cohort of mice. We showed that the heterogeneity in our model was driven by occlusion time and genotype. We continued to work with 45 minutes of MCAO, thereby embracing the heterogeneity of stroke volumes and functional outcome. This step widens the dynamic range of our prediction tool and integrates translationally relevant levels of variability into our experimental design. Although these levels are not inherent to the MCAO model if performed in young adult, male C57Bl/6 wildtype mice, the selected approach expands the applicability in studies with heterogeneity of various sources. We used MRI derived parameters such as the *lesion volume* and *lesion topology*, as well as early functional testing to predict post-stroke performance in a pellet reaching task. Studies with large sample sizes have transformed preclinical MRI research and corresponding open data repositories have proven extremely valuable, e.g. in the context of robust functional networks measured by resting state fMRI.^26^ Predictors of post-stroke outcome, such as imaging-derived parameters or functional data, have previously been extensively studied in humans.^2–6,8^ While one preclinical study on predictors of stroke outcome in a porcine MCAO model exists, evidence on prediction of stroke outcome in mice is scarce, despite rodents being used most commonly as model animals in stroke studies.^12,27^ Using a modified protocol for the staircase test of skilled reaching in mice, we were able to monitor motor function daily and to detect both subacute and long-term functional deficits.^21,28^ A side-specific disability of the forepaw was detected over three weeks post-surgery. This indicates that performance was governed by the stroke and not by general sickness behavior. We observed distinct trajectories of functional recovery. The differentiation between recoverees and non-recoverees is comparable to what can be observed in humans.^14^ The resemblance to the human recovery process supports the predictive value of stroke modeling in mice and the case for implementing sophisticated behavioral tests that involve a longer phase of training prior to stroke for assessing long-term impairments.^29^ For future applications, one has to consider the possible impact of food restriction included as part of the staircase training and testing on mortality and lesion volume.^30^

The *lesion volume* was the best predictor for the *subacute deficit* after stroke in mice as measured by *PE* in our *testing cohort*. In the *replication cohort* it was outperformed by the *segmented MRI*, which included information about the lesion topology. While *PEs* remained low, this is clearly a difference between the two cohorts. However, we also observed differences in stroke size and topology between the two cohorts. In our *replication cohort* lesion volumes were bigger and cortical regions accounted for higher proportions of the *lesion volume* compared to the *testing cohort*. While it has previously been stated that lesion volume functions as a good predictor for acute stroke outcome, our data suggests that in mice with greater cortical lesions taking into account topological information provides a more accurate prediction of early functional outcome.^6^ For the long-term motor-functional outcome, in the *testing cohort* the *segmented MRI* provided significantly better prediction accuracies than the *lesion volume*. This indicates that for functional recovery, lesion topology is more relevant than lesion volume, which again is in line with clinical data.^11,12^ It should be noted, that in the 45 min MCAO model, lesion volume and lesion topology are not fully independent predictors. Larger ischemic lesions typically include cortical regions while smaller lesions are restricted to subcortical structures. This phenomenon can be observed in the results of our *replication cohort*, where the difference between the *segmented MRI* and the *lesion volume* was only marginal but trending towards significance.

Following up on the influence of lesion topology, we identified anatomical regions that drive imaging-based prediction of the *residual deficit*. We found that considering only a subset of anatomical regions can explain functional long-term outcome. Prediction accuracy displayed a local maximum after 14 regions had been added to the prediction model. Among those regions were both, cortical and subcortical regions typically affected by a MCAO stroke such as the caudoputamen and the primary somatosensory area. Evidently, the caudoputamen is damaged and part of the lesion core in the transient occlusion model but also serves as the structure that best predicts the performance. Further, white matter regions such as the corpus callosum and the fiber tracts showed to be of particular importance for the prediction of long-term outcome. This is in line with previous studies in humans suggesting that lesions of the corticospinal tract seem to be more important to both early and long-term functional outcome than lesion volume.^31,32^ These findings could be replicated in our independent *replication cohort*, where a local minimum of *PE* could be observed after including the same 14 regions. Again, lesions in the caudoputamen had a major impact on prediction accuracies, outperforming both cortical grey and white matter structures.

Imaging-based predictors offered good estimates of post-stroke long-term outcome but the best predictor of the *residual deficit* was the degree of the *subacute deficit*, again confirming studies on predictors of post-stroke outcome in humans.^33^ Lastly, we found that independent of the chosen model, the prediction accuracy for both the *subacute* and *residual deficits* depended on the degree of the deficit. In line with clinical data, we found that severe *residual deficits* are more difficult to predict than moderate-to-mild deficits.^13,14^

### Study Limitations

We have restricted our prediction tools to 45 min of MCAO using endoluminal filament occlusion. As the data was pooled from 13+2 studies performed between 2015 and 2019, the dataset contained various co-variates. However, variance analysis revealed that sex, genotype, experiment number, or age did not significantly explain differences in performance after stroke. We cannot exclude confounding effects of housing, lab chow diet, anesthesia, or further co-variates. The degree of cerebral blood flow (CBF) reduction based on laser Doppler flow data in the MCA territory was only used during training periods of new surgeons following our standard operating procedure (SOP) for MCAO. Including the reduction of cerebral blood flow as an inclusion criterion, preferably along consensus guidelines such as those developed in the stroke preclinical assessment network (CBF drop by >70% of baseline), can help to reduce the heterogeneity of stroke volumes.^18^ However, our focus was the development of a prediction tool that embraces heterogeneity of stroke volumes in order to more accurately reflect the clinical reality and increase applicability for other research groups. Furthermore, we did not replicate our results with other behavioral readouts than the staircase test of skilled pellet reaching. Additionally, using MR imaging from a later time point, e.g. 21 days after MCAO, would have likely resulted in even better prediction accuracies. However, prediction of functional outcome that is based on late MR-imaging cannot be used for guiding intervention decisions in future preclinical studies as the window of opportunity in mice has proven to be narrow.^34^ Another limitation of our approach is the manual delineation of lesion masks. Although manual lesion segmentation is still the gold standard today, well-documented, multi-centric and openly available stroke T1-weighted MRI datasets pave the way towards fully automated algorithms for MRI stroke lesion segmentation in humans.^35–37^ Results from T2-weighted MRI in rodent stroke models are also promising with excellent similarity between manual segmentation and fully automated algorithms exceeding agreement between multiple raters.^38,39^ These studies have facilitated significant progress in the pursue to fully automate lesion segmentation.^39,40^ Nulling of the fluid signal via e.g. FLAIR MRI could have improved the discrimination between CSF and ischemic lesion. However, 24 hours after MCAO, CSF has still higher signal on RARE than the lesion and the walls of the lateral ventricles can be reliably identified.^23^ For later time points, an adjusted imaging protocol might have to be applied. Animals’ unspecific sickness behavior during the *subacute deficit* phase may have contributed to the degree of performance loss. The differences between sham and MCAO, as well as paretic and non-paretic paw, were, however, significant, indicating that our behavioral paradigm detects stroke-specific motor-functional deficits. There were no video recordings of the testing sessions to account for differences in kinematics but the entire behavioral data set has been uploaded in the repository linked to this study. Analysis of lesion volumes stratified by functional subgroups might be partly weakened by experimental bias, since lesion volume and performance are strongly correlating. Finally, we corrected only for multiple tests in a single test set and not for multiple comparisons between different tests or hypotheses.

## CONCLUSIONS

In conclusion, we have here for the first time identified and characterized predictors of post-stroke outcome in mice. The concordance with clinical data strengthens the usability of mice as model animals in preclinical studies. Prediction of outcome in intervention studies using non-invasive MRI measurements or motor-functional data should be utilized to compare expected and achieved motor-function outcomes, select mice with particularly poor predicted outcomes, and guide treatment or intervention options. This will help to improve the reproducibility of preclinical rodent stroke research and close the translational gap.

## Supporting information

Supplemental Material including Supplemental Material & Methods, Supplemental Results, Figure S1-S14 and Supplemental Table S1-S9

## NON-STANDARD ABBREVIATIONS AND ACRONYMS

AMBA: Allen Mouse Brain Atlas
CCA: Common Carotid Artery
ECA: External Carotid Artery
FOV: Field Of View
ICA: Internal Carotid Artery
IQR: Interquartile Range
LSMD: Least Square Mean Difference
MD: Mean difference
MCA: Middle Cerebral Artery
MCAO: Middle Cerebral Artery Occlusion
MedAE: Median Absolute Error
PA: Pterygopalatine Artery
pp: percentage points
PE: prediction error

## Acknowledgments

Janet Lips coordinated experiments and contributed with MRI, handling, and behavioral evaluation; Larissa Mosch (behavior, histology), Monika Dopatka (surgeries), and Marco Foddis (surgeries, behavioral assessment, and MRI) all provided excellent technical assistance. Figures were created with biorender.com

## Supplemental Materials & Methods

Checklist

Expanded Materials & Methods

Online Figures S1-S13

Tables S1-8

## References

1. Béjot Y, Bailly H, Durier J, Giroud M. Epidemiology of stroke in Europe and trends for the 21st century. Press. Medicale. 2016;45:e391–e398. doi:10.1016/j.lpm.2016.10.003

2. Eriksson M, Norrving B, Terént A, Stegmayr B. Functional Outcome 3 Months after Stroke Predicts Long-Term Survival. Cerebrovasc. Dis. 2008;25:423–429. doi:10.1159/000121343

3. Farr TD, Wegener S. Use of magnetic resonance imaging to predict outcome after stroke: A review of experimental and clinical evidence. J. Cereb. Blood Flow Metab. 2010;30:703–717. doi:10.1038/jcbfm.2010.5

4. Albers GW, Thijs VN, Wechsler L, Kemp S, Schlaug G, Skalabrin E, Bammer R, Kakuda W, Lansberg MG, Shuaib A, et al. Magnetic resonance imaging profiles predict clinical response to early reperfusion: The diffusion and perfusion imaging evaluation for understanding stroke evolution (DEFUSE) study. Ann. Neurol. 2006;60:508–517. doi:10.1002/ana.20976

5. Schiemanck SK, Post MWM, Kwakkel G, Witkamp TD, Kappelle LJ, Prevo AJH. Ischemic lesion volume correlates with long-term functional outcome and quality of life of middle cerebral artery stroke survivors. Restor. Neurol. Neurosci. 2005;23:257–263.

6. Lövblad KO, Baird AE, Schlaug G, Benfield A, Siewert B, Voetsch B, Connor A, Burzynski C, Edelman RR, Warach S. Ischemic lesion volumes in acute stroke by diffusion-weighted magnetic resonance imaging correlate with clinical outcome. Ann. Neurol. 1997;42:164–170. doi:10.1002/ana.410420206

7. Pinto A, McKinley R, Alves V, Wiest R, Silva CA, Reyes M. Stroke lesion outcome prediction based on MRI imaging combined with clinical information. Front. Neurol. 2018;9:1–10. doi:10.3389/fneur.2018.01060

8. Cheng B, Forkert ND, Zavaglia M, Hilgetag CC, Golsari A, Siemonsen S, Fiehler J, Pedraza S, Puig J, Cho TH, et al. Influence of stroke infarct location on functional outcome measured by the modified rankin scale. Stroke. 2014;45:1695–1702. doi:10.1161/STROKEAHA.114.005152

9. Borich MR, Brodie SM, Gray WA, Ionta S, Boyd LA. Understanding the role of the primary somatosensory cortex: Opportunities for rehabilitation. Neuropsychologia. 2015;79:246–255. doi:10.1016/j.neuropsychologia.2015.07.007

10. Munsch F, Sagnier S, Asselineau J, Bigourdan A, Guttmann CR, Debruxelles S, Poli M, Renou P, Perez P, Dousset V, et al. Stroke location is an independent predictor of cognitive outcome. Stroke. 2016;47:66–73. doi:10.1161/STROKEAHA.115.011242

11. Menezes NM, Ay H, Zhu MW, Lopez CJ, Singhal AB, Karonen JO, Aronen HJ, Liu Y, Nuutinen J, Koroshetz WJ, et al. The real estate factor: Quantifying the impact of infarct location on stroke severity. Stroke. 2007;38:194–197. doi:10.1161/01.STR.0000251792.76080.45

12. Scheulin KM, Jurgielewicz BJ, Spellicy SE, Waters ES, Baker EW, Kinder HA, Simchick GA, Sneed SE, Grimes JA, Zhao Q, et al. Exploring the predictive value of lesion topology on motor function outcomes in a porcine ischemic stroke model. Sci. Rep. 2021;11:1–15. doi:10.1038/s41598-021-83432-5

13. Zarahn E, Alon L, Ryan SL, Lazar RM, Vry MS, Weiller C, Marshall RS, Krakauer JW. Prediction of motor recovery using initial impairment and fMRI 48 h poststroke. Cereb. Cortex. 2011;21:2712–2721. doi:10.1093/cercor/bhr047

14. Vliet R, Selles RW, Andrinopoulou E, Nijland R, Ribbers GM, Frens MA, Meskers C, Kwakkel G. Predicting Upper Limb Motor Impairment Recovery after Stroke: A Mixture Model. Ann. Neurol. 2020;87:383–393. doi:10.1002/ana.25679

15. Weber RZ, Mulders G, Kaiser J, Tackenberg C, Rust R. Deep learning based behavioral profiling of rodent stroke recovery. bioRxiv. 2021;

16. Lyden PD, Bosetti F, Diniz MA, Rogatko A, Koenig JI, Lamb J, Nagarkatti KA, Cabeen RP, Hess DC, Kamat PK, et al. The Stroke Preclinical Assessment Network: Rationale, Design, Feasibility, and Stage 1 Results. Stroke. 2022;53:1802–1812. doi:10.1161/STROKEAHA.121.038047

17. Dirnagl U, Group M of the M-S. Standard operating procedures (SOP) in experimental stroke research: SOP for middle cerebral artery occlusion in the mouse. Nat. Preced. 2010;202213:1–14. doi:10.1038/npre.2010.3492.2

18. Taninishi H, Jung JY, Izutsu M, Wang Z, Sheng H, Warner DS. A blinded randomized assessment of laser Doppler flowmetry efficacy in standardizing outcome from intraluminal filament MCAO in the rat. J. Neurosci. Methods. 2015;241:111–120. doi:10.1016/j.jneumeth.2014.12.006

19. Morais A, Locascio JJ, Sansing LH, Lamb J, Nagarkatti K, Imai T, Van Leyen K, Aronowski J, Koenig JI, Bosetti F, et al. Embracing Heterogeneity in the Multicenter Stroke Preclinical Assessment Network (SPAN) Trial. Stroke. 2023;54:620–631. doi:10.1161/STROKEAHA.122.040638

20. Leithner C, Füchtemeier M, Jorks D, Mueller S, Dirnagl U, Royl G. Infarct Volume Prediction by Early Magnetic Resonance Imaging in a Murine Stroke Model Depends on Ischemia Duration and Time of Imaging. Stroke. 2015;46:3249–3259. doi:10.1161/STROKEAHA.114.007832

21. Baird AL, Meldrum A, Dunnett SB. The staircase test of skilled reaching in mice. Brain Res. Bull. 2001;54:243–250. doi:10.1016/S0361-9230(00)00457-3

22. Bamford JM, Sandercock PAG, Wariow CP, Slattery J. Interobserver agreement for the assessment of handicap in stroke patients: To the editor. Stroke. 1989;20:828. doi:10.1161/01.STR.20.6.828

23. Koch S, Mueller S, Foddis M, Bienert T, von Elverfeldt D, Knab F, Farr TD, Bernard R, Dopatka M, Rex A, et al. Atlas registration for edema-corrected MRI lesion volume in mouse stroke models. J. Cereb. Blood Flow Metab. 2019;39:313–323. doi:10.1177/0271678X17726635

24. Breiman L. Random Forests. Mach. Learn. 2001;45:5–32. doi:10.1023/A:1010933404324

25. Donath S, An J, Lee SLL, Gertz K, Datwyler AL, Harms U, Müller S, Farr TD, Füchtemeier M, Lättig-Tünnemann G, et al. Interaction of ARC and DAXX: A novel endogenous target to preserve motor function and cell loss after focal brain ischemia in mice. J. Neurosci. 2016;36:8132–8148. doi:10.1523/JNEUROSCI.4428-15.2016

26. Grandjean J, Canella C, Anckaerts C, AyrancI G, Bougacha S, Bienert T, Buehlmann D, Coletta L, Gallino D, Gass N, et al. Common functional networks in the mouse brain revealed by multi-centre resting-state fMRI analysis. Neuroimage. 2020;205.doi:10.1016/j.neuroimage.2019.116278

27. Casals JB, Pieri NCG, Feitosa MLT, Ercolin ACM, Roballo KCS, Barreto RSN, Bressan FF, Martins DS, Miglino MA, Ambrósio CE. The use of animal models for stroke research: a review. Comp. Med. 2011;61:305–13. doi:10.1109/TAES.1968.5408692

28. Bravo-Ferrer I, Cuartero MI, Zarruk JG, Pradillo JM, Hurtado O, Romera VG, Díaz-Alonso J, García-Segura JM, Guzmán M, Lizasoain I, et al. Cannabinoid type-2 receptor drives neurogenesis and improves functional outcome after stroke. Stroke. 2017;48:204–212. doi:10.1161/STROKEAHA.116.014793

29. Corbett D, Carmichael ST, Murphy TH, Jones TA, Schwab ME, Jolkkonen J, Clarkson AN, Dancause N, Wieloch T, Johansen-Berg H, et al. Enhancing the Alignment of the Preclinical and Clinical Stroke Recovery Research Pipeline: Consensus-Based Core Recommendations from the Stroke Recovery and Rehabilitation Roundtable Translational Working Group ∗. Neurorehabil. Neural Repair. 2017;31:699–707. doi:10.1177/1545968317724285

30. Manzanero S, Gelderblom M, Magnus T, Arumugam T V. Calorie restriction and stroke. Exp. Transl. Stroke Med. 2011;3:1–13. doi:10.1186/2040-7378-3-8

31. Lindenberg R, Renga V, Zhu LL, Betzler F, Alsop D, Schlaug G. Structural integrity of corticospinal motor fibers predicts motor impairment in chronic stroke. Neurology. 2010;74:280–287. doi:10.1212/WNL.0b013e3181ccc6d9

32. Feng W, Wang J, Chhatbar PY, Doughty C, Landsittel D, Lioutas V, Kautz SA, Schlaug G. Corticospinal tract lesion load: An imaging biomarker for stroke motor outcomes. Ann. Neurol. 2015;78:860–70. doi:10.1002/ana.24510

33. Frankel MR, Morgenstern LB, Kwiatkowski T, Lu M, Tilley BC, Broderick JP, Libman R, Levine SR, Brott T. Predicting prognosis after stroke: A placebo group analysis from the National Institute of Neurological Disorders and Stroke rt-PA stroke trial. Neurology. 2000;55:952–959. doi:10.1212/WNL.55.7.952

34. Zeynalov E, Jones SM, Elliott JP. Therapeutic time window for conivaptan treatment against stroke-evoked brain edema and blood-brain barrier disruption in mice. PLoS One. 2017;12:1–8. doi:10.1371/journal.pone.0183985

35. Liew SL, Lo BP, Donnelly MR, Zavaliangos-Petropulu A, Jeong JN, Barisano G, Hutton A, Simon JP, Juliano JM, Suri A, et al. A large, curated, open-source stroke neuroimaging dataset to improve lesion segmentation algorithms. Sci. Data. 2022;9:1–12. doi:10.1038/s41597-022-01401-7

36. Ito KL, Kim H, Liew SL. A comparison of automated lesion segmentation approaches for chronic stroke T1-weighted MRI data. Hum. Brain Mapp. 2019;40:4669–4685. doi:10.1002/hbm.24729

37. Verma K, Kumar S, Paydarfar D. Automatic Segmentation and Quantitative Assessment of Stroke Lesions on MR Images. Diagnostics. 2022;12. doi:10.3390/diagnostics12092055

38. Valverde JM, Shatillo A, De Feo R, Gröhn O, Sierra A, Tohka J. RatLesNetv2: A Fully Convolutional Network for Rodent Brain Lesion Segmentation. Front. Neurosci. 2020;14:1–11. doi:10.3389/fnins.2020.610239

39. Mulder IA, Khmelinskii A, Dzyubachyk O, de Jong S, Rieff N, Wermer MJH, Hoehn M, Lelieveldt BPF, van den Maagdenberg AMJM. Automated ischemic lesion segmentation in MRI mouse brain data after transient middle cerebral artery occlusion. Front. Neuroinform. 2017;11:1–15. doi:10.3389/fninf.2017.00003

40. An J, Wendt L, Wiese G, Herold T, Rzepka N, Mueller S, Koch SP, Hoffmann CJ, Harms C, Boehm-Sturm P. Deep learning-based automated lesion segmentation on mouse stroke magnetic resonance images. 2022;doi:https://www.biorxiv.org/content/10.1101/2022.08.09.503140v1

