## Supplemental Material including Supplemental Material & Methods, Supplemental Results, Figure S1-S14 and Supplemental Table S1-S9 for "Prediction of Stroke Outcome in Mice Based on Non-Invasive MRI and Behavioral Testing"

#### SUPPLEMENTAL METHODS

##### Experimental Design

In the *prediction cohort*, 148 mice received MCAO surgery (female: n = 40, male: n = 108). For comparison of functional outcome, 18 mice underwent sham surgery (female: n = 7, male: n = 11). In the *replication cohort*, 37 mice underwent MCAO surgery (female: n = 10, male: n = 27). For behavioral analysis, 12 mice received sham surgery (male: n = 12). A total of 15 different genotypes were used (Table S1). If activation of genotype was required (n = 50), either Tamoxifen (n = 43) or vehicle (n = 7) was used. A total of 19 mice were part of studies that included therapeutic treatment after the MCAO surgery of which 10 received an active agent and 9 received a placebo control (Table S1).

Mice were housed in groups of five in a 12 h day cycle and were food restricted but had free access to drinking water. Mice were weighed every day to control weight. They were familiarized with the pellets used in the behavioral task and with the test apparatus within the first week of handling by putting pellets in their cage. Eight days later, training sessions began (Figure 2A). Mice were initially food restricted for 20 h before training sessions; each session lasted 30 minutes. If weights stabilized around 85% of their free-feeding weights, food restriction was adjusted and increased to up to 22 h. After the training session, mice had individual access to standard lab chow diet to avoid food hoarding of the dominant mice. Mice were trained for fourteen days until average performance stabilized. We tested the mice for another seven consecutive days and the average performance during this period was used to normalize post-stroke pellet reaching. Performance was expressed as a percentage of average pre-stroke performance. Mice were randomized and allocated to either MCAO or sham surgery. 24 hours prior to the surgery mice had free access to food. After the surgery, mice had another two hours of access to food before the previously described cycle of food deprivation began. After MCAO lab chow diet pellets were homogenized using water to assist with feeding. 24 hours after the surgery MR imaging was performed. 48 hours after the surgery testing resumed for another twenty-one days at which point the animals were sacrificed.

##### Staircase Test of Skilled Reaching

The staircase test of skilled reaching in mice uses a testing box consisting of two stairs on the right and left side that were baited with up to two food pellets. The pellets consisted of sucrose (20 mg Dustless Precision Pellets®, Bio-Serv). Only one pellet was put onto the two highest stairs as mice would use their tongue to obtain these. In one study, number of pellets per side was 8 (n = 19), in one study it was 16 (n = 19) and in the remaining studies number of pellets per side was 14 (n = 147). For optimal comparability of motor-function, the individual performance was normalized to the pre-stroke performance. Therefore, average percentage of retrieved pellets was calculated for the period of baseline testing. Performance was then calculated as a percentage of the respective baseline performance.

**Effect of Study-Related Variables on Functional Outcome**

The effect of study-related concomitant variables (genotype, activation of genotype, therapeutic treatment, experiment number, sex, age at surgery, time to MCA occlusion and surgeon) on performance were investigated using separate analysis of variance linear mixed-effects models for *subacute* and *residual deficit* (response variable), respectively. For *subacute deficit*, a mixed-effects ANOVA revealed no significant effect of study-related variables (genotype: $F(11, 149) = 0.902$, $p = 0.540$; activation of genotype: $F(1, 149) = 0.849$, $p = 0.358$; therapeutic treatment: $F(4, 149) = 0.279$, $p = 0.891$; experiment number: $F(5, 149) = 0.955$, $p = 0.447$; sex: $F(1, 149) = 0.055$, $p = 0.814$; age: $F(1, 149) = 0.011$, $p = 0.917$; time to MCA occlusion: $F(1,149) = 0.919$, $p = 0.339$). The surgeon proved to not be an independent variable as each experiment was performed by one surgeon. Therefore, experiment number and surgeon were identical variables. For *residual deficit*, no significant effect of study-related variables was found (genotype: $F(11, 149) = 1.61$, $p = 0.102$; activation of genotype: $F(1, 149) = 3.503$, $p = 0.063$; therapeutic treatment: $F(4, 149) = 0.725$, $p = 0.576$; experiment number: $F(5, 149) = 1.60$, $p = 0.164$; sex: $F(1, 149) = 0.007$, $p = 0.931$; age: $F(1, 149) = 0.172$, $p = 0.679$; time to MCA occlusion: $F(1,149) = 0.227$, $p = 0.635$).

**MCAO Surgery**

Mice were anaesthetized with 1.5-3.5 % isoflurane and maintained with 1.0 % to 2.5 % in a mixture of 30 % $O_2$ and 70 % $N_2O$. During the procedure, body temperature was measured and maintained at 36 to 37.5 °C using a heating blanket and rectal probe. After disinfection of the skin, a midline neck incision was made and the common carotid artery (CCA) was carefully dissected from the surrounding nerves and ligated using 7.0 suture thread. The external carotid artery (ECA) was separated and ligated and the internal carotid artery (ICA) and the pterygopalatine (PA) arteries were clipped. A 180 µm diameter Doccol® filament was introduced into the common carotid artery, secured with a suture, the clip opened, and the filament advanced. The mice were allowed to recover in a heated cage at thermoneutral temperature (30-31 °C) for the duration of the MCAO. After 45 minutes, the mice were re-anaesthetized and the filament was withdrawn. The skin was sutured. As for the sham procedure, the filament was inserted to occlude the MCA and withdrawn immediately. For pain relief, Bupivacaine gel was topically applied in the vicinity of the wound. Standardized operation procedures were followed to minimize differences in total surgery duration in between the mice (doi: 10.1038/npre.2010.3492.2). Humane endpoints were reached as described in the section "2.2.5 Humane endpoint criteria" by Mei et al. 2019 (doi:10.14573/altex.1812231).

**MRI**

T2-weighted (T2w) images were acquired 24 h post stroke surgery on a 7 T MR scanner (Bruker, Ettlingen, Germany) and a 20 mm inner diameter transmit/receive volume coil (RAPID Biomedical, Rimpar, Germany). Anesthesia was achieved using 1-2 % isoflurane in a 70:30 nitrous oxide:oxygen mixture, and body temperature and respiration rate were monitored with MRI compatible equipment (Small Animal Instruments, Inc., Stony Brook, NY, USA). A 2D RARE sequence was used (repetition time/echo spacing/effective echo time=4.2 s/12 ms/36 ms, RARE factor=8, slice thickness 0.5 mm, FOV = $(25.6\ mm)^2$, image matrix=256x196 zerofilled to 256x256, 4 averages, acquisition time 6:43 min). The lesion was manually delineated by an experienced researcher using ANALYZE 5.0 software (AnalyzeDirect, Overland Park, KS, USA). MRI data were registered on the Allen Mouse Brain Atlas using the in-house developed MATLAB toolbox ANTx2 (https://github.com/ChariteExpMri/antx2). Incidence maps expressing percentage of animals with a lesion in a voxel were plotted for each group in atlas space and edema-corrected *lesion volume* was calculated as described previously.[21] Percentage of damage in a given region was calculated for each individual mouse using the delineated lesion masks and is referred to as *segmented MRI*. The rater of the MRI was blinded to both the type of intervention and the results of the staircase test.

**Statistics**

Behavioral data were analyzed using GraphPad Prism software, version 9.3.0. (La Jolla, CA, United States). A mixed-effects model followed by Šidák's post-hoc tests for multiple comparison was used to analyze the effect of side (right, paretic paw or left, non –paretic paw) and group (sham or MCAO) on performance. Alpha level was 0.05, mean difference and 95 % confidence intervals are reported. For the analysis of stroke volumes and deviation from group mean in percent when applying different occlusion times Kruskal-Wallis test was used, followed by Dunn's test to correct for multiple comparisons. For the comparison of lesion volumes and percentage of lesion within the caudoputamen in the *prediction* and *replication cohort*, Mann-Whitney test was used. For the comparison of *lesion volume* in different functional subgroups as well as comparison of different prediction models and their effect on the prediction error, either unpaired t-test or one-way ANOVA was used. ANOVA was followed by Tukey's test for multiple comparison. When the effects of the prediction model and the severity grades on the prediction error were analyzed, two-way-ANOVA followed by Tukey's test were performed.

**Data Availability**

Data is openly accessible on zenodo (https://doi.org/10.5281/zenodo.6534690). The repository contains all raw MRI T2w images, lesion masks and the registered atlases in NIFTI format, the input for machine learning algorithms (*lesion volume*, *segmented MRI*, behavioral scores), the trained classifiers and their output (predicted behavioral scores).

**Ethics approval**

All animal procedures were approved by and performed in concordance with local authorities (Landesamt für Gesundheit und Soziales) under license numbers G0197/12, G005/16, G0057/16, G0119/16, GG254/16, G0157/17, G0343/17.

**Division of Total Cohort into Training and Testing Group**

Since long-term outcome was the most important outcome measure in this study, we ensured a comparable distribution of the *residual deficit* in both groups. To achieve this, we first sorted the animals according to the *residual deficit*. Subsequently two animals from each consecutive triplet were randomly assigned to the training group and one to the test group, resulting in 98 animals in the training data set and 50 animals in the test data set. Analysis of behavioral data was done for the three groups (training cohort, *testing cohort*, sham animals) analogous to analysis of the entire cohort. Mean performance in the time period of *subacute deficit* was 58.56 % (51.81 to 65.30) on the non-paretic side and 36.50 % (29.29 to 43.71) on the paretic side of the *training cohort* and

67.56 % (58.24 to 76.89) on the non-paretic side and 39.73 % (30.60 to 48.85) on the paretic side of the *testing cohort* (Figure S12). Mixed-effects analysis revealed that the factors side and group had a significant impact on the performance (Side: $F(1, 164) = 30.29$, $p < 0.0001$; Group: $F(2, 164) = 7.09$, $p = 0.0011$). After correcting for multiple comparisons using Šidák's test, performance of the paretic paw was significantly lower in the *training* (t = 4.128, DF = 328, p = 0.0003) and *testing cohort* (t = 3.501, DF = 328, p = 0.0032) when compared to sham. There was no significant difference between *training* and *testing cohort* (t = 0.5506, DF = 328, p = 0.9947). There was also no significant difference between *training* (t = 2.293, DF = 328, p = 0.1275) or *testing cohort* (t = 1.167, DF = 328, p = 0.8133) on the non-paretic side when compared to sham.

Mean performance in the time period of *residual deficit* was 96.10 % (88.57 to 103.63) on the non-paretic side and 60.17 % (53.10 to 67.25) on the paretic side of the training cohort and 93.86 % (85.11 to 102.64) on the non-paretic side and 62.26 % (51.35 to 73.17) on the paretic side of the *testing cohort*. Mixed-effects analysis revealed that there was a statistically significant interaction between the effects side and group ($F(2, 165) = 4.875$, $p = 0.0088$). After correcting for multiple comparisons using Šidák's test, performance of the paretic paw was significantly lower in the *training* (t = 3.491, DF = 330, p = 0.0033) and *testing cohort* (t = 3.046, DF = 330, p = 0.0150) when compared to sham. There was no significant difference between *training* and *testing cohort* (t = 0.3449, DF = 330, p = 0.9996). There was also no significant difference between *training* (t = 0.4129, DF = 330, p = 0.9989) or *testing cohort* (t = 0.1542, DF = 330, p >0.9999) on the non-paretic side when compared to sham.

**Development of Prediction Models with Machine Learning**

For the development of prediction models with machine learning we divided the whole *prediction cohort* into a training (2/3 of the animals) and a testing cohort (1/3 of the animals). Using the training data, we tested several regression methods in the regression learner app in MATLAB (version 2021a) and chose random forest, which showed best performance on training data. We decided to train 50 independent models for each predictor additionally performing automated Bayesian hyperparameter optimization with 100 iterations for each single model.

**Estimation of Importance of Anatomical Regions Using Out-of-Bag Observations**

When training a random forest model, each tree is trained on a randomly selected subset of the training data, leaving out an average of one-third of the observations, which are then referred to as "out-of-bag" observations. These can be used to compute the importance of the variables, since they can serve as the test data set for the particular tree. First, the prediction error for each tree is computed using only the out-of-bag observations, then all values of one variable are permuted and the error is computed again for the out-of-bag observations, which now contain randomly permuted values for this particular variable. Subsequently, the difference between the errors before and after permutation is calculated. This procedure is repeated for each variable used to partition the nodes in the current tree. After each tree is processed, the mean and standard deviation of the error differences across all trees are calculated for each variable. Finally, the importance of each variable is calculated as the mean of the error differences divided by their standard deviation, which can also be considered as the signal-to-noise ratio. The out-of-bag predictor importance was estimated for each of the 50 random forest models trained on *segmented MRI* data. This was followed by calculation of the median value across all models and sorting of the regions by importance in descending order.

### SUPPLEMENTAL RESULTS

#### MRI Characterization and Lesion Topology in Prediction and Replication Cohort

Incidence maps were generated for the *prediction* and *replication cohort* (Figure S4A and S4B). Average *lesion volume* was 25.00mm$^3$ (21.46–28.54) in the *prediction cohort* and 30.43mm$^3$ (23.39-37.46) in the *replication cohort* (Figure S4C). Mean lesion volumes trended to be significantly larger in the *replication cohort* (actual difference: 13.57 mm$^3$, U=2186, p=0.058). Lesion volume in the caudoputamen accounted for 51.58% (46.42-56.75) in the *prediction cohort* but for only 38.15% (29.75-46.55) in the *replication cohort* (Figure S4D). Lesion volume in the caudoputamen accounted for a significantly larger proportion of the total stroke volume in the *prediction cohort* compared to the *replication cohort* (actual difference: 22.26%, U=1989, p=0.011). Incidence maps and stroke volumes were also generated and calculated for the functional subgroups of the *prediction cohort* (Figure S5A). Classified by degree of residual deficit, mean lesion volume was 14.03 mm$^3$ (9.82–18.25) in mild, 17.17 mm$^3$ (12.52–21.82) in moderate to mild, 26.29 mm$^3$ (16.86–35.73) in moderate, 33.62 mm$^3$ (24.04–43.20) in moderate to severe, and 47.23 mm$^3$ (36.79–57.68) in severe residual deficit mice (Figure S5B). One-Way ANOVA showed a significant difference in mean lesion volumes between functional subgroups (F(4,143)=14.14, p<0.0001; full report in Table S5).

#### Behavioral Analysis of the Replication Cohort

Behavioral and imaging data from two studies performed in 2015 and 2019 were pooled (Figure S2). After applying our exclusion criteria, 49 mice were included (MCAO: n = 37, sham: n = 12). Mean performance in the time period of *subacute deficit* was 70.98 % (60.93 to 81.02) on the non-paretic side and 44.21 % (35.78 to 52.65) on the paretic side (Figure S13A through S13C). In the sham animals, mean performance during the phase of *subacute deficit* was 101.15 % (93.17 to 109.13) on the non-paretic side and 103.96 % (97.89 to 110.02) on the paretic side. Mixed-effects analysis revealed that there was a statistically significant interaction between the effects side and group (F(1, 47) = 11.13, p = 0.0017). After correcting for multiple comparisons using Šidák's test, performance of the paretic paw was significantly lower in MCAO animals when compared to sham (least square mean difference = 59.74 p.p., t = 7.213, DF = 94, p < 0.0001). In the *replication cohort*, there was also a significant difference between groups on the non-paretic side (least square mean difference = 30.17 p.p., t = 3.643, DF = 94, p = 0.0009). Paired t-test was performed to test for a difference between paretic and non-paretic paw of the MCAO. Performance in the paretic paw was significantly lower than in the non-paretic paw (MD: 26.77 %, 16.85-36.68, t = 5.473, df = 36, p<0.0001) (Figure S7B). Mean performance in the period of *residual deficit* was 85.42 % (78.26 to 92.57) on the non-paretic side and 71.44 % (61.85 to 81.04) on the paretic side. In the sham animals, mean performance during the phase of *residual deficit* was 92.39 % (81.80 to 102.98) on the non-paretic side and 91.56 % (78.45 to 104.66) on the paretic side (Figure S13A, S13B and S13D). Mixed-effects analysis showed that a significant impact of the factor group on the degree of the *residual deficit* (F(1, 47) = 30.10, p < 0.0001). After correcting for multiple comparisons using Šidák's test, performance of the paretic paw was significantly lower in MCAO animals when compared to sham (least square mean difference = 38.85 p.p., t = 5.101, DF = 94, p < 0.0001). In the *replication cohort*, there was also a significant difference between groups on the non-paretic side (least square mean difference = 23.31 p.p., t = 3.061, DF = 94, p = 0.0057).

**Assessment of the Effects of General Well-Being of the Animals on Functional Outcome**

To test what effect the general well-being of MCAO animals had on the functional outcome, we performed correlation analyses using weight loss on day 2 and the Modified DeSimoni Neuroscore (from here on referred to as *neuroscore*) for the assessment of general deficits and compared this to the *lesion volume*. First, using Pearson's correlation for normally distributed data and Spearman's correlation for non-parametric data was used to test if weight loss on day 2, *neuroscore* and the *lesion volume* significantly correlated to the degree of the *subacute deficit*. The correlation was statistically significant for all three factors (weight loss: r(183) = 0.253, p = 0.0005; $R^2$ = 0.064; *neuroscore*: r(172) = -0.327, p<0.0001, $R^2$ = 0.107; *lesion volume*: rho(183) = -0.618, p < 0.0001) (Figure S8). Notably, while significant, both the *neuroscore* and the weight loss on day 2 did not seem to explain much of the variation of the *subacute deficit* as indicated by small r and $R^2$. This became even more clear when comparing these parameters with the *lesion volume*, where rho was much higher and seemed to explain more of the variation observed in the *subacute deficit* data.

Next, we repeated these analyses for the *residual deficit*, now also including the *subacute deficit* as an independent variable. Again, the correlation was statistically significant for all four factors (weight loss: r(183) = 0.236, p = 0.0012, $R^2$ = 0.056; *neuroscore*: r(172) = -0.241, p = 0.0013, $R^2$ = 0.058,; *lesion volume*: rho(183) = -0.466, p < 0.0001; subacute deficit: r(183) = 0.736, p < 0.0001, $R^2$ = 0.543) (Figure S9). Again, with r and $R^2$ being small for both, w*eight loss* and *neuroscore*, these parameters did not seem to explain much of the variation observed in the residual deficit data.

In conclusion, correlation analyses indicated that the general well-being of the animals explained only a small fraction of the variation seen in the *subacute* and *residual deficit* data.

**General well-being parameters as predictors of subacute and residual deficit**

Finally, we generated two new prediction models that were based on the *weight loss* on day 2 and the *neuroscore*. We compared these two new models to our previously described predictors, namely the *lesion volume*, the *segmented MRI*, and the *subacute deficit*. Notably, data on the *neuroscore* was only available for 94 % of our cohort (data available: total cohort: n = 174; training cohort: n = 99; *testing cohort*: n = 49; validation cohort: n = 37). In the *testing cohort, PE* was 23.33pp (23.09-23.57, $Q_1$: 13.95, $Q_3$: 36.58, IQR: 22.63) for the *neuroscore* and 28.26pp (28.06-28.46, $Q_1$: 10.66, $Q_3$: 42.03, IQR: 31.37) for the *weight loss* (Figure S10A and S10B) when used for the prediction of the *subacute deficit*. One-way ANOVA showed predictors differed significantly (F(3,196) = 3306, p<0.0001). After multiple comparisons were corrected using Tukey's test, the *PEs* of both the *weight loss* and the *neuroscore* were significantly higher than the *PEs* of the *segmented MRI* (MD = 12.04pp, 11.60-12.47, p<0.0001 and MD = 7.11pp, 6.67-7.54, p<0.0001, respectively) and the *lesion volume* (MD = 14.96pp, 14.53-15.40, p<0.0001 and MD = 10.03pp, 9.60-10.47, p<0.0001, respectively). The *neuroscore* outperformed the prediction accuracy of the *weight loss* (MD = 4.93pp, 4.50-5.35, p<0.0001). In the *replication cohort*, *PE* was 20.65pp (20.47-20.82, $Q_1$: 13.30, $Q_3$: 28.18, IQR: 14.88) for the *neuroscore* and 21.98pp (21.79-22.16, $Q_1$: 10.33, $Q_3$: 35.64, IQR: 25.31) for the *weight loss* (Figure S10C and S10D) when used for the prediction of the *subacute deficit*. A One-way ANOVA showed predictors differed significantly (F(3,196) = 130.3, p<0.0001). After multiple comparisons were corrected using Tukey's test, again, the *PEs* of both the *weight loss* and the *neuroscore* were significantly higher than *PEs* of the *segmented MRI* (MD = 1.97pp, 1.52-2.43, p<0.0001 and MD = 0.64pp, 0.19-1.10, p = 0.0019, respectively) and the *lesion volume* (MD = 1.21pp, 0.76-1.67, p<0.0001 and MD = 2.55pp, 2.09-3.00, p<0.0001). The *neuroscore* outperformed the prediction accuracy of the *weight loss* (MD = 1.33pp, 0.88-1.79, p<0.0001).

When used for the prediction of the *residual deficit* in the *testing cohort*, *PE* was 29.25pp (29.15-29.34, $Q_1$: 13.25, $Q_3$: 45.32, IQR: 32.07) for the *neuroscore* and 30.14pp (29.95-30.34, $Q_1$: 10.58, $Q_3$: 45.26, IQR: 34.67) for the *weight loss* (Figure S11A and S11B). One-way ANOVA showed predictors differed significantly (F(4,245) = 2285, p<0.0001). After multiple comparisons were corrected using Tukey's test, the *PEs* of both the *weight loss* and the *neuroscore* were significantly higher than *PEs* of the *segmented MRI* (MD = 7.47pp, 7.09-7.85, p<0.0001 and MD = 6.57pp, 6.19-6.95, p<0.0001, respectively) and the *lesion volume* (MD = 4.19pp, 3.81-4.57, p<0.0001 and MD = 3.29pp, 2.91-3.67, p<0.0001). The *subacute* deficit clearly outperformed both, the *weight loss* (MD = 11.25pp, 10.87-11.63, p<0.0001) and the *neuroscore* (MD = 10.36pp, 9.98-10.74, p<0.0001) as predictor of the *residual deficit*. Again, the *neuroscore* outperformed the prediction accuracy of the *weight loss* (MD = 0.90pp, 0.52-1.28, p<0.0001). In the *replication cohort*, *PE* was 24.37pp (23.89-24.86, $Q_1$:17.52, $Q_3$: 33.37, IQR: 15.86) for the *neuroscore* and 24.63pp (24.38-24.88, $Q_1$: 16.59, $Q_3$: 36.30, IQR: 19.71) for the *weight loss* (Figure S11C and S11D) when used for the prediction of the *residual deficit*. One-way ANOVA showed predictors differed significantly (F(4,245) = 494.4, p<0.0001). After multiple comparisons were corrected using Tukey's test, the *PEs* of both the *weight loss* and the *neuroscore* were significantly higher than the *PE* of the *subacute deficit* (MD = 13.87pp, 12.83-14.91, p<0.0001 and MD = 13.61pp, 12.58-14.65, p<0.0001, respectively). *PEs* of both the *weight loss* and the *neuroscore* were significantly higher than the *PEs* of the *segmented MRI* (MD = 1.68pp, 0.64-2.71, p<0.0001 and MD = 1.42pp, 0.38-2.45, p = 0.0021, respectively), but not significantly different from the PE of the *lesion volume* (MD = 0.70pp, 0.34-1.74, p = 0.347, and MD = 0.44pp, 0.60-1.48, p = 0.773).
In conclusion, for the prediction of the *subacute deficit*, in both the *testing cohort* and the *replication cohort*, *PEs* of the parameters we used as proxies for general well-being were significantly higher than our imaging-based predictors. Additionally, the *subacute deficit* and the segmented MRI outperformed the *weight loss* and the *neuroscore* as predictors of the *residual deficit*, underscoring the meaningfulness of analyzing the early functional outcome in mice.

**Prediction Accuracy Depends on Severity Grade**
Prediction errors were examined to better understand the relationship between severity grade and model accuracy (Figure S14). For the assessment of the prediction accuracy of the *subacute deficit* from *lesion volume* or from *segmented MRI*, severity grades were based on the subacute deficit. Two-Way ANOVA was performed to test whether severity grades and prediction model had an effect on the *prediction error*. Two-Way-ANOVA showed a statistically significant interaction between prediction model and severity grade (F(4, 490) = 682.3, p<0.0001). Simple main effect analysis revealed that severity grade (F(4, 490) = 2822, p<0.0001) and model (F(1, 490) = 47.49, p < 0.0001) had a statistically significant effect on the prediction error. For a full statistical report, see Table S6. For the assessment of the prediction accuracy of *residual deficit*, severity grades were based on the residual deficit. Besides the two imaging based parameters, the degree of *subacute deficit* was included in the analysis. Two-Way-ANOVA showed a statistically significant interaction between predictor and severity grade (F(8, 735) = 214.5, p<0.0001). Simple main effect analysis revealed that severity grade (F(4, 735) = 1819, p<0.0001) and predictor (F(2, 735) = 1241, p < 0.0001) had a statistically significant effect on the prediction error. For a full statistical report, see Table S7.
We repeated these analyses using the same models on an independent *replication cohort*. Two-Way ANOVA was performed to test whether severity grades and prediction model had an effect on the *prediction error*. Two-Way-ANOVA showed a statistically significant interaction between prediction model and severity grade (F(4, 490) = 1231, p < 0.0001). Simple main effect analysis

revealed that severity grade ($F(4, 490) = 3227$, $p<0.0001$) and model ($F(1, 490) = 152.3$, $p<0.0001$) had a statistically significant effect on the prediction error. For a full statistical report, see Table S8. For the assessment of the prediction accuracy of *residual deficit*, severity grades were based on the residual deficit. Besides the two imaging based parameters, the degree of *subacute deficit* was included in the analysis. Two-Way-ANOVA showed a statistically significant interaction between predictor and severity grade ($F(8, 735) = 380.2$, $p<0.0001$). Simple main effect analysis revealed that severity grade ($F(4, 735) = 1678$, $p < 0.0001$) and predictor ($F(2, 735) = 614.5$, $p < 0.0001$) had a statistically significant effect on the prediction error. For a full statistical report, see Table S9.

**Supplemental Figures and Tables**

**Prediction of Stroke Outcome in Mice Based on Non-Invasive MRI and Behavioral Testing**

Felix Knab, Stefan Paul Koch, Sebastian Major, Tracy D. Farr, Susanne Mueller, Philipp Euskirchen, Moritz Eggers, Melanie T.C. Kuffner, Josefine Walter, Daniel Berchtold, Samuel Knauss, Jens P. Dreier, Andreas Meisel, Matthias Endres, Ulrich Dirnagl, Nikolaus Wenger, Christian J. Hoffmann, Philipp Boehm-Sturm, Christoph Harms

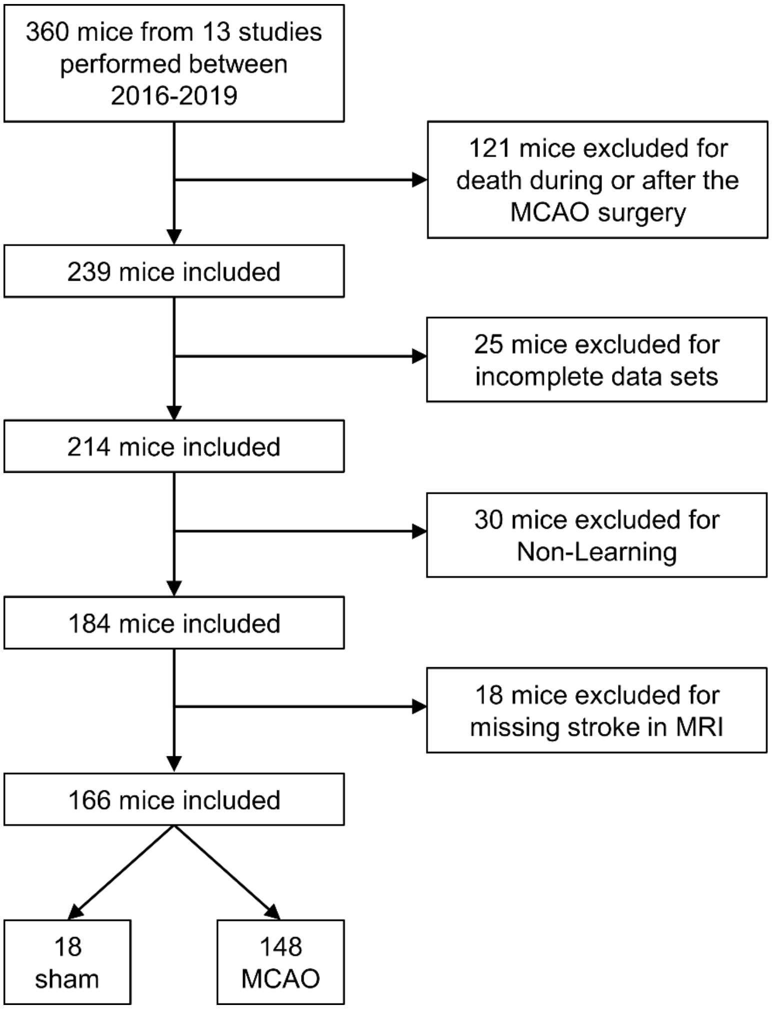


**Figure S1 An overview of the inclusion/exclusion of mice after combining data from 13 preclinical studies.** A total of four exclusion criteria were applied.

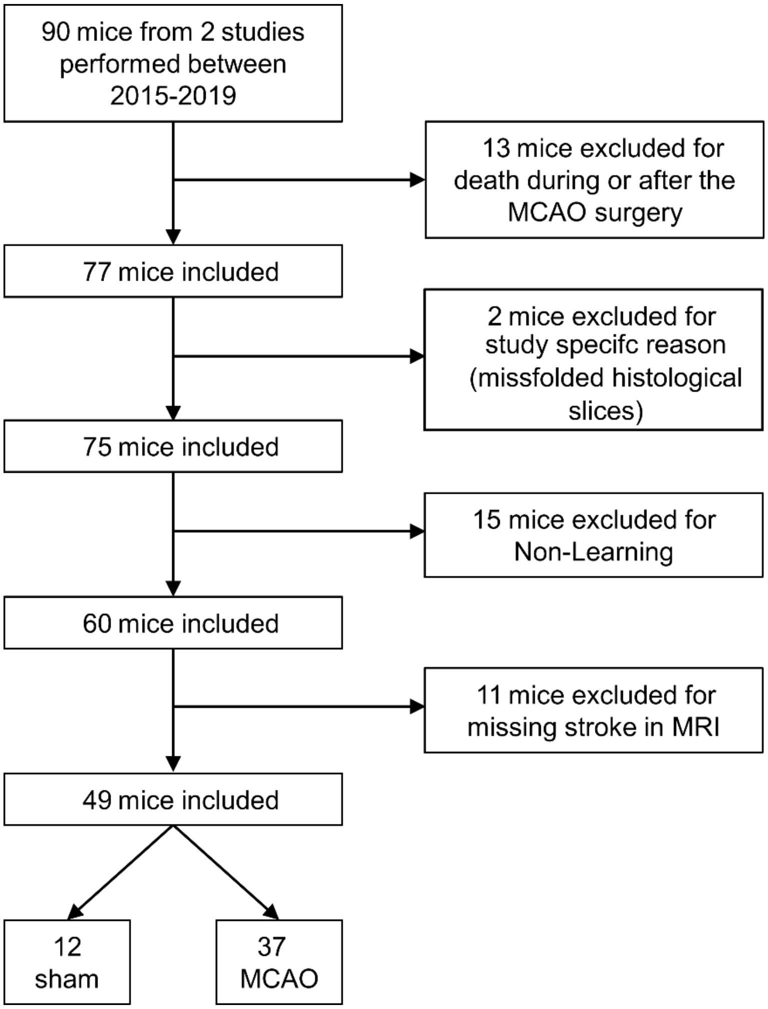


**Figure S2. Overview of the total cohort and exclusion process for the replication cohort**
Three exclusion criteria that had previously been defined and applied have been used for building the replication cohort. In addition, two mice were excluded from one study due to a study specific exclusion criteria that had not played a role in our previous studies. In the end, a total of 49 mice were included.

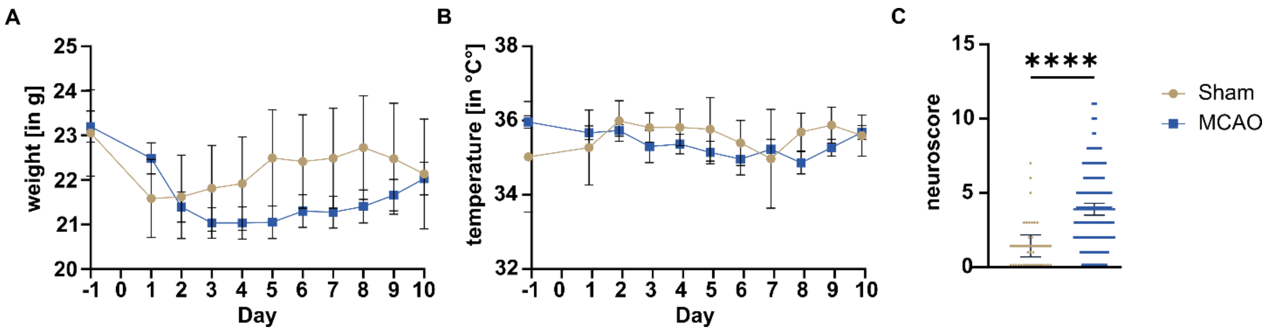


**Figure S3. Weight and Temperature of MCAO and sham animals after surgery.**
**A**, Weights and **B**, temperature of sham and MCAO animals within the first 10 days of surgery. Error bars indicate the 95% confidence interval. Days -1 to 8 (sham: n = 30; MCAO: n = 185) and days 9 to 10 (sham: n = 25-26; MCAO: n = 182-183). **C**, results of the modified DeSimoni neuroscore for the assessment of general deficits on day 2. Error bars indicate a 95% confidence interval (sham: n = 30; MCAO: n = 174).

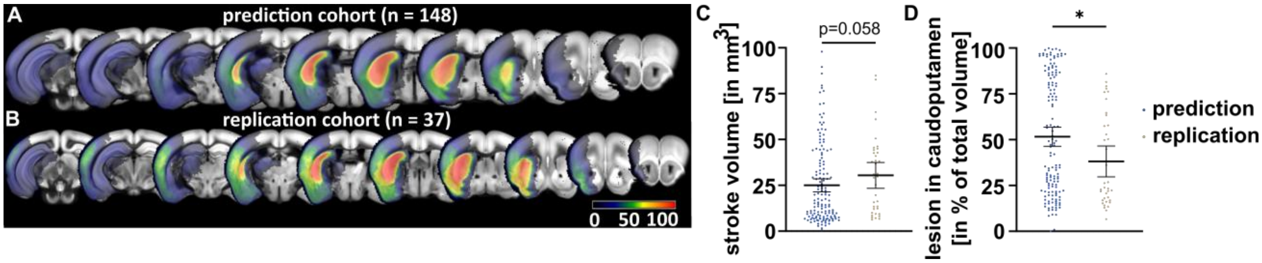


**Figure S4 MRI characterization and lesion topology.**

**A**, Lesion incidence maps of mice in the prediction and **B**, replication cohort. **C**, While stroke volumes were only trending towards being significantly larger in the replication cohort, **D** percentage of the total stroke volume that was within the caudoputamen was significantly smaller in the replication cohort. This indicates, that in the prediction cohort, cortical involvement was less frequent.

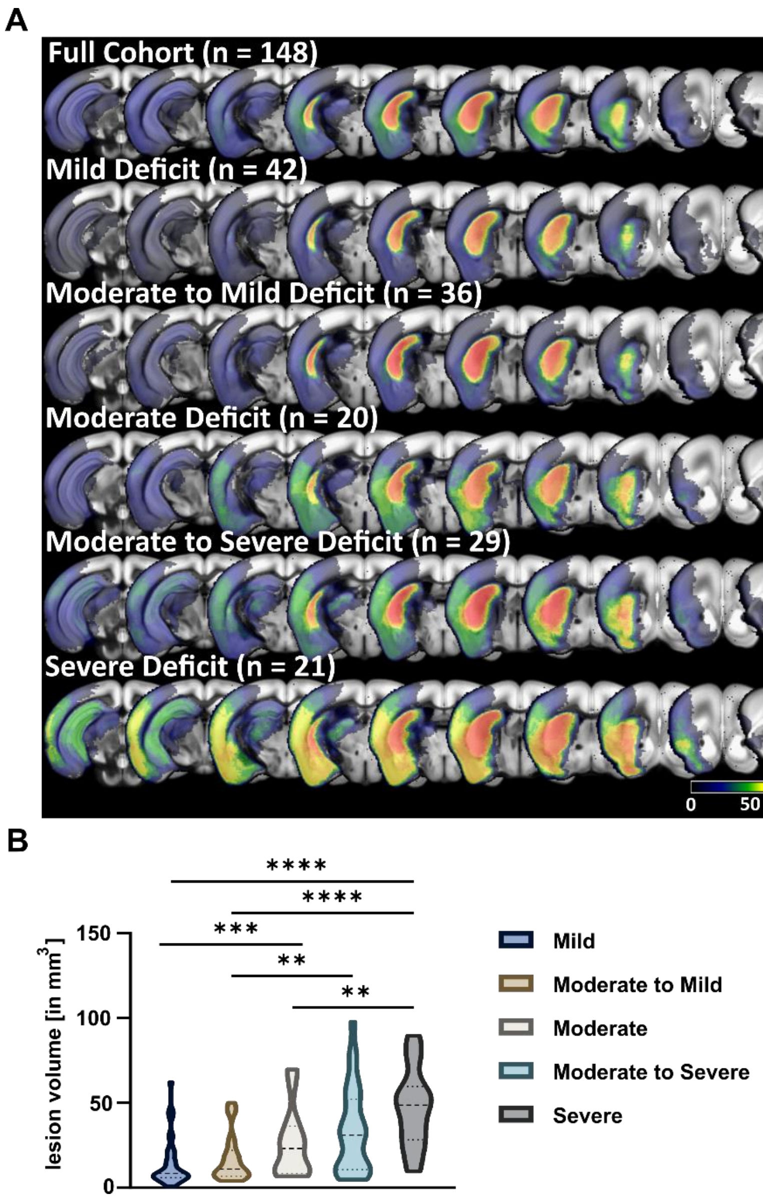


**Figure S5. MRI characterization of lesion volume and location.**

**A**, Coronal incidence maps of stroke lesions in the total cohort and the five different functional subgroups. Colours indicate percentage of mice with a lesion in a given region. Severity grades are based on degree of *residual deficit*. **B**, *lesion volume* by severity grade (based on degree of *residual deficit*). Great variability within functional subgroups could be observed and there was no significant difference when comparing *lesion volume* of consecutive severity grades, i.e. of mice with mild or moderate to mild *residual deficits*.

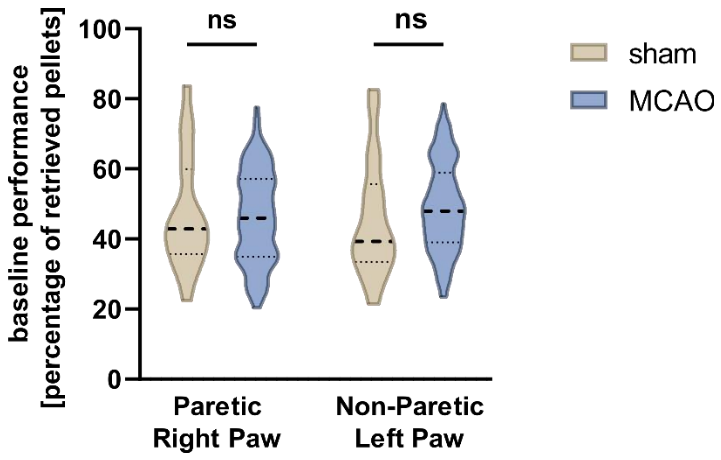


**Figure S6 Comparison of baseline performance between sham and MCAO reveals no significant difference.** Percentage of retrieved pellets during baseline testing (days -7 to -1). Mixed-effects model followed by Šidák's post-hoc tests for multiple comparison were used to test effect of group and side. No statistically significant differences were found. Dashed lines indicate medians, dotted line indicate quartiles.

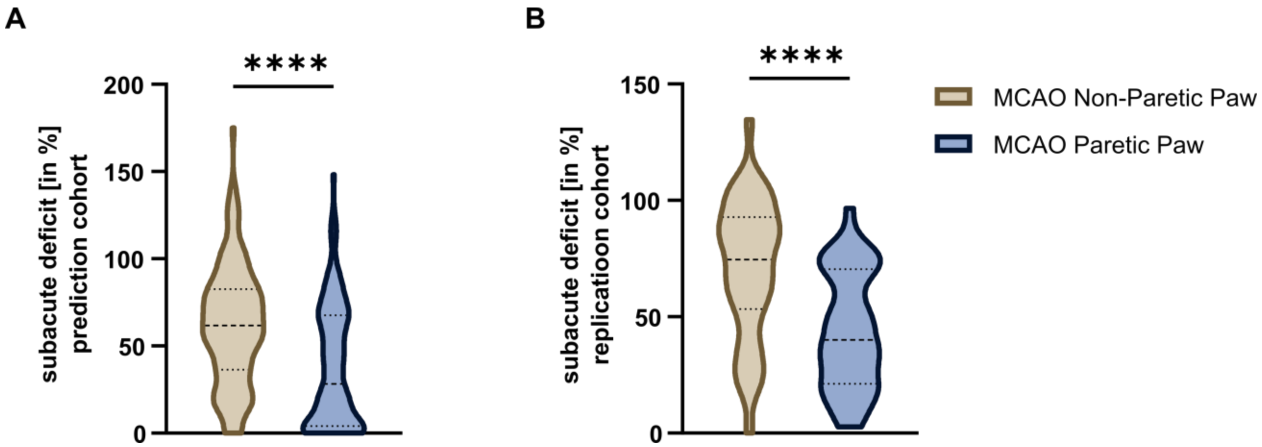


**Figure S7 Comparison of the subacute deficit of non-paretic and paretic paw in the MCAO mice. A**, Data on the subacute deficit are pooled from the training and testing cohorts. Paired t-test was performed to test for side-specific motor-functional deficits. Dashed lines indicate medians, dotted line indicate quartiles. **B**, Subacute deficit of the replication cohort. Again, a side specific decrease of the paretic paw was observed.

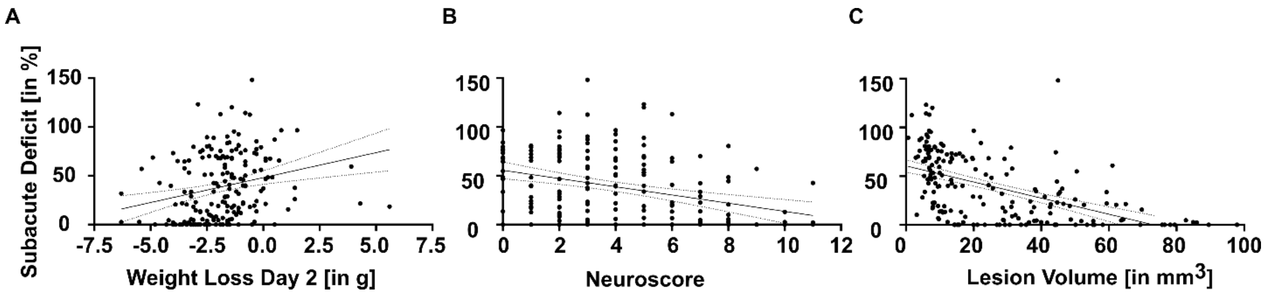


**Figure S8. Assessment of the Effects of General Well-Being of Animals on Subacute Deficit**
**A**, Simple linear regression analysis was performed for the weight loss on day 2 **B**, the neuroscore and **C**, the lesion volume to test whether these parameters correlate to the degree of the subacute deficit. Dotted line indicates 95% confidence band of the best-fit line.

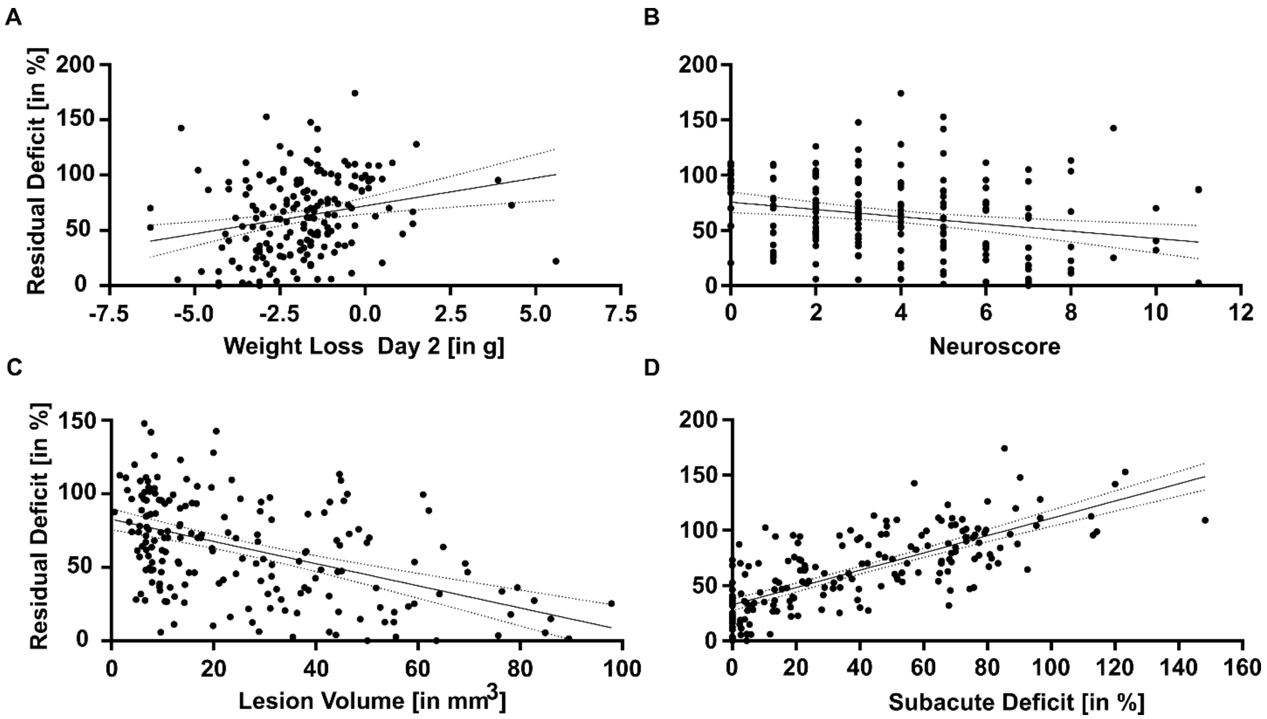


**Figure S9. Assessment of the Effects of General Well-Being of Animals on Residual Deficit** **A**, Simple linear regression analysis was performed for the weight loss on day 2 **B**, the neuroscore **C**, the lesion volume and **D**, the subacute deficit to test whether these parameters correlate to the degree of the subacute deficit. Dotted line indicates 95% confidence band of the best-fit line.

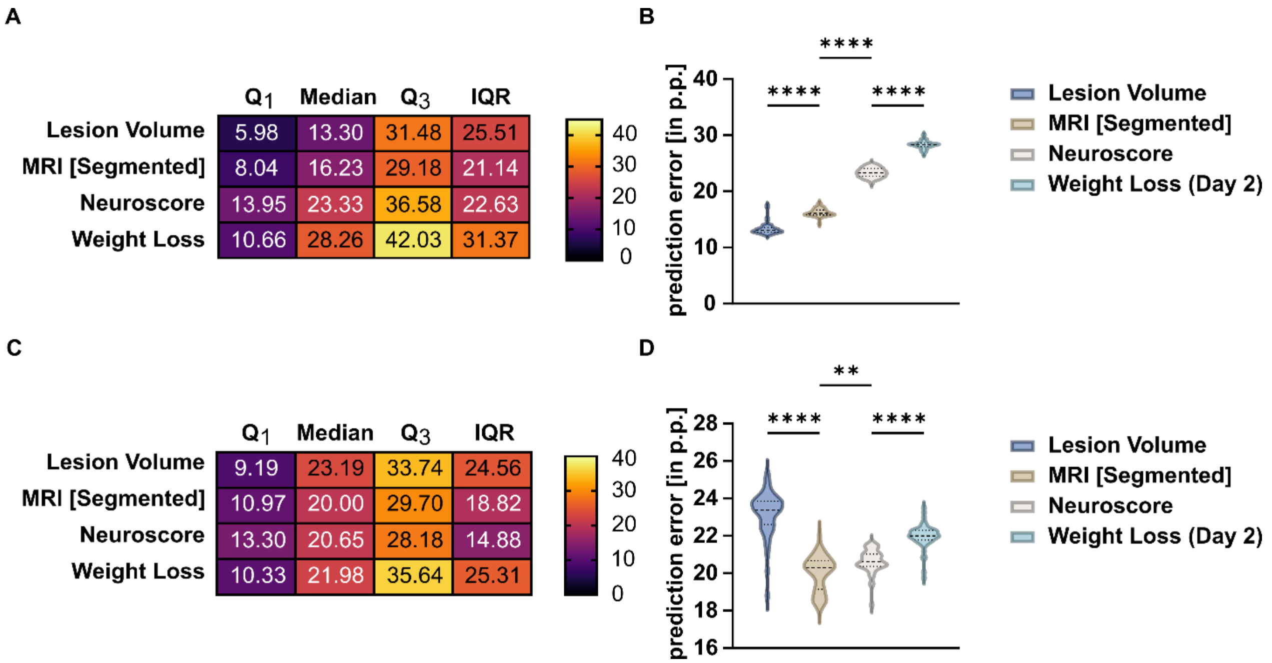


**Figure S10. General well-being as predictor of subacute deficit**

**A**, and **B**, *Prediction error* of the four models for the prediction of the *subacute deficit* in the testing cohort **C**, and **D** *Prediction error* of the four models for the prediction of the *subacute deficit* in the *replication cohort*.

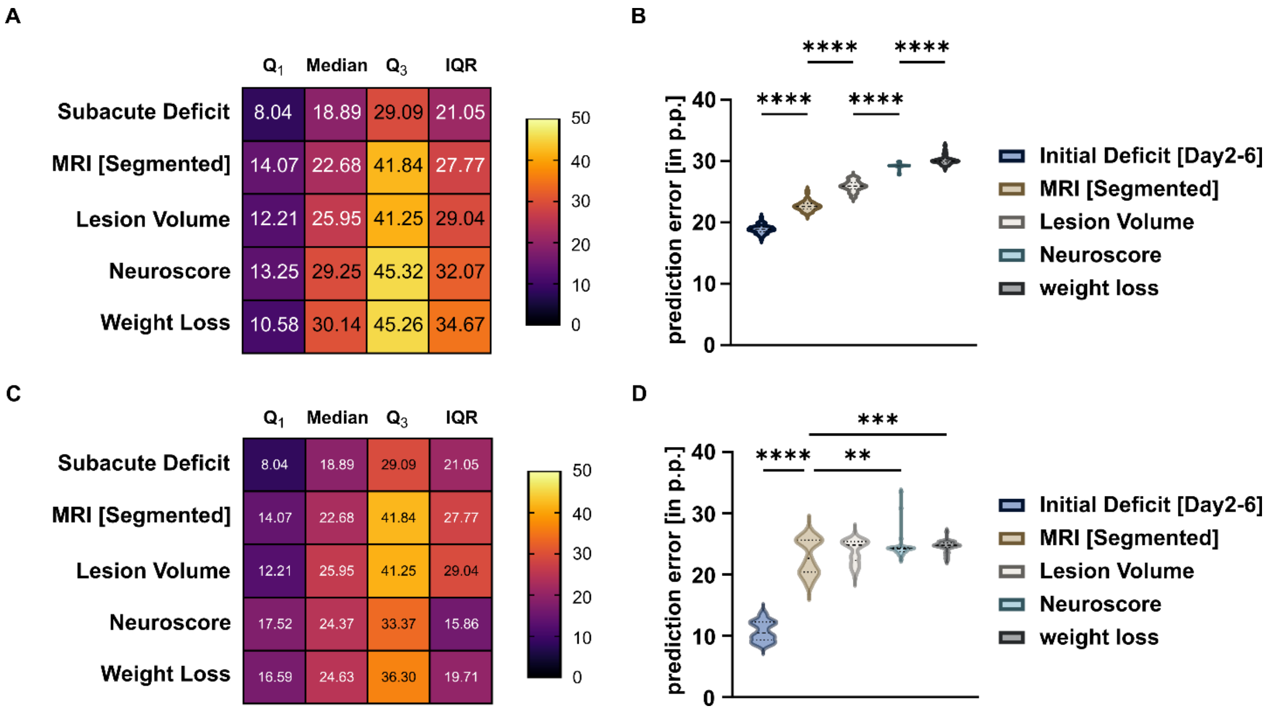


**Figure S11. General well-being as predictor of residual deficit**

**A**, and **B**, *Prediction error* of the four models for the prediction of the *residual deficit* in the testing cohort **C**, and **D** *Prediction error* of the four models for the prediction of the *residual deficit* in the *replication cohort*.

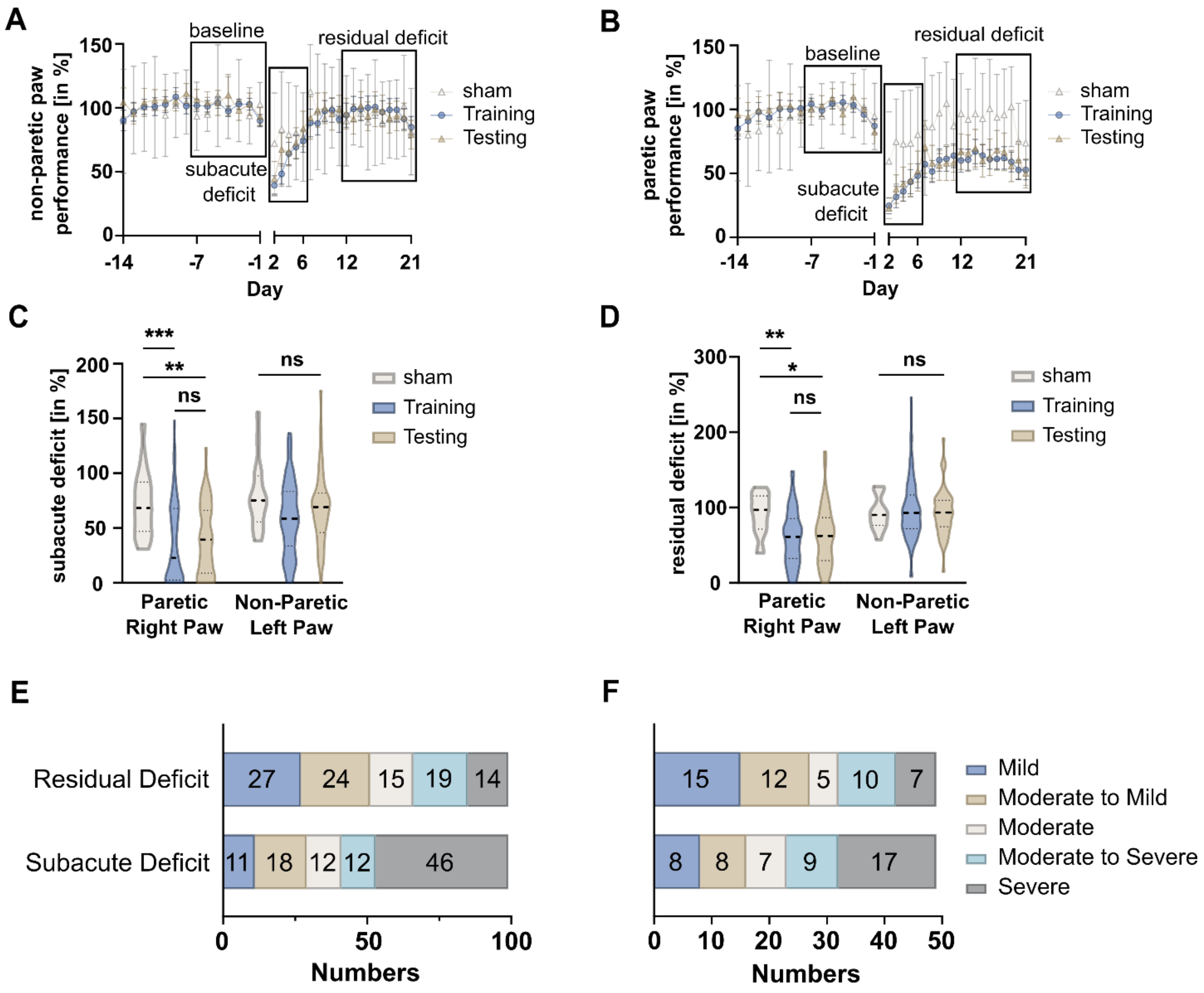


**Figure S12. Analysis of the behavioral data of the training and testing cohort**
**A**, Non-paretic left and **B**, paretic right paw performance over time, bars depict 95 % confidence interval. Performance on days -7 to -1 were averaged and taken as a baseline (first black rectangle). Data from days 2 to 6 were summarized and referred to as *subacute deficit* (second rectangle) while data from days 12 to 21 are referred to as *residual deficit* (third rectangle). **C**, *Subacute* and **D**, *residual deficit* of the non-paretic and paretic paw in sham (white plot), *training cohort* (blue plot) and *testing cohort* (brown plot). Dashed lines indicate medians, dotted line indicate quartiles. Mixed-effects model followed by Šidák's post-hoc tests for multiple comparison were used to test effect of group and side. **E**, Overview over sizes of functional subgroups in the *training cohort* and **F**, *testing cohort*. Groups are either based on the degree of *subacute* or *residual deficit*.

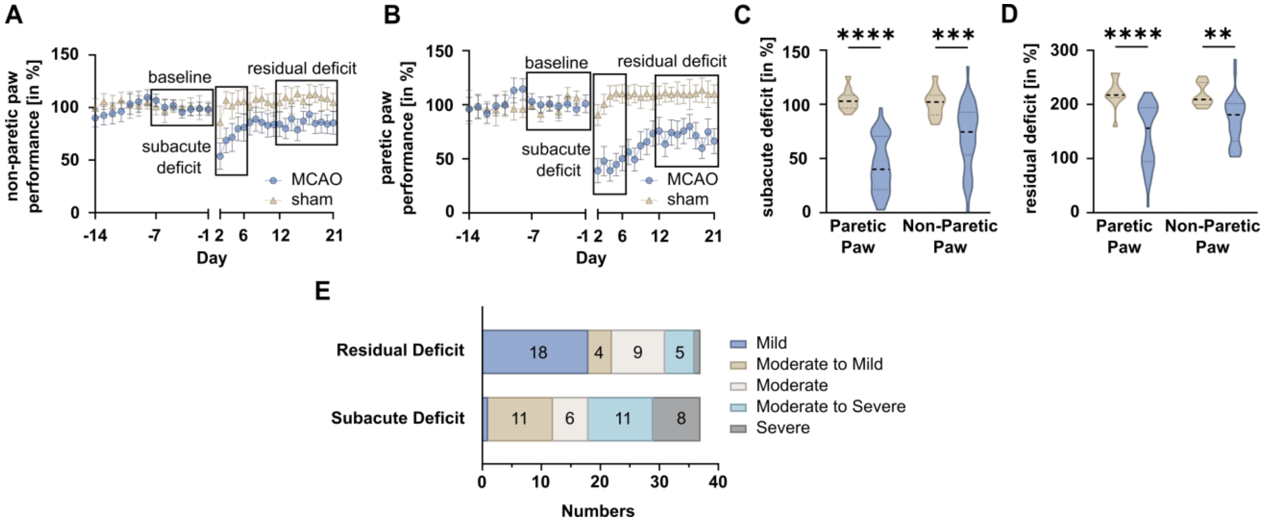


**Figure S13. Assessment of motor-functional deficit and stroke composition**
**A**, Non-paretic left and **B**, paretic right paw performance over time, bars depict 95 % confidence interval. Performance on days -7 to -1 were averaged and taken as a baseline (first black rectangle). Data from days 2 to 6 were summarized and referred to as *subacute deficit* (second rectangle) while data from days 12 to 21 are referred to as *residual deficit* (third rectangle). **C**, *Subacute* and **D**, *residual deficit* of the non-paretic and paretic paw in sham (brown plot) and MCAO (blue plot) animals. Dashed lines indicate medians, dotted line indicate quartiles. Mixed-effects model followed by Šidák's post-hoc tests for multiple comparison were used to test effect of group and side. **E**, Overview of sizes of functional subgroups. Groups are either based on the degree of *subacute* or *residual deficit*. In our replication cohort, only one mouse showed a mild *subacute* or severe *residual deficit* (not depicted). **F**, In the *replication cohort*, lesions in the caudoputamen accounted for a significantly smaller fraction when compared to the *testing cohort*.

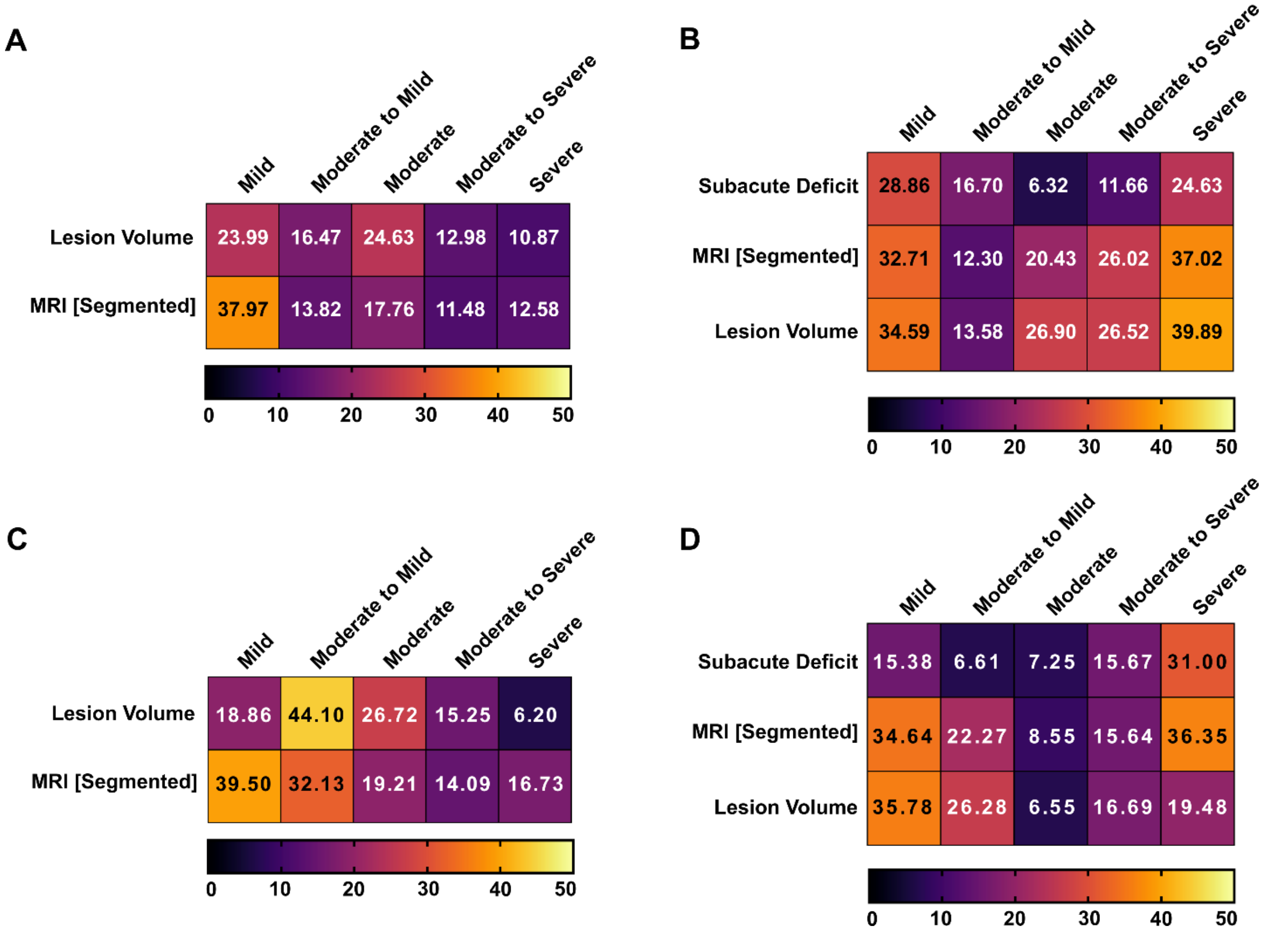


**Figure S14 Comparison of severity grade dependent prediction accuracy.**
**A**, *Prediction error* of the two predictors of the *subacute deficit* over the severity grades. Independent of the predictor, errors were higher in mice with mild compared to moderate to severe *subacute deficits*, indicating that the prediction models work especially well in those two crucial subgroups **B**, Prediction accuracy of the three predictors of the *residual deficit* over the severity grades. Again, a strong dependency of *prediction error* on the degree of deficit could be observed. **C**, *Prediction error* of the two predictors of the *subacute deficit* over the severity grades in the *replication cohort*. Again, *prediction errors* were smallest in mice with severe or moderate to severe subacute deficits. **D**, Prediction accuracy of the three predictors of the *residual deficit* over the severity grades in the *replication cohort*.

| Experiment ID | sex | age at MCAO (in weeks) | Genotype | Genotype Activation | Therapeutic Treatment | MCAO vs Sham | cohort |
|---|---|---|---|---|---|---|---|
| 2015-4 | m | 12 | C57Bl6_N | none | none | MCAO | replication cohort |
| 2015-4 | m | 12 | C57Bl6_N | none | none | MCAO | replication cohort |
| 2015-4 | m | 12 | C57Bl6_N | none | none | MCAO | replication cohort |
| 2015-4 | m | 12 | C57Bl6_N | none | none | MCAO | replication cohort |
| 2015-4 | m | 12 | C57Bl6_N | none | none | MCAO | replication cohort |
| 2015-4 | m | 12 | C57Bl6_N | none | none | MCAO | replication cohort |
| 2015-4 | m | 12 | C57Bl6_N | none | none | MCAO | replication cohort |
| 2015-4 | m | 12 | C57Bl6_N | none | none | sham | replication cohort |
| 2015-4 | m | 12 | C57Bl6_N | none | none | sham | replication cohort |
| 2015-4 | m | 12 | C57Bl6_N | none | none | sham | replication cohort |
| 2015-4 | m | 12 | C57Bl6_N | none | none | sham | replication cohort |
| 2015-4 | m | 12 | C57Bl6_N | none | none | sham | replication cohort |
| 2015-4 | m | 12 | C57Bl6_N | none | none | sham | replication cohort |
| 2015-4 | m | 12 | C57Bl6_N | none | none | sham | replication cohort |
| 2015-4 | m | 12 | C57Bl6_N | none | none | sham | replication cohort |
| 2015-4 | m | 12 | C57Bl6_N | none | none | sham | replication cohort |
| 2015-4 | m | 12 | C57Bl6_N | none | none | sham | replication cohort |
| 2015-4 | m | 12 | C57Bl6_N | none | none | sham | replication cohort |
| 2015-4 | m | 12 | C57Bl6_N | none | none | sham | replication cohort |
| 2015-8 | m | 13 | C57Bl6_N | none | none | MCAO | replication cohort |
| 2015-8 | m | 13 | C57Bl6_N | none | none | MCAO | replication cohort |
| 2015-8 | m | 13 | C57Bl6_N | none | none | MCAO | replication cohort |
| 2015-8 | m | 13 | C57Bl6_N | none | none | MCAO | replication cohort |

Continued Table S1

| | | | | | | | |
|---|---|---|---|---|---|---|---|
| 2015-8 | m | 13 | C57Bl6_N | none | none | MCAO | replication cohort |
| 2015-8 | m | 13 | C57Bl6_N | none | none | MCAO | replication cohort |
| 2015-8 | m | 13 | C57Bl6_N | none | none | MCAO | replication cohort |
| 2015-8 | m | 13 | C57Bl6_N | none | none | MCAO | replication cohort |
| 2015-8 | m | 13 | C57Bl6_N | none | none | MCAO | replication cohort |
| 2015-8 | m | 13 | C57Bl6_N | none | none | MCAO | replication cohort |
| 2015-8 | m | 13 | C57Bl6_N | none | none | MCAO | replication cohort |
| 2016-11 | m | 13 | B6.129 Mertktm1 Grl/J mutant | none | none | MCAO | Training |
| 2016-11 | m | 13 | B6.129 Mertktm1 Grl/J wildtype | none | none | MCAO | Testing |
| 2016-11 | m | 13 | B6.129 Mertktm1 Grl/J mutant | none | none | MCAO | Training |
| 2016-11 | m | 13 | Mfge8 mutant | none | none | MCAO | Training |
| 2016-11 | m | 14 | Mfge8 mutant | none | none | MCAO | Training |
| 2016-11 | m | 14 | Mfge8 mutant | none | none | MCAO | Testing |
| 2016-11 | m | 14 | B6 Mfge8 GtByg wildtype | none | none | MCAO | Training |
| 2016-11 | m | 14 | B6.129 Mertktm1 Grl/J wildtype | none | none | MCAO | Training |
| 2016-11 | m | 15 | B6.129 Mertktm1 Grl/J wildtype | none | none | MCAO | Training |
| 2016-11 | m | 15 | B6 Mfge8 GtByg mutant | none | none | MCAO | Training |
| 2016-11 | m | 16 | Mfge8 mutant | none | none | MCAO | Training |
| 2016-11 | m | 16 | Mfge8 mutant | none | none | MCAO | Training |
| 2016-11 | m | 17 | B6 Mfge8 GtByg mutant | none | none | MCAO | Training |
| 2016-11 | m | 17 | B6 Mfge8 GtByg mutant | none | none | MCAO | Training |
| 2016-11 | m | 17 | B6 Mfge8 GtByg mutant | none | none | MCAO | Training |
| 2016-11 | m | 17 | B6.129 Mertktm1 Grl/J mutant | none | none | MCAO | Training |

Continued Table S1

| | | | | | | | |
|---|---|---|---|---|---|---|---|
| 2016-11 | m | 18 | B6 Mfge8 GtByg wildtype | none | none | MCAO | Training |
| 2016-2 | f | 17 | VecdhxFlexIL6 Cre/tg | Tamoxifen | none | sham | sham |
| 2016-2 | f | 19 | VecdhxFlexIL6 Cre/tg | Tamoxifen | none | MCAO | Training |
| 2016-2 | m | 20 | VecdhxFlexIL6 Cre/tg | Tamoxifen | none | MCAO | Testing |
| 2016-2 | m | 21 | VecdhxFlexIL6 Cre/tg | Tamoxifen | none | sham | sham |
| 2016-2 | f | 21 | VecdhxFlexIL6 Cre/tg | Tamoxifen | none | sham | sham |
| 2016-2 | f | 23 | VecdhxFlexIL6 Cre/tg | Tamoxifen | none | MCAO | Testing |
| 2016-4 | m | 11 | B6.129 Mertktm1 Grl/J mutant | none | none | sham | sham |
| 2016-4 | m | 11 | B6.129 Mertktm1 Grl/J mutant | none | none | MCAO | Training |
| 2016-4 | m | 11 | B6.129 Mertktm1 Grl/J mutant | none | none | MCAO | Testing |
| 2016-4 | m | 11 | B6.129 Mertktm1 Grl/J wildtype | none | none | sham | sham |
| 2016-4 | m | 11 | B6.129 Mertktm1 Grl/J wildtype | none | none | sham | sham |
| 2016-9 | m | 15 | Mertk wildtype | none | none | MCAO | Training |
| 2016-9 | m | 16 | B6 Mfge8 GtByg mutant | none | none | MCAO | Training |
| 2016-9 | m | 16 | B6.129 Mertktm1 Grl/J mutant | none | none | MCAO | Training |
| 2016-9 | m | 16 | B6.129 Mertktm1 Grl/J mutant | none | none | MCAO | Testing |
| 2016-9 | m | 16 | B6 Mfge8 GtByg wildtype | none | none | MCAO | Training |
| 2016-9 | m | 16 | B6 Mfge8 GtByg wildtype | none | none | MCAO | Training |
| 2016-9 | m | 16 | B6.129 Mertktm1 Grl/J mutant | none | none | MCAO | Testing |

Continued Table S1

| | | | | | | | |
|---|---|---|---|---|---|---|---|
| 2016-9 | m | 16 | B6.129 Mertktm1 Grl/J wildtype | none | none | MCAO | Training |
| 2016-9 | m | 17 | B6 Mfge8 GtByg wildtype | none | none | MCAO | Testing |
| 2016-9 | m | 17 | B6 Mfge8 GtByg mutant | none | none | MCAO | Training |
| 2016-9 | m | 17 | Mertk wildtype | none | none | MCAO | Training |
| 2016-9 | m | 17 | B6 Mfge8 GtByg wildtype | none | none | MCAO | Testing |
| 2016-9 | m | 18 | Mfge8 wildtype | none | none | MCAO | Testing |
| 2016-9 | m | 19 | B6 Mfge8 GtByg wildtype | none | none | MCAO | Training |
| 2016-9 | m | 19 | B6 Mfge8 GtByg wildtype | none | none | MCAO | Training |
| 2016-9 | m | 20 | B6.129 Mertktm1 Grl/J wildtype | none | none | MCAO | Training |
| 2016-9 | m | 20 | Mfge8 wildtype | none | none | MCAO | Training |
| 2016-9 | m | 33 | Mfge8 wildtype | none | none | MCAO | Training |
| 2017-1 | m | 16 | Cx30xFlexIL6 wildtype/tg | Tamoxifen | none | MCAO | Training |
| 2017-1 | m | 16 | Cx30xFlexIL6 wildtype/tg | Tamoxifen | none | sham | sham |
| 2017-1 | f | 16 | Cx30xFlexIL6 wildtype/tg | Tamoxifen | none | MCAO | Training |
| 2017-1 | m | 16 | Cx30xFlexIL6 wildtype/tg | Tamoxifen | none | MCAO | Training |
| 2017-1 | f | 16 | Cx30xFlexIL6 wildtype/tg | Tamoxifen | none | MCAO | Testing |
| 2017-1 | f | 16 | Cx30xFlexIL6 wildtype/tg | Tamoxifen | none | MCAO | Testing |
| 2017-1 | m | 16 | Cx30xFlexIL6 wildtype/tg | Tamoxifen | none | sham | sham |
| 2017-1 | m | 17 | Cx30xFlexIL6 Cre/tg | Tamoxifen | none | MCAO | Training |
| 2017-1 | f | 17 | Cx30xFlexIL6 Cre/tg | Tamoxifen | none | MCAO | Testing |
| 2017-1 | f | 17 | Cx30xFlexIL6 Cre/tg | Tamoxifen | none | MCAO | Testing |
| 2017-1 | f | 17 | Cx30xFlexIL6 Cre/tg | Tamoxifen | none | sham | sham |

Continued Table S1

| | | | | | | | |
|---|---|---|---|---|---|---|---|
| 2017-1 | f | 17 | Cx30xFlexIL6 Cre/tg | Tamoxifen | none | sham | sham |
| 2017-1 | f | 18 | Cx30xFlexIL6 Cre/tg | Tamoxifen | none | sham | sham |
| 2017-1 | m | 18 | Cx30xFlexIL6 Cre/tg | Tamoxifen | none | MCAO | Testing |
| 2017-1 | f | 19 | Cx30xFlexIL6 Cre/tg | Tamoxifen | none | sham | sham |
| 2017-1 | m | 19 | Cx30xFlexIL6 Cre/tg | Tamoxifen | none | MCAO | Testing |
| 2017-1 | m | 21 | Cx30xFlexIL6 wildtype/tg | Tamoxifen | none | MCAO | Training |
| 2017-1 | m | 21 | Cx30xFlexIL6 wildtype/tg | Tamoxifen | none | sham | sham |
| 2017-1 | f | 22 | Cx30xFlexIL6 wildtype/tg | Tamoxifen | none | MCAO | Testing |
| 2017-1 | m | 24 | Cx30xFlexIL6 cre/tg | Tamoxifen | none | sham | sham |
| 2017-2 | f | 12 | Cx30xFlexIL6 wildtype/tg | Tamoxifen | none | MCAO | Training |
| 2017-2 | f | 12 | Cx30xFlexIL6 cre/tg | Tamoxifen | none | MCAO | Training |
| 2017-2 | f | 12 | Cx30xFlexIL6 cre/tg | Tamoxifen | none | MCAO | Training |
| 2017-2 | m | 13 | Cx30xFlexIL6 cre/tg | Tamoxifen | none | MCAO | Training |
| 2017-2 | m | 13 | Cx30xFlexIL6 cre/tg | Tamoxifen | none | MCAO | Testing |
| 2017-2 | f | 13 | Cx30xFlexIL6 cre/tg | Tamoxifen | none | MCAO | Testing |
| 2017-2 | m | 13 | Cx30xFlexIL6 wildtype/tg | Tamoxifen | none | MCAO | Training |
| 2017-2 | f | 13 | Cx30xFlexIL6 wildtype/tg | Tamoxifen | none | MCAO | Training |
| 2017-6 | m | 12 | Zfp580 tm1a HO | none | none | sham | sham |
| 2017-6 | m | 12 | Zfp580 tm1a wildtype | none | none | MCAO | Testing |
| 2017-6 | m | 12 | Zfp580 tm1a wildtype | none | none | MCAO | Training |
| 2017-6 | m | 12 | Zfp580 tm1a wildtype | none | none | MCAO | Testing |
| 2017-6 | f | 12 | Zfp580 tm1a wildtype | none | none | MCAO | Training |
| 2017-6 | f | 12 | Zfp580 tm1a HO | none | none | MCAO | Training |
| 2017-6 | m | 12 | Zfp580 tm1a HO | none | none | MCAO | Training |

Continued Table S1

| | | | | | | | |
|---|---|---|---|---|---|---|---|
| 2017-6 | m | 12 | Zfp580 tm1a HO | none | none | MCAO | Training |
| 2017-6 | m | 14 | Zfp580 tm1a wildtype | none | none | MCAO | Testing |
| 2017-6 | m | 15 | Zfp580 tm1a HO | none | none | MCAO | Testing |
| 2017-6 | f | 15 | Zfp580 tm1a HO | none | none | MCAO | Testing |
| 2017-6 | m | 17 | Zfp580 tm1a HO | none | none | MCAO | Training |
| 2017-6 | m | 17 | Zfp580 tm1a wiltdype | none | none | MCAO | Testing |
| 2017-6 | f | 17 | Zfp580 tm1a wildtype | none | none | MCAO | Training |
| 2017-6 | f | 17 | Zfp580 tm1a HO | none | none | MCAO | Training |
| 2017-6 | f | 17 | Zfp580 tm1a wildtype | none | none | MCAO | Training |
| 2017-6 | m | 17 | Zfp580 tm1a HO | none | none | MCAO | Testing |
| 2017-6 | f | 17 | Zfp580 tm1a HO | none | none | MCAO | Testing |
| 2017-6 | f | 19 | Zfp580 tm1a wildtype | none | none | MCAO | Training |
| 2017-6 | f | 19 | Zfp580 tm1a HO | none | none | MCAO | Training |
| 2017-6 | f | 19 | Zfp580 tm1a HO | none | none | MCAO | Training |
| 2017-6 | f | 19 | Zfp580 tm1a wildtype | none | none | MCAO | Testing |
| 2017-6 | f | 19 | Zfp580 tm1a HO | none | none | MCAO | Training |
| 2017-6 | f | 21 | Zfp580 tm1a wildtype | none | none | MCAO | Training |
| 2017-6 | f | 21 | Zfp580 tm1a wildtype | none | none | sham | sham |
| 2017-8 | m | 14 | Cx30xFlexIL6 cre/tg | Tamoxifen | none | MCAO | Training |
| 2017-8 | m | 14 | Cx30xFlexIL6 cre/tg | Tamoxifen | none | MCAO | Training |
| 2017-8 | m | 14 | Cx30xFlexIL6 cre/tg | Tamoxifen | none | MCAO | Testing |
| 2017-8 | m | 14 | Cx30xFlexIL6 cre/tg | Vehikel | none | MCAO | Training |
| 2017-8 | m | 14 | Cx30xFlexIL6 cre/tg | Vehikel | none | MCAO | Training |
| 2017-8 | f | 14 | Cx30xFlexIL6 cre/tg | Vehikel | none | MCAO | Training |

Continued Table S1

| | | | | | | | |
|---|---|---|---|---|---|---|---|
| 2017-8 | m | 15 | Cx30xFlexIL6 cre/tg | Vehikel | none | MCAO | Training |
| 2017-8 | f | 15 | Cx30xFlexIL6 cre/tg | Tamoxifen | none | MCAO | Training |
| 2017-8 | m | 15 | Cx30xFlexIL6 cre/tg | Vehikel | none | MCAO | Training |
| 2017-8 | f | 15 | Cx30xFlexIL6 cre/tg | Tamoxifen | none | MCAO | Training |
| 2017-8 | m | 15 | Cx30xFlexIL6 cre/tg | Tamoxifen | none | MCAO | Training |
| 2017-8 | m | 16 | Cx30xFlexIL6 cre/tg | Tamoxifen | none | MCAO | Testing |
| 2017-8 | f | 16 | Cx30xFlexIL6 cre/tg | Tamoxifen | none | MCAO | Testing |
| 2017-8 | f | 16 | Cx30xFlexIL6 cre/tg | Vehikel | none | MCAO | Testing |
| 2017-8 | m | 16 | Cx30xFlexIL6 cre/tg | Tamoxifen | none | MCAO | Training |
| 2017-8 | f | 16 | Cx30xFlexIL6 cre/tg | Vehikel | none | MCAO | Training |
| 2018-4 | f | 13 | Zfp580 tm1a HO | none | none | MCAO | Testing |
| 2018-4 | m | 13 | Zfp580 tm1a HO | none | none | MCAO | Training |
| 2018-4 | f | 14 | Zfp580 tm1a wildtype | none | none | MCAO | Training |
| 2018-4 | m | 14 | Zfp580 tm1a wildtype | none | none | MCAO | Testing |
| 2018-4 | m | 14 | Zfp580 tm1a HO | none | none | MCAO | Training |
| 2018-4 | m | 14 | Zfp580 tm1a wildtype | none | none | MCAO | Training |
| 2018-4 | m | 16 | Zfp580 tm1a HO | none | none | MCAO | Testing |
| 2018-4 | f | 17 | Zfp580 tm1a HO | none | none | MCAO | Training |
| 2018-4 | m | 17 | Zfp580 tm1a HO | none | none | MCAO | Training |
| 2018-4 | m | 18 | Zfp580 tm1a HO | none | none | MCAO | Testing |
| 2018-4 | m | 18 | Zfp580 tm1a HO | none | none | MCAO | Training |
| 2018-4 | f | 18 | Zfp580 tm1a HO | none | none | MCAO | Training |
| 2018-4 | f | 18 | Zfp580 tm1a wildtype | none | none | MCAO | Training |
| 2018-4 | f | 19 | Zfp580 tm1a wildtype | none | none | MCAO | Training |

Continued Table S1

| | | | | | | | |
|---|---|---|---|---|---|---|---|
| 2018-4 | m | 20 | Zfp580 tm1a wildtype | none | none | MCAO | Training |
| 2018-4 | m | 20 | Zfp580 tm1a wildtype | none | none | MCAO | Training |
| 2018-4 | m | 20 | Zfp580 tm1a wildtype | none | none | MCAO | Training |
| 2018-4 | m | 20 | Zfp580 tm1a wildtype | none | none | MCAO | Training |
| 2018-4 | m | 20 | Zfp580 tm1a wildtype | none | none | MCAO | Training |
| 2018-4 | f | 20 | Zfp580 tm1a HO | none | none | MCAO | Training |
| 2018-4 | f | 20 | Zfp580 tm1a wildtype | none | none | sham | sham |
| 2018-8 | m | 13 | Sorcs wildtype | none | none | MCAO | Testing |
| 2018-8 | m | 14 | Sorcs knockout | none | none | MCAO | Testing |
| 2018-8 | m | 14 | Sorcs wildtype | none | none | sham | sham |
| 2018-8 | m | 14 | Sorcs knockout | none | none | MCAO | Training |
| 2018-8 | m | 14 | Sorcs knockout | none | none | MCAO | Training |
| 2018-8 | m | 14 | Sorcs wildtype | none | none | MCAO | Testing |
| 2018-8 | m | 14 | Sorcs knockout | none | none | MCAO | Training |
| 2018-8 | m | 14 | Sorcs wildtype | none | none | MCAO | Testing |
| 2018-8 | m | 14 | Sorcs knockout | none | none | MCAO | Testing |
| 2018-8 | m | 17 | Sorcs wildtype | none | none | MCAO | Testing |
| 2018-8 | m | 17 | Sorcs wildtype | none | none | MCAO | Training |
| 2019-1 | m | 15 | C57Bl6_N wildtype | none | Kontroll-Peptid | MCAO | Testing |
| 2019-1 | m | 15 | C57Bl6_N wildtype | none | Cycl (RADM) | MCAO | Training |
| 2019-1 | m | 15 | C57Bl6_N wildtype | none | Cycl (RADM) | MCAO | Testing |
| 2019-1 | m | 15 | C57Bl6_N wildtype | none | Kontroll-Peptid | MCAO | Testing |
| 2019-1 | m | 15 | C57Bl6_N wildtype | none | PBS | MCAO | Testing |
| 2019-1 | m | 15 | C57Bl6_N wildtype | none | PBS | MCAO | Training |
| 2019-1 | m | 15 | C57Bl6_N wildtype | none | Cycl (RADM) | MCAO | Training |
| 2019-1 | m | 15 | C57Bl6_N wildtype | none | Cilengitide | MCAO | Training |
| 2019-1 | m | 15 | C57Bl6_N wildtype | none | Cycl (RADM) | MCAO | Testing |
| 2019-1 | m | 15 | C57Bl6_N wildtype | none | Kontroll-Peptid | MCAO | Testing |
| 2019-1 | m | 15 | C57Bl6_N wildtype | none | Cilengitide | MCAO | Training |

Continued Table S1

| | | | | | | | |
|---|---|---|---|---|---|---|---|
| 2019-1 | m | 15 | C57Bl6_N wildtype | none | PBS | MCAO | Training |
| 2019-1 | m | 15 | C57Bl6_N wildtype | none | Cilengitide | MCAO | Training |
| 2019-1 | m | 15 | C57Bl6_N wildtype | none | Cycl (RADM) | MCAO | Training |
| 2019-1 | m | 15 | C57Bl6_N wildtype | none | Cycl (RADM) | MCAO | Training |
| 2019-1 | m | 15 | C57Bl6_N wildtype | none | PBS | MCAO | Training |
| 2019-1 | m | 15 | C57Bl6_N wildtype | none | Kontroll-Peptid | MCAO | Training |
| 2019-1 | m | 15 | C57Bl6_N wildtype | none | Cilengitide | MCAO | Testing |
| 2019-1 | m | 15 | C57Bl6_N wildtype | none | Kontroll-Peptid | MCAO | Training |
| 2019-8 | f | 14 | C57Bl6_N | none | metamizole | MCAO | replication cohort |
| 2019-8 | m | 14 | C57Bl6_N | none | none | MCAO | replication cohort |
| 2019-8 | m | 14 | C57Bl6_N | none | none | MCAO | replication cohort |
| 2019-8 | m | 14 | C57Bl6_N | none | metamizole | MCAO | replication cohort |
| 2019-8 | f | 14 | C57Bl6_N | none | metamizole | MCAO | replication cohort |
| 2019-8 | f | 14 | C57Bl6_N | none | none | MCAO | replication cohort |
| 2019-8 | f | 14 | C57Bl6_N | none | none | MCAO | replication cohort |
| 2019-8 | m | 15 | C57Bl6_N | none | none | MCAO | replication cohort |
| 2019-8 | m | 15 | C57Bl6_N | none | none | MCAO | replication cohort |
| 2019-8 | m | 15 | C57Bl6_N | none | none | MCAO | replication cohort |
| 2019-8 | f | 15 | C57Bl6_N | none | metamizole | MCAO | replication cohort |
| 2019-8 | f | 15 | C57Bl6_N | none | none | MCAO | replication cohort |
| 2019-8 | f | 15 | C57Bl6_N | none | none | MCAO | replication cohort |
| 2019-8 | m | 15 | C57Bl6_N | none | metamizole | MCAO | replication cohort |
| 2019-8 | m | 15 | C57Bl6_N | none | metamizole | MCAO | replication cohort |
| 2019-8 | m | 15 | C57Bl6_N | none | none | MCAO | replication cohort |

Continued Table S1

| | | | | | | | |
|---|---|---|---|---|---|---|---|
| 2019-8 | f | 15 | C57Bl6_N | none | metamizole | MCAO | replication cohort |
| 2019-8 | f | 15 | C57Bl6_N | none | none | MCAO | replication cohort |
| 2019-8 | f | 15 | C57Bl6_N | none | none | MCAO | replication cohort |

**Table S1 Overview over the genotypes, treatments and sex of the mice included in the study.**

| | Total | Female | Male |
|---|---|---|---|
| **MCAO** | 185 | 30 | 135 |
| Baseline age in weeks † | 15.39 (15.00-15.78) | 16.07 (15.34-16.79) | 15.14 (14.68-15.60) |
| Baseline weight in grams | 23.20 (22.85-23.55) | 21.33 (20.81-21.85) | 23.89 (23.51-24.27) |
| weight Day 1 | 22.48 (22.14-22.83) | 20.63 (20.07-21.19) | 23.17 (22.80-23.54) |
| weight Day 2 | 21.39 (21.05-21.73) | 20.05 (19.42-20.68) | 21.89 (21.52-22.26) |
| Neuroscore Day 2 * | 3.90 (3.51-4.30) | 2.84 (2.22-3.46) | 4.33 (3.85-4.81) |

**Table S2:** Overview of the basic characteristics of animals. † denotes age in weeks at the time of the surgery. * denotes the modified DeSimoni neuroscore for the assessment of general deficits. The neuroscore was available for a total of 174 MCAO mice, of which 124 were male and 50 were female.

| **Label** | **Lesion percentage** | **SEM** |
|---|---|---|
| Caudoputamen | 55.75 | 1.37 |
| Claustrum | 40.41 | 2.61 |
| Agranular insular area, dorsal part, layer 6b | 44.63 | 2.95 |
| Fundus of striatum | 44.15 | 2.99 |
| Central amygdalar nucleus, lateral part | 39.92 | 2.92 |
| Visceral area, layer 6b | 47.56 | 3.59 |
| Globus pallidus, external segment | 18.58 | 1.40 |
| Agranular insular area, posterior part, layer 6a | 48.21 | 3.71 |
| Endopiriform nucleus, dorsal part | 35.03 | 2.73 |
| Olfactory tubercle, polymorph layer | 25.42 | 2.01 |
| Gustatory areas, layer 6b | 34.47 | 2.75 |
| Nucleus accumbens | 14.22 | 1.14 |
| Agranular insular area, ventral part, layer 6a | 20.15 | 1.68 |
| Substantia innominata | 14.09 | 1.21 |
| Basic cell groups and regions | 7.77 | 0.67 |
| Piriform area, polymorph layer | 35.15 | 3.13 |
| Agranular insular area, posterior part, layer 5 | 42.03 | 3.77 |
| Agranular insular area, dorsal part, layer 6a | 25.15 | 2.26 |
| Piriform area, pyramidal layer | 31.49 | 2.87 |
| Supplemental somatosensory area, layer 6b | 33.98 | 3.10 |
| Lateral amygdalar nucleus | 34.65 | 3.18 |
| Central amygdalar nucleus, capsular part | 35.22 | 3.30 |
| Visceral area, layer 6a | 39.98 | 3.75 |
| Cerebral cortex | 26.47 | 2.48 |

Continued Table S3

| | | |
|---|---|---|
| Central amygdalar nucleus, medial part | 31.68 | 2.97 |
| Islands of Calleja | 12.27 | 1.17 |
| Olfactory tubercle, pyramidal layer | 15.37 | 1.47 |
| Gustatory areas, layer 6a | 29.37 | 2.82 |
| Endopiriform nucleus, ventral part | 34.82 | 3.38 |
| Piriform area, molecular layer | 27.52 | 2.67 |
| Anterior amygdalar area | 29.33 | 2.90 |
| Agranular insular area, posterior part, layer 2/3 | 37.24 | 3.68 |
| Striatum-like amygdalar nuclei | 24.45 | 2.43 |
| Agranular insular area, ventral part, layer 5 | 22.79 | 2.36 |
| Olfactory areas | 8.37 | 0.87 |
| Supplemental somatosensory area, layer 6a | 32.96 | 3.45 |
| Intercalated amygdalar nucleus | 32.68 | 3.42 |
| Cortical amygdalar area, anterior part, layer 2 | 33.42 | 3.52 |
| Visceral area, layer 5 | 35.08 | 3.69 |
| Basomedial amygdalar nucleus, anterior part | 34.00 | 3.62 |
| Basolateral amygdalar nucleus, anterior part | 32.45 | 3.47 |
| Entorhinal area, lateral part, layer 4/5 | 33.22 | 3.55 |
| Agranular insular area, posterior part, layer 1 | 31.56 | 3.39 |
| Cortical amygdalar area, anterior part, layer 1 | 27.38 | 2.94 |
| Supplemental somatosensory area, layer 5 | 32.09 | 3.59 |
| Visceral area, layer 4 | 32.22 | 3.62 |
| Nucleus of the lateral olfactory tract, pyramidal layer | 30.72 | 3.47 |
| Nucleus of the lateral olfactory tract, layer 3 | 31.28 | 3.54 |
| Perirhinal area, layer 6a | 24.33 | 2.77 |
| Ventral auditory area, layer 6a | 23.92 | 2.72 |

Continued Table S3

| | | |
|---|---|---|
| Visceral area, layer 2/3 | 31.17 | 3.56 |
| Entorhinal area, lateral part, layer 2/3 | 28.74 | 3.28 |
| Supplemental somatosensory area, layer 4 | 30.88 | 3.58 |
| Supplemental somatosensory area, layer 2/3 | 29.81 | 3.47 |
| Agranular insular area, ventral part, layer 2/3 | 23.77 | 2.77 |
| Perirhinal area, layer 5 | 25.73 | 3.02 |
| Ventral auditory area, layer 6b | 19.09 | 2.26 |
| Agranular insular area, dorsal part, layer 5 | 15.24 | 1.81 |
| Gustatory areas, layer 5 | 25.25 | 3.01 |
| Visceral area, layer 1 | 25.69 | 3.07 |
| Piriform-amygdalar area, polymorph layer | 29.81 | 3.57 |
| Ventral auditory area, layer 5 | 25.86 | 3.11 |
| Nucleus of the lateral olfactory tract, molecular layer | 22.69 | 2.75 |
| Olfactory tubercle, molecular layer | 7.92 | 0.96 |
| Perirhinal area, layer 2/3 | 25.04 | 3.04 |
| Piriform-amygdalar area, pyramidal layer | 28.17 | 3.43 |
| Globus pallidus, internal segment | 11.82 | 1.44 |
| Magnocellular nucleus | 10.24 | 1.25 |
| Ventral auditory area, layer 4 | 26.55 | 3.26 |
| Basomedial amygdalar nucleus, posterior part | 27.55 | 3.38 |
| Medial amygdalar nucleus, anterodorsal part | 23.41 | 2.88 |
| Basolateral amygdalar nucleus, ventral part | 28.94 | 3.58 |
| Piriform-amygdalar area, molecular layer | 23.62 | 2.94 |
| Cortical amygdalar area, posterior part, lateral zone, layer 2 | 25.40 | 3.21 |
| Gustatory areas, layer 4 | 24.85 | 3.15 |
| Agranular insular area, ventral part, layer 1 | 21.64 | 2.74 |

Continued Table S3

| | | |
|---|---|---|
| Ventral auditory area, layer 2/3 | 25.78 | 3.27 |
| Entorhinal area, lateral part, layer 2 | 27.27 | 3.47 |
| Primary somatosensory area, nose, layer 6b | 22.08 | 2.81 |
| Bed nucleus of the accessory olfactory tract | 25.16 | 3.24 |
| Perirhinal area, layer 1 | 18.09 | 2.34 |
| Gustatory areas, layer 2/3 | 23.34 | 3.06 |
| Perirhinal area, layer 6b | 20.53 | 2.70 |
| Medial amygdalar nucleus, anteroventral part | 21.33 | 2.82 |
| Temporal association areas, layer 6a | 18.86 | 2.51 |
| Medial amygdalar nucleus, posterodorsal part, sublayer b | 22.60 | 3.01 |
| Cortical amygdalar area, posterior part, lateral zone, layer 3 | 24.20 | 3.25 |
| Gustatory areas, layer 1 | 20.62 | 2.77 |
| Temporal association areas, layer 4 | 20.12 | 2.70 |
| Temporal association areas, layer 5 | 20.15 | 2.71 |
| Basolateral amygdalar nucleus, posterior part | 23.52 | 3.17 |
| Primary auditory area, layer 5 | 22.41 | 3.03 |
| Supplemental somatosensory area, layer 1 | 17.86 | 2.42 |
| Medial amygdalar nucleus, posterodorsal part, sublayer c | 20.15 | 2.73 |
| Primary auditory area, layer 2/3 | 23.53 | 3.20 |
| Primary auditory area, layer 4 | 22.87 | 3.11 |
| Temporal association areas, layer 2/3 | 19.22 | 2.61 |
| Ectorhinal area/Layer 6a | 17.78 | 2.43 |
| Dorsal auditory area, layer 2/3 | 22.60 | 3.10 |
| Dorsal auditory area, layer 4 | 22.77 | 3.13 |
| Medial amygdalar nucleus, posterodorsal part, sublayer a | 20.40 | 2.80 |
| Primary somatosensory area, barrel field, layer 2/3 | 20.35 | 2.81 |
| Continued Table S3 | | |

| | | |
|---|---|---|
| Primary somatosensory area, barrel field, layer 4 | 20.57 | 2.84 |
| Primary somatosensory area, nose, layer 6a | 23.64 | 3.27 |
| Ectorhinal area/Layer 5 | 18.21 | 2.52 |
| Dorsal auditory area, layer 5 | 21.18 | 2.94 |
| Primary somatosensory area, nose, layer 5 | 24.30 | 3.39 |
| Entorhinal area, lateral part, layer 6b | 20.32 | 2.84 |
| Cortical amygdalar area, posterior part, lateral zone, layer 1 | 18.02 | 2.53 |
| Field CA3, stratum lacunosum-moleculare | 23.62 | 3.32 |
| Primary somatosensory area, nose, layer 2/3 | 22.58 | 3.17 |
| Primary somatosensory area, nose, layer 4 | 23.85 | 3.35 |
| Primary somatosensory area, barrel field, layer 5 | 18.79 | 2.65 |
| Primary somatosensory area, barrel field, layer 6a | 15.17 | 2.14 |
| Temporal association areas, layer 1 | 13.81 | 1.95 |
| Ventral auditory area, layer 1 | 18.92 | 2.67 |
| Ventral posterolateral nucleus of the thalamus | 13.02 | 1.85 |
| Field CA2, stratum oriens | 19.72 | 2.81 |
| Field CA2, pyramidal layer | 20.59 | 2.94 |
| Ectorhinal area/Layer 2/3 | 16.48 | 2.36 |
| Ectorhinal area/Layer 1 | 12.11 | 1.74 |
| Dorsal auditory area, layer 1 | 18.15 | 2.61 |
| Field CA3, pyramidal layer | 18.47 | 2.66 |
| Field CA2, stratum lacunosum-moleculare | 20.87 | 3.00 |
| Ectorhinal area/Layer 6b | 17.53 | 2.52 |
| Field CA2, stratum radiatum | 19.88 | 2.86 |
| Primary auditory area, layer 1 | 17.96 | 2.59 |
| Field CA3, stratum oriens | 15.61 | 2.26 |
| Continued Table S3 | | |

| | | |
|---|---|---|
| Reticular nucleus of the thalamus | 10.16 | 1.47 |
| Primary auditory area, layer 6a | 15.65 | 2.27 |
| Field CA3, stratum lucidum | 19.39 | 2.82 |
| Field CA3, stratum radiatum | 20.52 | 2.99 |
| Agranular insular area, dorsal part, layer 2/3 | 12.03 | 1.75 |
| Lateral dorsal nucleus of thalamus | 18.67 | 2.72 |
| Postpiriform transition area, layers 1 | 18.92 | 2.80 |
| Field CA1, stratum oriens | 15.67 | 2.32 |
| Agranular insular area, dorsal part, layer 1 | 10.44 | 1.55 |
| Postpiriform transition area, layers 2 | 19.15 | 2.85 |
| Postpiriform transition area, layers 3 | 19.92 | 2.98 |
| Ventral posteromedial nucleus of the thalamus | 12.40 | 1.85 |
| Field CA1, stratum radiatum | 18.52 | 2.78 |
| Primary somatosensory area, barrel field, layer 6b | 9.59 | 1.44 |
| Dentate gyrus, polymorph layer | 18.43 | 2.78 |
| Cortical amygdalar area, posterior part, medial zone, layer 3 | 17.45 | 2.66 |
| Dentate gyrus, granule cell layer | 15.11 | 2.32 |
| Thalamus | 4.84 | 0.74 |
| Medial amygdalar nucleus, posteroventral part | 17.39 | 2.67 |
| Entorhinal area, lateral part, layer 1 | 11.68 | 1.80 |
| Field CA1, stratum lacunosum-moleculare | 17.52 | 2.72 |
| Field CA1, pyramidal layer | 17.55 | 2.73 |
| Primary somatosensory area, mouth, layer 4 | 18.17 | 2.83 |
| Primary somatosensory area, barrel field, layer 1 | 14.24 | 2.22 |
| Posterior complex of the thalamus | 10.46 | 1.64 |
| Continued Table S3 | | |

| | | |
|---|---|---|
| Primary somatosensory area, mouth, layer 2/3 | 16.88 | 2.65 |
| Temporal association areas, layer 6b | 15.05 | 2.38 |
| Entorhinal area, lateral part, layer 5 | 16.12 | 2.56 |
| Dentate gyrus, molecular layer | 12.53 | 2.00 |
| Primary somatosensory area, mouth, layer 5 | 16.52 | 2.65 |
| Primary somatosensory area, unassigned, layer 6b | 5.98 | 0.96 |
| Entorhinal area, lateral part, layer 3 | 15.37 | 2.48 |
| Entorhinal area, lateral part, layer 4 | 16.22 | 2.62 |
| Dorsal auditory area, layer 6a | 12.60 | 2.04 |
| Posterior amygdalar nucleus | 15.80 | 2.56 |
| Diagonal band nucleus | 1.06 | 0.17 |
| Posterior parietal association areas, layer 4 | 7.37 | 1.20 |
| Entorhinal area, lateral part, layer 2a | 13.60 | 2.24 |
| Cortical amygdalar area, posterior part, medial zone, layer 2 | 13.12 | 2.16 |
| Posterior parietal association areas, layer 5 | 6.06 | 1.00 |
| Posterior parietal association areas, layer 2/3 | 7.17 | 1.18 |
| Primary somatosensory area, mouth, layer 6a | 14.34 | 2.38 |
| Primary somatosensory area, nose, layer 1 | 13.35 | 2.23 |
| Primary somatosensory area, unassigned, layer 4 | 13.71 | 2.29 |
| Entorhinal area, lateral part, layer 2b | 14.28 | 2.41 |
| Primary somatosensory area, unassigned, layer 2/3 | 14.05 | 2.38 |
| Entorhinal area, lateral part, layer 6a | 14.06 | 2.42 |
| Primary somatosensory area, trunk, layer 6b | 3.60 | 0.62 |
| Anteroventral nucleus of thalamus | 11.59 | 2.01 |
| Anterolateral visual area, layer 2/3 | 14.50 | 2.52 |

Continued Table S3

| | | |
|---|---|---|
| Posterior parietal association areas, layer 6b | 4.94 | 0.86 |
| Posterior parietal association areas, layer 1 | 5.05 | 0.88 |
| Primary somatosensory area, unassigned, layer 5 | 12.10 | 2.14 |
| Anterolateral visual area, layer 1 | 10.64 | 1.89 |
| Posterior auditory area, layer 2/3 | 16.26 | 2.89 |
| Anterolateral visual area, layer 4 | 13.96 | 2.49 |
| Posterior auditory area, layer 4 | 16.59 | 2.98 |
| Primary somatosensory area, unassigned, layer 1 | 9.93 | 1.81 |
| Cortical amygdalar area, posterior part, medial zone, layer 1 | 8.91 | 1.63 |
| Ventral anterior-lateral complex of the thalamus | 3.48 | 0.64 |
| Central lateral nucleus of the thalamus | 10.29 | 1.90 |
| Primary somatosensory area, mouth, layer 6b | 9.68 | 1.80 |
| Primary somatosensory area, mouth, layer 1 | 9.46 | 1.77 |
| Posterior auditory area, layer 1 | 12.68 | 2.38 |
| Dorsal auditory area, layer 6b | 8.55 | 1.61 |
| Primary auditory area, layer 6b | 9.83 | 1.85 |
| Posterior auditory area, layer 5 | 15.08 | 2.85 |
| Anterodorsal nucleus | 9.10 | 1.72 |
| Lateral visual area, layer 2/3 | 13.27 | 2.53 |
| Lateral visual area, layer 4 | 14.46 | 2.76 |
| Anterolateral visual area, layer 5 | 12.11 | 2.32 |
| Primary somatosensory area, upper limb, layer 4 | 10.49 | 2.03 |
| Primary somatosensory area, unassigned, layer 6a | 7.76 | 1.50 |
| Subiculum, dorsal part, stratum radiatum | 8.57 | 1.67 |
| Lateral visual area, layer 5 | 13.81 | 2.69 |

Continued Table S3

| | | |
|---|---|---|
| Subiculum, ventral part, stratum radiatum | 12.26 | 2.39 |
| Subiculum, ventral part, molecular layer | 12.56 | 2.45 |
| Primary somatosensory area, upper limb, layer 2/3 | 10.29 | 2.03 |
| Lateral hypothalamic area | 0.48 | 0.09 |
| Posterior parietal association areas, layer 6a | 3.24 | 0.65 |
| Primary somatosensory area, upper limb, layer 5 | 9.10 | 1.83 |
| Hypothalamus | 0.20 | 0.04 |
| Orbital area, lateral part, layer 5 | 2.17 | 0.44 |
| Subiculum, ventral part, pyramidal layer | 10.88 | 2.21 |
| Dorsal part of the lateral geniculate complex | 8.80 | 1.82 |
| Subiculum, dorsal part, molecular layer | 8.07 | 1.68 |
| Lateral visual area, layer 1 | 6.25 | 1.32 |
| Posterolateral visual area, layer 4 | 11.78 | 2.49 |
| Lateral posterior nucleus of the thalamus | 8.41 | 1.78 |
| Primary visual area, layer 2/3 | 7.00 | 1.48 |
| Subiculum, dorsal part, pyramidal layer | 6.45 | 1.37 |
| Primary visual area, layer 4 | 7.74 | 1.66 |
| Medial geniculate complex, ventral part | 6.47 | 1.40 |
| Posterior auditory area, layer 6a | 12.64 | 2.73 |
| Posterolateral visual area, layer 5 | 11.62 | 2.51 |
| Presubiculum, layer 1 | 7.53 | 1.63 |
| Lateral visual area, layer 6a | 10.91 | 2.36 |
| Orbital area, lateral part, layer 2/3 | 2.30 | 0.50 |
| Posterior auditory area, layer 6b | 12.52 | 2.73 |
| Primary visual area, layer 5 | 7.45 | 1.63 |
| Mediodorsal nucleus of the thalamus, lateral part | 4.70 | 1.03 |
| Continued Table S3 | | |

| | | |
|---|---|---|
| Supraoptic nucleus | 1.54 | 0.34 |
| Posterolateral visual area, layer 2/3 | 9.52 | 2.11 |
| Primary somatosensory area, upper limb, layer 6a | 6.64 | 1.50 |
| Presubiculum, layer 2 | 8.25 | 1.88 |
| Orbital area, ventrolateral part, layer 5 | 0.40 | 0.09 |
| Posterolateral visual area, layer 6a | 10.51 | 2.41 |
| Anterolateral visual area, layer 6a | 8.13 | 1.88 |
| Entorhinal area, medial part, dorsal zone, layer 6 | 8.59 | 1.99 |
| Parasubiculum, layer 2 | 8.48 | 1.96 |
| Parasubiculum, layer 3 | 9.39 | 2.18 |
| Primary motor area, Layer 2/3 | 4.48 | 1.05 |
| Primary visual area, layer 1 | 3.95 | 0.93 |
| Entorhinal area, medial part, dorsal zone, layer 5 | 8.42 | 1.98 |
| Primary somatosensory area, upper limb, layer 1 | 6.35 | 1.50 |
| posteromedial visual area, layer 6a | 8.07 | 1.93 |
| Entorhinal area, medial part, dorsal zone, layer 4 | 8.44 | 2.02 |
| Primary visual area, layer 6a | 6.15 | 1.48 |
| Ventral part of the lateral geniculate complex | 5.05 | 1.22 |
| Posterolateral visual area, layer 6b | 9.44 | 2.27 |
| Midbrain | 1.18 | 0.29 |
| Superior colliculus, motor related, intermediate gray layer, sublayer b | 6.38 | 1.54 |
| Medial geniculate complex, dorsal part | 6.40 | 1.55 |
| Postsubiculum, layer 2 | 6.62 | 1.60 |
| Parasubiculum, layer 1 | 6.86 | 1.66 |
| Presubiculum, layer 3 | 7.37 | 1.79 |

Continued Table S3

| | | |
|---|---|---|
| Primary motor area, Layer 1 | 3.38 | 0.82 |
| Postsubiculum, layer 3 | 6.56 | 1.62 |
| Zona incerta | 1.93 | 0.48 |
| posteromedial visual area, layer 5 | 6.88 | 1.71 |
| Lateral visual area, layer 6b | 8.82 | 2.21 |
| Entorhinal area, medial part, dorsal zone, layer 3 | 7.28 | 1.83 |
| Paracentral nucleus | 1.94 | 0.49 |
| Superior colliculus, motor related, intermediate white layer | 6.25 | 1.57 |
| posteromedial visual area, layer 2/3 | 4.01 | 1.01 |
| Primary somatosensory area, trunk, layer 2/3 | 4.02 | 1.02 |
| Orbital area, lateral part, layer 6a | 0.82 | 0.21 |
| Retrosplenial area, dorsal part, layer 5 | 2.18 | 0.56 |
| Entorhinal area, medial part, dorsal zone, layer 2 | 6.28 | 1.60 |
| Anterolateral visual area, layer 6b | 5.79 | 1.48 |
| Entorhinal area, medial part, dorsal zone, layer 1 | 4.12 | 1.06 |
| Superior colliculus, motor related, intermediate gray layer, sublayer a | 7.54 | 1.94 |
| Primary somatosensory area, trunk, layer 6a | 1.61 | 0.41 |
| Suprageniculate nucleus | 6.12 | 1.58 |
| Anterior pretectal nucleus | 4.88 | 1.26 |
| posteromedial visual area, layer 4 | 5.54 | 1.43 |
| Primary somatosensory area, trunk, layer 4 | 3.81 | 0.99 |
| Retrosplenial area, dorsal part, layer 6a | 2.88 | 0.75 |
| Nucleus of the optic tract | 6.80 | 1.77 |
| posteromedial visual area, layer 6b | 7.07 | 1.84 |
| Superior colliculus, motor related, intermediate gray layer, sublayer c | 4.25 | 1.11 |

Continued Table S3

| | | |
|---|---|---|
| Retrosplenial area, dorsal part, layer 2/3 | 1.36 | 0.36 |
| Entorhinal area, medial part, ventral zone, layer 2 | 5.95 | 1.57 |
| Peripeduncular nucleus | 2.71 | 0.72 |
| Posterolateral visual area, layer 1 | 3.55 | 0.95 |
| Primary visual area, layer 6b | 4.10 | 1.10 |
| Posterior limiting nucleus of the thalamus | 3.44 | 0.92 |
| Intergeniculate leaflet of the lateral geniculate complex | 5.25 | 1.41 |
| Lateral habenula | 2.96 | 0.80 |
| Superior colliculus, motor related, deep gray layer | 3.32 | 0.90 |
| Superior colliculus, motor related, intermediate gray layer | 5.38 | 1.47 |
| Entorhinal area, medial part, ventral zone, layer 5/6 | 7.88 | 2.16 |
| Anterior olfactory nucleus, posteroventral part | 1.30 | 0.36 |
| Primary motor area, Layer 5 | 2.33 | 0.64 |
| Subthalamic nucleus | 1.62 | 0.45 |
| Posterior pretectal nucleus | 6.73 | 1.85 |
| Olivary pretectal nucleus | 5.61 | 1.56 |
| Primary somatosensory area, trunk, layer 1 | 3.14 | 0.87 |
| Postsubiculum, layer 1 | 4.57 | 1.28 |
| Mediodorsal nucleus of the thalamus, central part | 3.40 | 0.95 |
| Entorhinal area, medial part, ventral zone, layer 3 | 6.13 | 1.72 |
| Primary somatosensory area, trunk, layer 5 | 2.69 | 0.76 |
| Primary somatosensory area, lower limb, layer 4 | 2.71 | 0.77 |
| Primary somatosensory area, upper limb, layer 6b | 3.58 | 1.02 |
| Medial geniculate complex, medial part | 2.88 | 0.82 |
| posteromedial visual area, layer 1 | 2.48 | 0.71 |
| Retrosplenial area, dorsal part, layer 6b | 1.41 | 0.41 |
| Continued Table S3 | | |

| | | |
|---|---|---|
| Subparafascicular nucleus, parvicellular part | 1.12 | 0.33 |
| Primary somatosensory area, lower limb, layer 2/3 | 2.63 | 0.77 |
| Entorhinal area, medial part, ventral zone, layer 1 | 2.37 | 0.69 |
| Retrosplenial area, dorsal part, layer 1 | 0.77 | 0.22 |
| Entorhinal area, medial part, dorsal zone, layer 2b | 6.28 | 1.85 |
| Superior colliculus, optic layer | 4.35 | 1.30 |
| Superior colliculus, motor related, deep white layer | 2.25 | 0.68 |
| Retrosplenial area, lateral agranular part, layer 6a | 2.74 | 0.84 |
| Orbital area, lateral part, layer 1 | 1.08 | 0.33 |
| Retrosplenial area, lateral agranular part, layer 5 | 2.39 | 0.73 |
| Parafascicular nucleus | 0.25 | 0.08 |
| Primary motor area, Layer 6b | 0.29 | 0.09 |
| Orbital area, ventrolateral part, layer 6a | 0.62 | 0.19 |
| Primary somatosensory area, lower limb, layer 5 | 2.12 | 0.66 |
| Anteromedial nucleus, dorsal part | 1.40 | 0.44 |
| Retrosplenial area, ventral part, layer 2/3 | 2.05 | 0.65 |
| Primary somatosensory area, lower limb, layer 1 | 1.62 | 0.52 |
| Superior colliculus, zonal layer | 3.55 | 1.14 |
| Retrosplenial area, ventral part, layer 5 | 1.75 | 0.56 |
| Superior colliculus, superficial gray layer | 4.28 | 1.38 |
| Medial habenula | 0.53 | 0.17 |
| Lateral terminal nucleus of the accessory optic tract | 1.19 | 0.39 |
| Interanterodorsal nucleus of the thalamus | 1.82 | 0.60 |
| Inferior colliculus, external nucleus | 1.02 | 0.34 |
| Retrosplenial area, ventral part, layer 1 | 1.98 | 0.66 |
| Retrosplenial area, lateral agranular part, layer 6b | 2.08 | 0.70 |

Continued Table S3

| | | |
|---|---|---|
| Periaqueductal gray | 0.36 | 0.12 |
| Retrosplenial area, ventral part, layer 2 | 1.91 | 0.64 |
| Substantia nigra, reticular part | 0.76 | 0.26 |
| Midbrain reticular nucleus | 0.11 | 0.04 |
| Retrosplenial area, ventral part, layer 6a | 2.02 | 0.71 |
| Entorhinal area, medial part, dorsal zone, layer 2a | 3.93 | 1.38 |
| Anteromedial visual area, layer 4 | 1.55 | 0.55 |
| Anteromedial visual area, layer 5 | 0.96 | 0.34 |
| Retrosplenial area, lateral agranular part, layer 2/3 | 1.30 | 0.46 |
| Medial pretectal area | 1.17 | 0.42 |
| Primary somatosensory area, lower limb, layer 6a | 1.47 | 0.53 |
| Fasciola cinerea | 1.27 | 0.47 |
| Anteromedial visual area, layer 6b | 0.46 | 0.17 |
| Primary motor area, Layer 6a | 0.76 | 0.29 |
| Mediodorsal nucleus of the thalamus, medial part | 0.87 | 0.33 |
| Simple lobule, molecular layer | 0.18 | 0.07 |
| Nucleus of the posterior commissure | 0.62 | 0.24 |
| Bed nuclei of the stria terminalis, anterior division, anterolateral area | 0.85 | 0.33 |
| Lateral septal nucleus, caudal (caudodorsal) part | 0.64 | 0.25 |
| Primary somatosensory area, lower limb, layer 6b | 0.69 | 0.27 |
| Subgeniculate nucleus | 2.65 | 1.04 |
| Substantia nigra, compact part | 0.75 | 0.30 |
| Secondary motor area, layer 1 | 0.33 | 0.14 |
| Secondary motor area, layer 2/3 | 0.34 | 0.14 |
| Orbital area, ventrolateral part, layer 2/3 | 0.24 | 0.10 |

Continued Table S3

| | | |
|---|---|---|
| Retrosplenial area, ventral part, layer 6b | 1.02 | 0.43 |
| Anteromedial visual area, layer 2/3 | 1.57 | 0.66 |
| Precommissural nucleus | 0.05 | 0.02 |
| Flocculus, molecular layer | 0.09 | 0.04 |
| Anteromedial visual area, layer 6a | 0.41 | 0.19 |
| Cerebellum | 0.01 | 0.00 |
| Bed nuclei of the stria terminalis, anterior division, oval nucleus | 1.31 | 0.60 |
| Paraventricular nucleus of the thalamus | 0.34 | 0.16 |
| Pons | 0.03 | 0.01 |
| Nucleus of the brachium of the inferior colliculus | 1.50 | 0.71 |
| Lateral preoptic area | 0.41 | 0.19 |
| Lateral septal nucleus, rostral (rostroventral) part | 0.12 | 0.06 |
| Simple lobule, granular layer | 0.09 | 0.04 |
| Lobules IV-V, molecular layer | 0.03 | 0.01 |
| Bed nuclei of the stria terminalis, anterior division, juxtacapsular nucleus | 2.05 | 1.02 |
| Retrosplenial area, lateral agranular part, layer 1 | 0.39 | 0.20 |

**Table S3 Regions with increased T2-intensities in MRI.**
Edema/damage is presented as lesion percentage, meaning percentage of a given region that showed a lesion in tT2w-MR images.

| Label | Rank | Importance |
|---|---|---|
| Caudoputamen | 1 | 0.32308629 |
| genu of corpus callosum | 2 | 0.2332126 |
| Magnocellular nucleus | 3 | 0.21865599 |
| Primary somatosensory area, mouth, layer 6b | 4 | 0.18925805 |
| cranial nerves | 5 | 0.18675973 |
| Substantia innominata | 6 | 0.1851075 |
| Basic cell groups and regions | 7 | 0.17853399 |
| Striatum-like amygdalar nuclei | 8 | 0.16087243 |
| Primary somatosensory area, barrel field, layer 6b | 9 | 0.15391005 |
| Basolateral amygdalar nucleus, anterior part | 10 | 0.15348099 |
| Central amygdalar nucleus, capsular part | 11 | 0.1428494 |
| Islands of Calleja | 12 | 0.13133924 |
| Gustatory areas, layer 6b | 13 | 0.12603834 |
| fiber tracts | 14 | 0.12539283 |
| Primary somatosensory area, nose, layer 6b | 15 | 0.11755629 |
| supra-callosal cerebral white matter | 16 | 0.11395881 |
| corpus callosum body | 17 | 0.11391755 |
| Fundus of striatum | 18 | 0.10824129 |
| Olfactory tubercle, polymorph layer | 19 | 0.10756712 |
| Agranular insular area, dorsal part, layer 6a | 20 | 0.10229244 |
| Entorhinal area, lateral part, layer 6b | 21 | 0.10175451 |
| Supplemental somatosensory area, layer 2/3 | 22 | 0.09889486 |
| anterior commissure temporal limb | 23 | 0.09201019 |

Continued Table S4

| | | |
|---|---|---|
| Primary somatosensory area, nose, layer 6a | 24 | 0.09112598 |
| corpus callosum | 25 | 0.08295043 |
| Intercalated amygdalar nucleus | 26 | 0.07695747 |
| Claustrum | 27 | 0.07074763 |
| lateral forebrain bundle system | 28 | 0.06814444 |
| external capsule | 29 | 0.06672324 |
| Gustatory areas, layer 1 | 30 | 0.06626382 |
| Ventral auditory area, layer 2/3 | 31 | 0.06552788 |
| Primary somatosensory area, unassigned, layer 6a | 32 | 0.06303901 |
| Agranular insular area, ventral part, layer 5 | 33 | 0.06251169 |
| Olfactory tubercle, pyramidal layer | 34 | 0.06074569 |
| Visceral area, layer 4 | 35 | 0.06024039 |
| Ventral auditory area, layer 4 | 36 | 0.06023796 |
| Primary somatosensory area, barrel field, layer 6a | 37 | 0.0600351 |
| cerebrum related | 38 | 0.05941571 |
| Gustatory areas, layer 6a | 39 | 0.05928908 |
| Entorhinal area, lateral part, layer 6a | 40 | 0.05552282 |
| Agranular insular area, dorsal part, layer 6b | 41 | 0.05478822 |
| Anterior amygdalar area | 42 | 0.05442298 |
| Ectorhinal area/Layer 5 | 43 | 0.05199313 |
| corticospinal tract | 44 | 0.05029466 |
| Agranular insular area, posterior part, layer 1 | 45 | 0.04948235 |
| Temporal association areas, layer 1 | 46 | 0.04874207 |
| medial forebrain bundle system | 47 | 0.04870997 |

Continued Table S4

| | | |
|---|---|---|
| Supplemental somatosensory area, layer 4 | 48 | 0.04857274 |
| Agranular insular area, dorsal part, layer 5 | 49 | 0.04831752 |
| Supplemental somatosensory area, layer 6b | 50 | 0.04770422 |
| amygdalar capsule | 51 | 0.04683343 |
| Anterolateral visual area, layer 2/3 | 52 | 0.04644172 |
| Visceral area, layer 2/3 | 53 | 0.04627575 |
| Visceral area, layer 1 | 54 | 0.0451319 |
| Endopiriform nucleus, ventral part | 55 | 0.045062 |
| Gustatory areas, layer 5 | 56 | 0.04502237 |
| Visceral area, layer 6b | 57 | 0.04488031 |
| corpus callosum anterior forceps | 58 | 0.04487091 |
| Diagonal band nucleus | 59 | 0.04485613 |
| Agranular insular area, posterior part, layer 6a | 60 | 0.04373156 |
| Temporal association areas, layer 4 | 61 | 0.04304704 |
| Supplemental somatosensory area, layer 1 | 62 | 0.04086983 |
| alveus | 63 | 0.03928596 |
| internal capsule | 64 | 0.03884521 |
| Ectorhinal area/Layer 6b | 65 | 0.03723898 |
| Primary motor area, Layer 5 | 66 | 0.03679916 |
| Agranular insular area, ventral part, layer 6a | 67 | 0.03597981 |
| Agranular insular area, dorsal part, layer 2/3 | 68 | 0.03149397 |
| Piriform area, pyramidal layer | 69 | 0.02824495 |
| Field CA3, stratum oriens | 70 | 0.02097705 |
| Primary somatosensory area, unassigned, layer 5 | 71 | 0.02072315 |

Continued Table S4

| | | |
|---|---|---|
| Agranular insular area, posterior part, layer 2/3 | 72 | 0.01995215 |
| Field CA3, pyramidal layer | 73 | 0.01334997 |
| Globus pallidus, internal segment | 74 | 0.01334802 |
| Agranular insular area, dorsal part, layer 1 | 75 | 0.01082569 |
| Central amygdalar nucleus, medial part | 76 | 0.01056376 |
| Central amygdalar nucleus, lateral part | 77 | 0.00766024 |
| Nucleus accumbens | 78 | 0.00516727 |
| optic tract | 79 | 0.00468865 |
| Ventral auditory area, layer 5 | 80 | 0.00264418 |
| Olfactory tubercle, molecular layer | 81 | 0.00103826 |

**Table S4 Regions with out-of-bag importance above 0.**

**A**

| Tukey's multiple comparisons test | Mean Diff. | 95% CI of Diff. | Adjusted P Value |
|---|---|---|---|
| Mild vs. Moderate | -12.26 | -26.31 to 1.794 | 0.1185 |
| Mild vs. Moderate to Mild | -3.136 | -14.88 to 8.613 | 0.9474 |
| Mild vs. Severe | -33.2 | -47.03 to -19.38 | <0.0001 |
| Mild vs. Severe to Moderate | -19.59 | -32.07 to -7.097 | 0.0003 |
| Moderate vs. Moderate to Mild | 9.124 | -5.302 to 23.55 | 0.4085 |
| Moderate vs. Severe | -20.94 | -37.10 to -4.780 | 0.0042 |
| Moderate vs. Severe to Moderate | -7.326 | -22.36 to 7.708 | 0.6628 |
| Moderate to Mild vs. Severe | -30.06 | -44.27 to -15.86 | <0.0001 |
| Moderate to Mild vs. Severe to Moderate | -16.45 | -29.36 to -3.543 | 0.0051 |
| Severe vs. Severe to Moderate | 13.61 | -1.207 to 28.44 | 0.088 |

**B**

| Test details | Mean 1 | Mean 2 | Mean Diff. | SE of Diff. |
|---|---|---|---|---|
| Mild vs. Moderate | 14.03 | 26.29 | -12.26 | 5.086 |
| Mild vs. Moderate to Mild | 14.03 | 17.17 | -3.136 | 4.252 |
| Mild vs. Severe | 14.03 | 47.23 | -33.2 | 5.004 |
| Mild vs. Severe to Moderate | 14.03 | 33.62 | -19.59 | 4.52 |
| Moderate vs. Moderate to Mild | 26.29 | 17.17 | 9.124 | 5.221 |
| Moderate vs. Severe | 26.29 | 47.23 | -20.94 | 5.849 |
| Moderate vs. Severe to Moderate | 26.29 | 33.62 | -7.326 | 5.442 |
| Moderate to Mild vs. Severe | 17.17 | 47.23 | -30.06 | 5.141 |
| Moderate to Mild vs. Severe to Moderate | 17.17 | 33.62 | -16.45 | 4.671 |
| Severe vs. Severe to Moderate | 47.23 | 33.62 | 13.61 | 5.364 |

**Table S5 Statistical report on the analysis of severity grades and stroke lesion volumes (A,B)**

| **Šídák's multiple comparisons test** | **Mean Diff.** | **95% CI of Diff.** | **Adjusted P Value** |
|---|---|---|---|
| Mild: Lesion Volume vs. Mild: MRI [Segmented] | -13.99 | -14.98 to -12.99 | <0.0001 |
| Mild: Lesion Volume vs. Moderate to Mild: Lesion Volume | 7.513 | 6.521 to 8.505 | <0.0001 |
| Mild: Lesion Volume vs. Moderate to Mild: MRI [Segmented] | 10.17 | 9.181 to 11.17 | <0.0001 |
| Mild: Lesion Volume vs. Moderate: Lesion Volume | -0.6437 | -1.636 to 0.3482 | 0.7907 |
| Mild: Lesion Volume vs. Moderate: MRI [Segmented] | 6.225 | 5.233 to 7.217 | <0.0001 |
| Mild: Lesion Volume vs. Moderate to Severe: Lesion Volume | 11.01 | 10.02 to 12.00 | <0.0001 |
| Mild: Lesion Volume vs. Moderate to Severe: MRI [Segmented] | 12.51 | 11.51 to 13.50 | <0.0001 |
| Mild: Lesion Volume vs. Severe: Lesion Volume | 13.12 | 12.12 to 14.11 | <0.0001 |
| Mild: Lesion Volume vs. Severe: MRI [Segmented] | 11.4 | 10.41 to 12.40 | <0.0001 |
| Mild: MRI [Segmented] vs. Moderate to Mild: Lesion Volume | 21.5 | 20.51 to 22.49 | <0.0001 |
| Mild: MRI [Segmented] vs. Moderate to Mild: MRI [Segmented] | 24.16 | 23.17 to 25.15 | <0.0001 |
| Mild: MRI [Segmented] vs. Moderate: Lesion Volume | 13.34 | 12.35 to 14.33 | <0.0001 |
| Mild: MRI [Segmented] vs. Moderate: MRI [Segmented] | 20.21 | 19.22 to 21.20 | <0.0001 |
| Mild: MRI [Segmented] vs. Moderate to Severe: Lesion Volume | 24.99 | 24.00 to 25.99 | <0.0001 |
| Mild: MRI [Segmented] vs. Moderate to Severe: MRI [Segmented] | 26.49 | 25.50 to 27.48 | <0.0001 |
| Mild: MRI [Segmented] vs. Severe: Lesion Volume | 27.1 | 26.11 to 28.09 | <0.0001 |
| Mild: MRI [Segmented] vs. Severe: MRI [Segmented] | 25.39 | 24.40 to 26.38 | <0.0001 |
| Moderate to Mild: Lesion Volume vs. Moderate to Mild: MRI [Segmented] | 2.66 | 1.668 to 3.652 | <0.0001 |
| Moderate to Mild: Lesion Volume vs. Moderate: Lesion Volume | -8.157 | -9.149 to -7.165 | <0.0001 |
| Moderate to Mild: Lesion Volume vs. Moderate: MRI [Segmented] | -1.289 | -2.281 to -0.2968 | 0.0011 |

Continued Table S6

| | | | |
|---|---|---|---|
| Moderate to Mild: Lesion Volume vs. Moderate to Severe: Lesion Volume | 3.494 | 2.502 to 4.486 | <0.0001 |
| Moderate to Mild: Lesion Volume vs. Moderate to Severe: MRI [Segmented] | 4.992 | 4.000 to 5.984 | <0.0001 |
| Moderate to Mild: Lesion Volume vs. Severe: Lesion Volume | 5.602 | 4.610 to 6.594 | <0.0001 |
| Moderate to Mild: Lesion Volume vs. Severe: MRI [Segmented] | 3.891 | 2.899 to 4.883 | <0.0001 |
| Moderate to Mild: MRI [Segmented] vs. Moderate: Lesion Volume | -10.82 | -11.81 to -9.825 | <0.0001 |
| Moderate to Mild: MRI [Segmented] vs. Moderate: MRI [Segmented] | -3.949 | -4.941 to -2.957 | <0.0001 |
| Moderate to Mild: MRI [Segmented] vs. Moderate to Severe: Lesion Volume | 0.834 | -0.1579 to 1.826 | 0.2421 |
| Moderate to Mild: MRI [Segmented] vs. Moderate to Severe: MRI [Segmented] | 2.332 | 1.340 to 3.324 | <0.0001 |
| Moderate to Mild: MRI [Segmented] vs. Severe: Lesion Volume | 2.942 | 1.950 to 3.934 | <0.0001 |
| Moderate to Mild: MRI [Segmented] vs. Severe: MRI [Segmented] | 1.231 | 0.2395 to 2.223 | 0.0025 |
| Moderate: Lesion Volume vs. Moderate: MRI [Segmented] | 6.868 | 5.876 to 7.860 | <0.0001 |
| Moderate: Lesion Volume vs. Moderate to Severe: Lesion Volume | 11.65 | 10.66 to 12.64 | <0.0001 |
| Moderate: Lesion Volume vs. Moderate to Severe: MRI [Segmented] | 13.15 | 12.16 to 14.14 | <0.0001 |
| Moderate: Lesion Volume vs. Severe: Lesion Volume | 13.76 | 12.77 to 14.75 | <0.0001 |
| Moderate: Lesion Volume vs. Severe: MRI [Segmented] | 12.05 | 11.06 to 13.04 | <0.0001 |
| Moderate: MRI [Segmented] vs. Moderate to Severe: Lesion Volume | 4.783 | 3.791 to 5.775 | <0.0001 |
| Moderate: MRI [Segmented] vs. Moderate to Severe: MRI [Segmented] | 6.281 | 5.289 to 7.273 | <0.0001 |
| Moderate: MRI [Segmented] vs. Severe: Lesion Volume | 6.891 | 5.899 to 7.883 | <0.0001 |
| Moderate: MRI [Segmented] vs. Severe: MRI [Segmented] | 5.18 | 4.188 to 6.172 | <0.0001 |

Continued Table S6

| | | | |
|---|---|---|---|
| Moderate to Severe: Lesion Volume vs. Moderate to Severe: MRI [Segmented] | 1.498 | 0.5063 to 2.490 | <0.0001 |
| Moderate to Severe: Lesion Volume vs. Severe: Lesion Volume | 2.108 | 1.116 to 3.100 | <0.0001 |
| Moderate to Severe: Lesion Volume vs. Severe: MRI [Segmented] | 0.3974 | -0.5945 to 1.389 | >0.9999 |
| Moderate to Severe: MRI [Segmented] vs. Severe: Lesion Volume | 0.6101 | -0.3819 to 1.602 | 0.8721 |
| Moderate to Severe: MRI [Segmented] vs. Severe: MRI [Segmented] | -1.101 | -2.093 to -0.1088 | 0.0139 |
| Severe: Lesion Volume vs. Severe: MRI [Segmented] | -1.711 | -2.703 to -0.7189 | <0.0001 |

| **Test details** | **Mean 1** | **Mean 2** | **Mean Diff.** | **SE of Diff.** | **t** | **DF** |
|---|---|---|---|---|---|---|
| Mild: Lesion Volume vs. Mild: MRI [Segmented] | 23.99 | 37.97 | -13.99 | 0.3031 | 46.15 | 490 |
| Mild: Lesion Volume vs. Moderate to Mild: Lesion Volume | 23.99 | 16.47 | 7.513 | 0.3031 | 24.79 | 490 |
| Mild: Lesion Volume vs. Moderate to Mild: MRI [Segmented] | 23.99 | 13.82 | 10.17 | 0.3031 | 33.57 | 490 |
| Mild: Lesion Volume vs. Moderate: Lesion Volume | 23.99 | 24.63 | -0.6437 | 0.3031 | 2.124 | 490 |
| Mild: Lesion Volume vs. Moderate: MRI [Segmented] | 23.99 | 17.76 | 6.225 | 0.3031 | 20.54 | 490 |
| Mild: Lesion Volume vs. Moderate to Severe: Lesion Volume | 23.99 | 12.98 | 11.01 | 0.3031 | 36.32 | 490 |
| Mild: Lesion Volume vs. Moderate to Severe: MRI [Segmented] | 23.99 | 11.48 | 12.51 | 0.3031 | 41.26 | 490 |
| Mild: Lesion Volume vs. Severe: Lesion Volume | 23.99 | 10.87 | 13.12 | 0.3031 | 43.28 | 490 |
| Mild: Lesion Volume vs. Severe: MRI [Segmented] | 23.99 | 12.58 | 11.4 | 0.3031 | 37.63 | 490 |
| Mild: MRI [Segmented] vs. Moderate to Mild: Lesion Volume | 37.97 | 16.47 | 21.5 | 0.3031 | 70.94 | 490 |
| Continued Table S6 | | | | | | |

| | | | | | | |
|---|---|---|---|---|---|---|
| Mild: MRI [Segmented] vs. Moderate to Mild: MRI [Segmented] | 37.97 | 13.82 | 24.16 | 0.3031 | 79.72 | 490 |
| Mild: MRI [Segmented] vs. Moderate: Lesion Volume | 37.97 | 24.63 | 13.34 | 0.3031 | 44.02 | 490 |
| Mild: MRI [Segmented] vs. Moderate: MRI [Segmented] | 37.97 | 17.76 | 20.21 | 0.3031 | 66.69 | 490 |
| Mild: MRI [Segmented] vs. Moderate to Severe: Lesion Volume | 37.97 | 12.98 | 24.99 | 0.3031 | 82.47 | 490 |
| Mild: MRI [Segmented] vs. Moderate to Severe: MRI [Segmented] | 37.97 | 11.48 | 26.49 | 0.3031 | 87.41 | 490 |
| Mild: MRI [Segmented] vs. Severe: Lesion Volume | 37.97 | 10.87 | 27.1 | 0.3031 | 89.42 | 490 |
| Mild: MRI [Segmented] vs. Severe: MRI [Segmented] | 37.97 | 12.58 | 25.39 | 0.3031 | 83.78 | 490 |
| Moderate to Mild: Lesion Volume vs. Moderate to Mild: MRI [Segmented] | 16.47 | 13.82 | 2.66 | 0.3031 | 8.777 | 490 |
| Moderate to Mild: Lesion Volume vs. Moderate: Lesion Volume | 16.47 | 24.63 | -8.157 | 0.3031 | 26.92 | 490 |
| Moderate to Mild: Lesion Volume vs. Moderate: MRI [Segmented] | 16.47 | 17.76 | -1.289 | 0.3031 | 4.252 | 490 |
| Moderate to Mild: Lesion Volume vs. Moderate to Severe: Lesion Volume | 16.47 | 12.98 | 3.494 | 0.3031 | 11.53 | 490 |
| Moderate to Mild: Lesion Volume vs. Moderate to Severe: MRI [Segmented] | 16.47 | 11.48 | 4.992 | 0.3031 | 16.47 | 490 |
| Moderate to Mild: Lesion Volume vs. Severe: Lesion Volume | 16.47 | 10.87 | 5.602 | 0.3031 | 18.49 | 490 |
| Moderate to Mild: Lesion Volume vs. Severe: MRI [Segmented] | 16.47 | 12.58 | 3.891 | 0.3031 | 12.84 | 490 |
| Moderate to Mild: MRI [Segmented] vs. Moderate: Lesion Volume | 13.82 | 24.63 | -10.82 | 0.3031 | 35.69 | 490 |

Continued Table S6

| | | | | | | |
|---|---|---|---|---|---|---|
| Moderate to Mild: MRI [Segmented] vs. Moderate: MRI [Segmented] | 13.82 | 17.76 | -3.949 | 0.3031 | 13.03 | 490 |
| Moderate to Mild: MRI [Segmented] vs. Moderate to Severe: Lesion Volume | 13.82 | 12.98 | 0.834 | 0.3031 | 2.752 | 490 |
| Moderate to Mild: MRI [Segmented] vs. Moderate to Severe: MRI [Segmented] | 13.82 | 11.48 | 2.332 | 0.3031 | 7.696 | 490 |
| Moderate to Mild: MRI [Segmented] vs. Severe: Lesion Volume | 13.82 | 10.87 | 2.942 | 0.3031 | 9.708 | 490 |
| Moderate to Mild: MRI [Segmented] vs. Severe: MRI [Segmented] | 13.82 | 12.58 | 1.231 | 0.3031 | 4.063 | 490 |
| Moderate: Lesion Volume vs. Moderate: MRI [Segmented] | 24.63 | 17.76 | 6.868 | 0.3031 | 22.66 | 490 |
| Moderate: Lesion Volume vs. Moderate to Severe: Lesion Volume | 24.63 | 12.98 | 11.65 | 0.3031 | 38.44 | 490 |
| Moderate: Lesion Volume vs. Moderate to Severe: MRI [Segmented] | 24.63 | 11.48 | 13.15 | 0.3031 | 43.39 | 490 |
| Moderate: Lesion Volume vs. Severe: Lesion Volume | 24.63 | 10.87 | 13.76 | 0.3031 | 45.4 | 490 |
| Moderate: Lesion Volume vs. Severe: MRI [Segmented] | 24.63 | 12.58 | 12.05 | 0.3031 | 39.76 | 490 |
| Moderate: MRI [Segmented] vs. Moderate to Severe: Lesion Volume | 17.76 | 12.98 | 4.783 | 0.3031 | 15.78 | 490 |
| Moderate: MRI [Segmented] vs. Moderate to Severe: MRI [Segmented] | 17.76 | 11.48 | 6.281 | 0.3031 | 20.72 | 490 |
| Moderate: MRI [Segmented] vs. Severe: Lesion Volume | 17.76 | 10.87 | 6.891 | 0.3031 | 22.74 | 490 |
| Moderate: MRI [Segmented] vs. Severe: MRI [Segmented] | 17.76 | 12.58 | 5.18 | 0.3031 | 17.09 | 490 |

Continued Table S6

| | | | | | | |
|---|---|---|---|---|---|---|
| Moderate to Severe: Lesion Volume vs. Moderate to Severe: MRI [Segmented] | 12.98 | 11.48 | 1.498 | 0.3031 | 4.943 | 490 |
| Moderate to Severe: Lesion Volume vs. Severe: Lesion Volume | 12.98 | 10.87 | 2.108 | 0.3031 | 6.956 | 490 |
| Moderate to Severe: Lesion Volume vs. Severe: MRI [Segmented] | 12.98 | 12.58 | 0.3974 | 0.3031 | 1.311 | 490 |
| Moderate to Severe: MRI [Segmented] vs. Severe: Lesion Volume | 11.48 | 10.87 | 0.6101 | 0.3031 | 2.013 | 490 |
| Moderate to Severe: MRI [Segmented] vs. Severe: MRI [Segmented] | 11.48 | 12.58 | -1.101 | 0.3031 | 3.632 | 490 |
| Severe: Lesion Volume vs. Severe: MRI [Segmented] | 10.87 | 12.58 | -1.711 | 0.3031 | 5.645 | 490 |

**Table S6 Full statistical report on the analysis of prediction accuracy and severity grade for the subacute deficit.**

| **Šídák's multiple comparisons test** | **Mean Diff.** | **95% CI of Diff.** | **Adjusted P Value** |
|---|---|---|---|
| Mild: Subacute Deficit vs. Mild: MRI [Segmented] | -3.853 | -5.602 to -2.105 | <0.0001 |
| Mild: Subacute Deficit vs. Mild: Lesion Volume | -5.732 | -7.480 to -3.984 | <0.0001 |
| Mild: Subacute Deficit vs. Moderate to Mild: Subacute Deficit | 12.16 | 10.41 to 13.90 | <0.0001 |
| Mild: Subacute Deficit vs. Moderate to Mild: MRI [Segmented] | 16.56 | 14.81 to 18.31 | <0.0001 |
| Mild: Subacute Deficit vs. Moderate to Mild: Lesion Volume | 15.28 | 13.53 to 17.03 | <0.0001 |
| Mild: Subacute Deficit vs. Moderate: Subacute Deficit | 22.54 | 20.79 to 24.28 | <0.0001 |
| Mild: Subacute Deficit vs. Moderate: MRI [Segmented] | 8.424 | 6.676 to 10.17 | <0.0001 |
| Mild: Subacute Deficit vs. Moderate: Lesion Volume | 1.963 | 0.2143 to 3.711 | 0.0096 |
| Mild: Subacute Deficit vs. Moderate to Severe: Subacute Deficit | 17.2 | 15.45 to 18.94 | <0.0001 |
| Mild: Subacute Deficit vs. Moderate to Severe: MRI [Segmented] | 2.834 | 1.085 to 4.582 | <0.0001 |
| Mild: Subacute Deficit vs. Moderate to Severe: Lesion Volume | 2.337 | 0.5887 to 4.085 | 0.0004 |
| Mild: Subacute Deficit vs. Severe: Subacute Deficit | 4.231 | 2.483 to 5.980 | <0.0001 |
| Mild: Subacute Deficit vs. Severe: MRI [Segmented] | -8.166 | -9.914 to -6.417 | <0.0001 |
| Mild: Subacute Deficit vs. Severe: Lesion Volume | -11.03 | -12.78 to -9.283 | <0.0001 |
| Mild: MRI [Segmented] vs. Mild: Lesion Volume | -1.879 | -3.627 to - 0.1305 | 0.0188 |
| Mild: MRI [Segmented] vs. Moderate to Mild: Subacute Deficit | 16.01 | 14.26 to 17.76 | <0.0001 |
| Mild: MRI [Segmented] vs. Moderate to Mild: MRI [Segmented] | 20.41 | 18.67 to 22.16 | <0.0001 |
| Mild: MRI [Segmented] vs. Moderate to Mild: Lesion Volume | 19.13 | 17.38 to 20.88 | <0.0001 |
| Mild: MRI [Segmented] vs. Moderate: Subacute Deficit | 26.39 | 24.64 to 28.14 | <0.0001 |
| Mild: MRI [Segmented] vs. Moderate: MRI [Segmented] | 12.28 | 10.53 to 14.03 | <0.0001 |

Continued Table S7

| | | | |
|---|---|---|---|
| Mild: MRI [Segmented] vs. Moderate: Lesion Volume | 5.816 | 4.068 to 7.564 | <0.0001 |
| Mild: MRI [Segmented] vs. Moderate to Severe: Subacute Deficit | 21.05 | 19.30 to 22.80 | <0.0001 |
| Mild: MRI [Segmented] vs. Moderate to Severe: MRI [Segmented] | 6.687 | 4.939 to 8.435 | <0.0001 |
| Mild: MRI [Segmented] vs. Moderate to Severe: Lesion Volume | 6.19 | 4.442 to 7.939 | <0.0001 |
| Mild: MRI [Segmented] vs. Severe: Subacute Deficit | 8.085 | 6.336 to 9.833 | <0.0001 |
| Mild: MRI [Segmented] vs. Severe: MRI [Segmented] | -4.312 | -6.061 to -2.564 | <0.0001 |
| Mild: MRI [Segmented] vs. Severe: Lesion Volume | -7.178 | -8.927 to -5.430 | <0.0001 |
| Mild: Lesion Volume vs. Moderate to Mild: Subacute Deficit | 17.89 | 16.14 to 19.64 | <0.0001 |
| Mild: Lesion Volume vs. Moderate to Mild: MRI [Segmented] | 22.29 | 20.54 to 24.04 | <0.0001 |
| Mild: Lesion Volume vs. Moderate to Mild: Lesion Volume | 21.01 | 19.26 to 22.76 | <0.0001 |
| Mild: Lesion Volume vs. Moderate: Subacute Deficit | 28.27 | 26.52 to 30.02 | <0.0001 |
| Mild: Lesion Volume vs. Moderate: MRI [Segmented] | 14.16 | 12.41 to 15.90 | <0.0001 |
| Mild: Lesion Volume vs. Moderate: Lesion Volume | 7.695 | 5.946 to 9.443 | <0.0001 |
| Mild: Lesion Volume vs. Moderate to Severe: Subacute Deficit | 22.93 | 21.18 to 24.68 | <0.0001 |
| Mild: Lesion Volume vs. Moderate to Severe: MRI [Segmented] | 8.566 | 6.818 to 10.31 | <0.0001 |
| Mild: Lesion Volume vs. Moderate to Severe: Lesion Volume | 8.069 | 6.321 to 9.817 | <0.0001 |
| Mild: Lesion Volume vs. Severe: Subacute Deficit | 9.963 | 8.215 to 11.71 | <0.0001 |
| Mild: Lesion Volume vs. Severe: MRI [Segmented] | -2.434 | -4.182 to -0.6853 | 0.0001 |
| Mild: Lesion Volume vs. Severe: Lesion Volume | -5.3 | -7.048 to -3.551 | <0.0001 |
| Moderate to Mild: Subacute Deficit vs. Moderate to Mild: MRI [Segmented] | 4.406 | 2.657 to 6.154 | <0.0001 |
| Moderate to Mild: Subacute Deficit vs. Moderate to Mild: Lesion Volume | 3.122 | 1.374 to 4.870 | <0.0001 |

Continued Table S7

| | | | |
|---|---|---|---|
| Moderate to Mild: Subacute Deficit vs. Moderate: Subacute Deficit | 10.38 | 8.631 to 12.13 | <0.0001 |
| Moderate to Mild: Subacute Deficit vs. Moderate: MRI [Segmented] | -3.731 | -5.480 to -1.983 | <0.0001 |
| Moderate to Mild: Subacute Deficit vs. Moderate: Lesion Volume | -10.19 | -11.94 to -8.445 | <0.0001 |
| Moderate to Mild: Subacute Deficit vs. Moderate to Severe: Subacute Deficit | 5.04 | 3.292 to 6.789 | <0.0001 |
| Moderate to Mild: Subacute Deficit vs. Moderate to Severe: MRI [Segmented] | -9.322 | -11.07 to -7.574 | <0.0001 |
| Moderate to Mild: Subacute Deficit vs. Moderate to Severe: Lesion Volume | -9.819 | -11.57 to -8.070 | <0.0001 |
| Moderate to Mild: Subacute Deficit vs. Severe: Subacute Deficit | -7.924 | -9.672 to -6.176 | <0.0001 |
| Moderate to Mild: Subacute Deficit vs. Severe: MRI [Segmented] | -20.32 | -22.07 to -18.57 | <0.0001 |
| Moderate to Mild: Subacute Deficit vs. Severe: Lesion Volume | -23.19 | -24.94 to -21.44 | <0.0001 |
| Moderate to Mild: MRI [Segmented] vs. Moderate to Mild: Lesion Volume | -1.284 | -3.032 to 0.4647 | 0.6635 |
| Moderate to Mild: MRI [Segmented] vs. Moderate: Subacute Deficit | 5.974 | 4.226 to 7.722 | <0.0001 |
| Moderate to Mild: MRI [Segmented] vs. Moderate: MRI [Segmented] | -8.137 | -9.885 to -6.389 | <0.0001 |
| Moderate to Mild: MRI [Segmented] vs. Moderate: Lesion Volume | -14.6 | -16.35 to -12.85 | <0.0001 |
| Moderate to Mild: MRI [Segmented] vs. Moderate to Severe: Subacute Deficit | 0.6347 | -1.114 to 2.383 | >0.9999 |
| Moderate to Mild: MRI [Segmented] vs. Moderate to Severe: MRI [Segmented] | -13.73 | -15.48 to -11.98 | <0.0001 |
| Moderate to Mild: MRI [Segmented] vs. Moderate to Severe: Lesion Volume | -14.22 | -15.97 to -12.48 | <0.0001 |
| Moderate to Mild: MRI [Segmented] vs. Severe: Subacute Deficit | -12.33 | -14.08 to -10.58 | <0.0001 |

Continued Table S7

| | | | |
|---|---|---|---|
| Moderate to Mild: MRI [Segmented] vs. Severe: MRI [Segmented] | -24.73 | -26.48 to -22.98 | <0.0001 |
| Moderate to Mild: MRI [Segmented] vs. Severe: Lesion Volume | -27.59 | -29.34 to -25.84 | <0.0001 |
| Moderate to Mild: Lesion Volume vs. Moderate: Subacute Deficit | 7.258 | 5.509 to 9.006 | <0.0001 |
| Moderate to Mild: Lesion Volume vs. Moderate: MRI [Segmented] | -6.853 | -8.602 to -5.105 | <0.0001 |
| Moderate to Mild: Lesion Volume vs. Moderate: Lesion Volume | -13.32 | -15.06 to -11.57 | <0.0001 |
| Moderate to Mild: Lesion Volume vs. Moderate to Severe: Subacute Deficit | 1.918 | 0.1700 to 3.667 | 0.0138 |
| Moderate to Mild: Lesion Volume vs. Moderate to Severe: MRI [Segmented] | -12.44 | -14.19 to -10.70 | <0.0001 |
| Moderate to Mild: Lesion Volume vs. Moderate to Severe: Lesion Volume | -12.94 | -14.69 to -11.19 | <0.0001 |
| Moderate to Mild: Lesion Volume vs. Severe: Subacute Deficit | -11.05 | -12.79 to -9.298 | <0.0001 |
| Moderate to Mild: Lesion Volume vs. Severe: MRI [Segmented] | -23.44 | -25.19 to -21.70 | <0.0001 |
| Moderate to Mild: Lesion Volume vs. Severe: Lesion Volume | -26.31 | -28.06 to -24.56 | <0.0001 |
| Moderate: Subacute Deficit vs. Moderate: MRI [Segmented] | -14.11 | -15.86 to -12.36 | <0.0001 |
| Moderate: Subacute Deficit vs. Moderate: Lesion Volume | -20.57 | -22.32 to -18.82 | <0.0001 |
| Moderate: Subacute Deficit vs. Moderate to Severe: Subacute Deficit | -5.339 | -7.088 to -3.591 | <0.0001 |
| Moderate: Subacute Deficit vs. Moderate to Severe: MRI [Segmented] | -19.7 | -21.45 to -17.95 | <0.0001 |
| Moderate: Subacute Deficit vs. Moderate to Severe: Lesion Volume | -20.2 | -21.95 to -18.45 | <0.0001 |
| Moderate: Subacute Deficit vs. Severe: Subacute Deficit | -18.3 | -20.05 to -16.56 | <0.0001 |
| Moderate: Subacute Deficit vs. Severe: MRI [Segmented] | -30.7 | -32.45 to -28.95 | <0.0001 |

Continued Table S7

| | | | |
|---|---|---|---|
| Moderate: Subacute Deficit vs. Severe: Lesion Volume | -33.57 | -35.32 to -31.82 | <0.0001 |
| Moderate: MRI [Segmented] vs. Moderate: Lesion Volume | -6.462 | -8.210 to -4.713 | <0.0001 |
| Moderate: MRI [Segmented] vs. Moderate to Severe: Subacute Deficit | 8.772 | 7.023 to 10.52 | <0.0001 |
| Moderate: MRI [Segmented] vs. Moderate to Severe: MRI [Segmented] | -5.591 | -7.339 to -3.842 | <0.0001 |
| Moderate: MRI [Segmented] vs. Moderate to Severe: Lesion Volume | -6.087 | -7.836 to -4.339 | <0.0001 |
| Moderate: MRI [Segmented] vs. Severe: Subacute Deficit | -4.193 | -5.941 to -2.445 | <0.0001 |
| Moderate: MRI [Segmented] vs. Severe: MRI [Segmented] | -16.59 | -18.34 to -14.84 | <0.0001 |
| Moderate: MRI [Segmented] vs. Severe: Lesion Volume | -19.46 | -21.20 to -17.71 | <0.0001 |
| Moderate: Lesion Volume vs. Moderate to Severe: Subacute Deficit | 15.23 | 13.49 to 16.98 | <0.0001 |
| Moderate: Lesion Volume vs. Moderate to Severe: MRI [Segmented] | 0.8712 | -0.8771 to 2.619 | 0.9999 |
| Moderate: Lesion Volume vs. Moderate to Severe: Lesion Volume | 0.3745 | -1.374 to 2.123 | >0.9999 |
| Moderate: Lesion Volume vs. Severe: Subacute Deficit | 2.269 | 0.5205 to 4.017 | 0.0007 |
| Moderate: Lesion Volume vs. Severe: MRI [Segmented] | -10.13 | -11.88 to -8.380 | <0.0001 |
| Moderate: Lesion Volume vs. Severe: Lesion Volume | -12.99 | -14.74 to -11.25 | <0.0001 |
| Moderate to Severe: Subacute Deficit vs. Moderate to Severe: MRI [Segmented] | -14.36 | -16.11 to -12.61 | <0.0001 |
| Moderate to Severe: Subacute Deficit vs. Moderate to Severe: Lesion Volume | -14.86 | -16.61 to -13.11 | <0.0001 |
| Moderate to Severe: Subacute Deficit vs. Severe: Subacute Deficit | -12.96 | -14.71 to -11.22 | <0.0001 |
| Moderate to Severe: Subacute Deficit vs. Severe: MRI [Segmented] | -25.36 | -27.11 to -23.61 | <0.0001 |
| Moderate to Severe: Subacute Deficit vs. Severe: Lesion Volume | -28.23 | -29.98 to -26.48 | <0.0001 |

Continued Table S7

| | | | |
|---|---|---|---|
| Moderate to Severe: MRI [Segmented] vs. Moderate to Severe: Lesion Volume | -0.4967 | -2.245 to 1.252 | >0.9999 |
| Moderate to Severe: MRI [Segmented] vs. Severe: Subacute Deficit | 1.398 | -0.3506 to 3.146 | 0.4241 |
| Moderate to Severe: MRI [Segmented] vs. Severe: MRI [Segmented] | -11 | -12.75 to -9.251 | <0.0001 |
| Moderate to Severe: MRI [Segmented] vs. Severe: Lesion Volume | -13.87 | -15.61 to -12.12 | <0.0001 |
| Moderate to Severe: Lesion Volume vs. Severe: Subacute Deficit | 1.894 | 0.1460 to 3.643 | 0.0166 |
| Moderate to Severe: Lesion Volume vs. Severe: MRI [Segmented] | -10.5 | -12.25 to -8.754 | <0.0001 |
| Moderate to Severe: Lesion Volume vs. Severe: Lesion Volume | -13.37 | -15.12 to -11.62 | <0.0001 |
| Severe: Subacute Deficit vs. Severe: MRI [Segmented] | -12.4 | -14.15 to -10.65 | <0.0001 |
| Severe: Subacute Deficit vs. Severe: Lesion Volume | -15.26 | -17.01 to -13.51 | <0.0001 |
| Severe: MRI [Segmented] vs. Severe: Lesion Volume | -2.866 | -4.614 to -1.118 | <0.0001 |

| **Test details** | **Mean 1** | **Mean 2** | **Mean Diff.** | **SE of Diff.** | **N1** | **N2** | **t** | **DF** |
|---|---|---|---|---|---|---|---|---|
| Mild: Subacute Deficit vs. Mild: MRI [Segmented] | 28.86 | 32.71 | -3.853 | 0.4991 | 50 | 50 | 7.72 | 735 |
| Mild: Subacute Deficit vs. Mild: Lesion Volume | 28.86 | 34.59 | -5.732 | 0.4991 | 50 | 50 | 11.48 | 735 |
| Mild: Subacute Deficit vs. Moderate to Mild: Subacute Deficit | 28.86 | 16.7 | 12.16 | 0.4991 | 50 | 50 | 24.35 | 735 |
| Mild: Subacute Deficit vs. Moderate to Mild: MRI [Segmented] | 28.86 | 12.3 | 16.56 | 0.4991 | 50 | 50 | 33.18 | 735 |
| Mild: Subacute Deficit vs. Moderate to Mild: Lesion Volume | 28.86 | 13.58 | 15.28 | 0.4991 | 50 | 50 | 30.61 | 735 |
| Mild: Subacute Deficit vs. Moderate: Subacute Deficit | 28.86 | 6.323 | 22.54 | 0.4991 | 50 | 50 | 45.15 | 735 |

| | | | | | | | | |
|---|---|---|---|---|---|---|---|---|
| Mild: Subacute Deficit vs. Moderate: MRI [Segmented] | 28.86 | 20.43 | 8.424 | 0.4991 | 50 | 50 | 16.88 | 735 |
| Mild: Subacute Deficit vs. Moderate: Lesion Volume | 28.86 | 26.9 | 1.963 | 0.4991 | 50 | 50 | 3.932 | 735 |
| Mild: Subacute Deficit vs. Moderate to Severe: Subacute Deficit | 28.86 | 11.66 | 17.2 | 0.4991 | 50 | 50 | 34.45 | 735 |
| Mild: Subacute Deficit vs. Moderate to Severe: MRI [Segmented] | 28.86 | 26.02 | 2.834 | 0.4991 | 50 | 50 | 5.677 | 735 |
| Mild: Subacute Deficit vs. Moderate to Severe: Lesion Volume | 28.86 | 26.52 | 2.337 | 0.4991 | 50 | 50 | 4.682 | 735 |
| Mild: Subacute Deficit vs. Severe: Subacute Deficit | 28.86 | 24.63 | 4.231 | 0.4991 | 50 | 50 | 8.478 | 735 |
| Mild: Subacute Deficit vs. Severe: MRI [Segmented] | 28.86 | 37.02 | -8.166 | 0.4991 | 50 | 50 | 16.36 | 735 |
| Mild: Subacute Deficit vs. Severe: Lesion Volume | 28.86 | 39.89 | -11.03 | 0.4991 | 50 | 50 | 22.1 | 735 |
| Mild: MRI [Segmented] vs. Mild: Lesion Volume | 32.71 | 34.59 | -1.879 | 0.4991 | 50 | 50 | 3.764 | 735 |
| Mild: MRI [Segmented] vs. Moderate to Mild: Subacute Deficit | 32.71 | 16.7 | 16.01 | 0.4991 | 50 | 50 | 32.07 | 735 |
| Mild: MRI [Segmented] vs. Moderate to Mild: MRI [Segmented] | 32.71 | 12.3 | 20.41 | 0.4991 | 50 | 50 | 40.9 | 735 |
| Mild: MRI [Segmented] vs. Moderate to Mild: Lesion Volume | 32.71 | 13.58 | 19.13 | 0.4991 | 50 | 50 | 38.33 | 735 |
| Mild: MRI [Segmented] vs. Moderate: Subacute Deficit | 32.71 | 6.323 | 26.39 | 0.4991 | 50 | 50 | 52.87 | 735 |
| Mild: MRI [Segmented] vs. Moderate: MRI [Segmented] | 32.71 | 20.43 | 12.28 | 0.4991 | 50 | 50 | 24.6 | 735 |
| Mild: MRI [Segmented] vs. Moderate: Lesion Volume | 32.71 | 26.9 | 5.816 | 0.4991 | 50 | 50 | 11.65 | 735 |

Continued Table S7

| | | | | | | | | |
|---|---|---|---|---|---|---|---|---|
| Mild: MRI [Segmented] vs. Moderate to Severe: Subacute Deficit | 32.71 | 11.66 | 21.05 | 0.4991 | 50 | 50 | 42.17 | 735 |
| Mild: MRI [Segmented] vs. Moderate to Severe: MRI [Segmented] | 32.71 | 26.02 | 6.687 | 0.4991 | 50 | 50 | 13.4 | 735 |
| Mild: MRI [Segmented] vs. Moderate to Severe: Lesion Volume | 32.71 | 26.52 | 6.19 | 0.4991 | 50 | 50 | 12.4 | 735 |
| Mild: MRI [Segmented] vs. Severe: Subacute Deficit | 32.71 | 24.63 | 8.085 | 0.4991 | 50 | 50 | 16.2 | 735 |
| Mild: MRI [Segmented] vs. Severe: MRI [Segmented] | 32.71 | 37.02 | -4.312 | 0.4991 | 50 | 50 | 8.64 | 735 |
| Mild: MRI [Segmented] vs. Severe: Lesion Volume | 32.71 | 39.89 | -7.178 | 0.4991 | 50 | 50 | 14.38 | 735 |
| Mild: Lesion Volume vs. Moderate to Mild: Subacute Deficit | 34.59 | 16.7 | 17.89 | 0.4991 | 50 | 50 | 35.84 | 735 |
| Mild: Lesion Volume vs. Moderate to Mild: MRI [Segmented] | 34.59 | 12.3 | 22.29 | 0.4991 | 50 | 50 | 44.67 | 735 |
| Mild: Lesion Volume vs. Moderate to Mild: Lesion Volume | 34.59 | 13.58 | 21.01 | 0.4991 | 50 | 50 | 42.09 | 735 |
| Mild: Lesion Volume vs. Moderate: Subacute Deficit | 34.59 | 6.323 | 28.27 | 0.4991 | 50 | 50 | 56.63 | 735 |
| Mild: Lesion Volume vs. Moderate: MRI [Segmented] | 34.59 | 20.43 | 14.16 | 0.4991 | 50 | 50 | 28.36 | 735 |
| Mild: Lesion Volume vs. Moderate: Lesion Volume | 34.59 | 26.9 | 7.695 | 0.4991 | 50 | 50 | 15.42 | 735 |
| Mild: Lesion Volume vs. Moderate to Severe: Subacute Deficit | 34.59 | 11.66 | 22.93 | 0.4991 | 50 | 50 | 45.94 | 735 |
| Mild: Lesion Volume vs. Moderate to Severe: MRI [Segmented] | 34.59 | 26.02 | 8.566 | 0.4991 | 50 | 50 | 17.16 | 735 |

Continued Table S7

| | | | | | | | | |
|---|---|---|---|---|---|---|---|---|
| Mild: Lesion Volume vs. Moderate to Severe: Lesion Volume | 34.59 | 26.52 | 8.069 | 0.4991 | 50 | 50 | 16.17 | 735 |
| Mild: Lesion Volume vs. Severe: Subacute Deficit | 34.59 | 24.63 | 9.963 | 0.4991 | 50 | 50 | 19.96 | 735 |
| Mild: Lesion Volume vs. Severe: MRI [Segmented] | 34.59 | 37.02 | -2.434 | 0.4991 | 50 | 50 | 4.876 | 735 |
| Mild: Lesion Volume vs. Severe: Lesion Volume | 34.59 | 39.89 | -5.3 | 0.4991 | 50 | 50 | 10.62 | 735 |
| Moderate to Mild: Subacute Deficit vs. Moderate to Mild: MRI [Segmented] | 16.7 | 12.3 | 4.406 | 0.4991 | 50 | 50 | 8.827 | 735 |
| Moderate to Mild: Subacute Deficit vs. Moderate to Mild: Lesion Volume | 16.7 | 13.58 | 3.122 | 0.4991 | 50 | 50 | 6.255 | 735 |
| Moderate to Mild: Subacute Deficit vs. Moderate: Subacute Deficit | 16.7 | 6.323 | 10.38 | 0.4991 | 50 | 50 | 20.8 | 735 |
| Moderate to Mild: Subacute Deficit vs. Moderate: MRI [Segmented] | 16.7 | 20.43 | -3.731 | 0.4991 | 50 | 50 | 7.476 | 735 |
| Moderate to Mild: Subacute Deficit vs. Moderate: Lesion Volume | 16.7 | 26.9 | -10.19 | 0.4991 | 50 | 50 | 20.42 | 735 |
| Moderate to Mild: Subacute Deficit vs. Moderate to Severe: Subacute Deficit | 16.7 | 11.66 | 5.04 | 0.4991 | 50 | 50 | 10.1 | 735 |
| Moderate to Mild: Subacute Deficit vs. Moderate to Severe: MRI [Segmented] | 16.7 | 26.02 | -9.322 | 0.4991 | 50 | 50 | 18.68 | 735 |
| Moderate to Mild: Subacute Deficit vs. Moderate to Severe: Lesion Volume | 16.7 | 26.52 | -9.819 | 0.4991 | 50 | 50 | 19.67 | 735 |
| Moderate to Mild: Subacute Deficit vs. Severe: Subacute Deficit | 16.7 | 24.63 | -7.924 | 0.4991 | 50 | 50 | 15.88 | 735 |

Continued Table S7

| | | | | | | | | |
|---|---|---|---|---|---|---|---|---|
| Moderate to Mild: Subacute Deficit vs. Severe: MRI [Segmented] | 16.7 | 37.02 | -20.32 | 0.4991 | 50 | 50 | 40.71 | 735 |
| Moderate to Mild: Subacute Deficit vs. Severe: Lesion Volume | 16.7 | 39.89 | -23.19 | 0.4991 | 50 | 50 | 46.46 | 735 |
| Moderate to Mild: MRI [Segmented] vs. Moderate to Mild: Lesion Volume | 12.3 | 13.58 | -1.284 | 0.4991 | 50 | 50 | 2.572 | 735 |
| Moderate to Mild: MRI [Segmented] vs. Moderate: Subacute Deficit | 12.3 | 6.323 | 5.974 | 0.4991 | 50 | 50 | 11.97 | 735 |
| Moderate to Mild: MRI [Segmented] vs. Moderate: MRI [Segmented] | 12.3 | 20.43 | -8.137 | 0.4991 | 50 | 50 | 16.3 | 735 |
| Moderate to Mild: MRI [Segmented] vs. Moderate: Lesion Volume | 12.3 | 26.9 | -14.6 | 0.4991 | 50 | 50 | 29.25 | 735 |
| Moderate to Mild: MRI [Segmented] vs. Moderate to Severe: Subacute Deficit | 12.3 | 11.66 | 0.6347 | 0.4991 | 50 | 50 | 1.272 | 735 |
| Moderate to Mild: MRI [Segmented] vs. Moderate to Severe: Lesion Volume | 12.3 | 26.52 | -14.22 | 0.4991 | 50 | 50 | 28.5 | 735 |
| Moderate to Mild: MRI [Segmented] vs. Severe: Subacute Deficit | 12.3 | 24.63 | -12.33 | 0.4991 | 50 | 50 | 24.7 | 735 |
| Moderate to Mild: MRI [Segmented] vs. Severe: MRI [Segmented] | 12.3 | 37.02 | -24.73 | 0.4991 | 50 | 50 | 49.54 | 735 |
| Moderate to Mild: MRI [Segmented] vs. Severe: Lesion Volume | 12.3 | 39.89 | -27.59 | 0.4991 | 50 | 50 | 55.28 | 735 |
| Moderate to Mild: Lesion Volume vs. Moderate: Subacute Deficit | 13.58 | 6.323 | 7.258 | 0.4991 | 50 | 50 | 14.54 | 735 |

Continued Table S7

| | | | | | | | | |
|---|---|---|---|---|---|---|---|---|
| Moderate to Mild: Lesion Volume vs. Moderate: MRI [Segmented] | 13.58 | 20.43 | -6.853 | 0.4991 | 50 | 50 | 13.73 | 735 |
| Moderate to Mild: Lesion Volume vs. Moderate: Lesion Volume | 13.58 | 26.9 | -13.32 | 0.4991 | 50 | 50 | 26.68 | 735 |
| Moderate to Mild: Lesion Volume vs. Moderate to Severe: Subacute Deficit | 13.58 | 11.66 | 1.918 | 0.4991 | 50 | 50 | 3.843 | 735 |
| Moderate to Mild: Lesion Volume vs. Moderate to Severe: MRI [Segmented] | 13.58 | 26.02 | -12.44 | 0.4991 | 50 | 50 | 24.93 | 735 |
| Moderate to Mild: Lesion Volume vs. Moderate to Severe: Lesion Volume | 13.58 | 26.52 | -12.94 | 0.4991 | 50 | 50 | 25.93 | 735 |
| Moderate to Mild: Lesion Volume vs. Severe: Subacute Deficit | 13.58 | 24.63 | -11.05 | 0.4991 | 50 | 50 | 22.13 | 735 |
| Moderate to Mild: Lesion Volume vs. Severe: MRI [Segmented] | 13.58 | 37.02 | -23.44 | 0.4991 | 50 | 50 | 46.97 | 735 |
| Moderate to Mild: Lesion Volume vs. Severe: Lesion Volume | 13.58 | 39.89 | -26.31 | 0.4991 | 50 | 50 | 52.71 | 735 |
| Moderate: Subacute Deficit vs. Moderate: MRI [Segmented] | 6.323 | 20.43 | -14.11 | 0.4991 | 50 | 50 | 28.27 | 735 |
| Moderate: Subacute Deficit vs. Moderate: Lesion Volume | 6.323 | 26.9 | -20.57 | 0.4991 | 50 | 50 | 41.22 | 735 |
| Moderate: Subacute Deficit vs. Moderate to Severe: Subacute Deficit | 6.323 | 11.66 | -5.339 | 0.4991 | 50 | 50 | 10.7 | 735 |
| Moderate: Subacute Deficit vs. Moderate to Severe: MRI [Segmented] | 6.323 | 26.02 | -19.7 | 0.4991 | 50 | 50 | 39.47 | 735 |
| Moderate: Subacute Deficit vs. Moderate to Severe: Lesion Volume | 6.323 | 26.52 | -20.2 | 0.4991 | 50 | 50 | 40.47 | 735 |
| Continued Table S7 | | | | | | | | |

| | | | | | | | | |
|---|---|---|---|---|---|---|---|---|
| Moderate: Subacute Deficit vs. Severe: Subacute Deficit | 6.323 | 24.63 | -18.3 | 0.4991 | 50 | 50 | 36.67 | 735 |
| Moderate: Subacute Deficit vs. Severe: MRI [Segmented] | 6.323 | 37.02 | -30.7 | 0.4991 | 50 | 50 | 61.51 | 735 |
| Moderate: Subacute Deficit vs. Severe: Lesion Volume | 6.323 | 39.89 | -33.57 | 0.4991 | 50 | 50 | 67.25 | 735 |
| Moderate: MRI [Segmented] vs. Moderate: Lesion Volume | 20.43 | 26.9 | -6.462 | 0.4991 | 50 | 50 | 12.95 | 735 |
| Moderate: MRI [Segmented] vs. Moderate to Severe: Subacute Deficit | 20.43 | 11.66 | 8.772 | 0.4991 | 50 | 50 | 17.57 | 735 |
| Moderate: MRI [Segmented] vs. Moderate to Severe: MRI [Segmented] | 20.43 | 26.02 | -5.591 | 0.4991 | 50 | 50 | 11.2 | 735 |
| Moderate: MRI [Segmented] vs. Moderate to Severe: Lesion Volume | 20.43 | 26.52 | -6.087 | 0.4991 | 50 | 50 | 12.2 | 735 |
| Moderate: MRI [Segmented] vs. Severe: Subacute Deficit | 20.43 | 24.63 | -4.193 | 0.4991 | 50 | 50 | 8.401 | 735 |
| Moderate: MRI [Segmented] vs. Severe: MRI [Segmented] | 20.43 | 37.02 | -16.59 | 0.4991 | 50 | 50 | 33.24 | 735 |
| Moderate: MRI [Segmented] vs. Severe: Lesion Volume | 20.43 | 39.89 | -19.46 | 0.4991 | 50 | 50 | 38.98 | 735 |
| Moderate: Lesion Volume vs. Moderate to Severe: Subacute Deficit | 26.9 | 11.66 | 15.23 | 0.4991 | 50 | 50 | 30.52 | 735 |
| Moderate: Lesion Volume vs. Moderate to Severe: MRI [Segmented] | 26.9 | 26.02 | 0.8712 | 0.4991 | 50 | 50 | 1.745 | 735 |
| Moderate: Lesion Volume vs. Moderate to Severe: Lesion Volume | 26.9 | 26.52 | 0.3745 | 0.4991 | 50 | 50 | 0.7503 | 735 |
| Moderate: Lesion Volume vs. Severe: Subacute Deficit | 26.9 | 24.63 | 2.269 | 0.4991 | 50 | 50 | 4.546 | 735 |
| Moderate: Lesion Volume vs. Severe: MRI [Segmented] | 26.9 | 37.02 | -10.13 | 0.4991 | 50 | 50 | 20.29 | 735 |

Continued Table S7

| | | | | | | | | |
|---|---|---|---|---|---|---|---|---|
| Moderate: Lesion Volume vs. Severe: Lesion Volume | 26.9 | 39.89 | -12.99 | 0.4991 | 50 | 50 | 26.03 | 735 |
| Moderate to Severe: Subacute Deficit vs. Moderate to Severe: MRI [Segmented] | 11.66 | 26.02 | -14.36 | 0.4991 | 50 | 50 | 28.78 | 735 |
| Moderate to Severe: Subacute Deficit vs. Moderate to Severe: Lesion Volume | 11.66 | 26.52 | -14.86 | 0.4991 | 50 | 50 | 29.77 | 735 |
| Moderate to Severe: Subacute Deficit vs. Severe: Subacute Deficit | 11.66 | 24.63 | -12.96 | 0.4991 | 50 | 50 | 25.98 | 735 |
| Moderate to Severe: Subacute Deficit vs. Severe: MRI [Segmented] | 11.66 | 37.02 | -25.36 | 0.4991 | 50 | 50 | 50.81 | 735 |
| Moderate to Severe: Subacute Deficit vs. Severe: Lesion Volume | 11.66 | 39.89 | -28.23 | 0.4991 | 50 | 50 | 56.56 | 735 |
| Moderate to Severe: MRI [Segmented] vs. Moderate to Severe: Lesion Volume | 26.02 | 26.52 | -0.4967 | 0.4991 | 50 | 50 | 0.9951 | 735 |
| Moderate to Severe: MRI [Segmented] vs. Severe: Subacute Deficit | 26.02 | 24.63 | 1.398 | 0.4991 | 50 | 50 | 2.8 | 735 |
| Moderate to Severe: MRI [Segmented] vs. Severe: MRI [Segmented] | 26.02 | 37.02 | -11 | 0.4991 | 50 | 50 | 22.04 | 735 |
| Moderate to Severe: MRI [Segmented] vs. Severe: Lesion Volume | 26.02 | 39.89 | -13.87 | 0.4991 | 50 | 50 | 27.78 | 735 |
| Moderate to Severe: Lesion Volume vs. Severe: Subacute Deficit | 26.52 | 24.63 | 1.894 | 0.4991 | 50 | 50 | 3.795 | 735 |
| Moderate to Severe: Lesion Volume vs. Severe: MRI [Segmented] | 26.52 | 37.02 | -10.5 | 0.4991 | 50 | 50 | 21.04 | 735 |
| Moderate to Severe: Lesion Volume vs. Severe: Lesion Volume | 26.52 | 39.89 | -13.37 | 0.4991 | 50 | 50 | 26.78 | 735 |

Continued Table S7

| | | | | | | | | |
|---|---|---|---|---|---|---|---|---|
| Severe: Subacute Deficit vs. Severe: MRI [Segmented] | 24.63 | 37.02 | -12.4 | 0.4991 | 50 | 50 | 24.84 | 735 |
| Severe: Subacute Deficit vs. Severe: Lesion Volume | 24.63 | 39.89 | -15.26 | 0.4991 | 50 | 50 | 30.58 | 735 |
| Severe: MRI [Segmented] vs. Severe: Lesion Volume | 37.02 | 39.89 | -2.866 | 0.4991 | 50 | 50 | 5.742 | 735 |

**Table S7 Full statistical report on the analysis of prediction accuracy and severity grade for the residual deficit.**

| Šídák's multiple comparisons test | Mean Diff. | 95.00% CI of Diff. | Adjusted P Value |
|---|---|---|---|
| Mild: Lesion Volume vs. Mild: MRI [Segmented] | -20.63 | -21.88 to -19.39 | <0.0001 |
| Mild: Lesion Volume vs. Moderate to Mild: Lesion Volume | -25.24 | -26.49 to -23.99 | <0.0001 |
| Mild: Lesion Volume vs. Moderate to Mild: MRI [Segmented] | -13.26 | -14.51 to -12.02 | <0.0001 |
| Mild: Lesion Volume vs. Moderate: Lesion Volume | -7.857 | -9.105 to -6.610 | <0.0001 |
| Mild: Lesion Volume vs. Moderate: MRI [Segmented] | -0.3505 | -1.598 to 0.8970 | >0.9999 |
| Mild: Lesion Volume vs. Moderate to Severe: Lesion Volume | 3.616 | 2.369 to 4.864 | <0.0001 |
| Mild: Lesion Volume vs. Moderate to Severe: MRI [Segmented] | 4.773 | 3.526 to 6.021 | <0.0001 |
| Mild: Lesion Volume vs. Severe: Lesion Volume | 12.66 | 11.41 to 13.91 | <0.0001 |
| Mild: Lesion Volume vs. Severe: MRI [Segmented] | 2.137 | 0.8896 to 3.384 | <0.0001 |
| Mild:MRI [Segmented] vs. Moderate to Mild: Lesion Volume | -4.604 | -5.851 to -3.356 | <0.0001 |
| Mild: MRI [Segmented] vs. Moderate to Mild: MRI [Segmented] | 7.371 | 6.124 to 8.619 | <0.0001 |
| Mild: MRI [Segmented] vs. Moderate: Lesion Volume | 12.78 | 11.53 to 14.02 | <0.0001 |
| Mild: MRI [Segmented] vs. Moderate: MRI [Segmented] | 20.28 | 19.04 to 21.53 | <0.0001 |
| Mild: MRI [Segmented] vs. Moderate to Severe: Lesion Volume | 24.25 | 23.00 to 25.50 | <0.0001 |
| Mild: MRI [Segmented] vs. Moderate to Severe: MRI [Segmented] | 25.41 | 24.16 to 26.66 | <0.0001 |
| Mild: MRI [Segmented] vs. Severe: Lesion Volume | 33.29 | 32.05 to 34.54 | <0.0001 |
| Mild: MRI [Segmented] vs. Severe: MRI [Segmented] | 22.77 | 21.52 to 24.02 | <0.0001 |
| Moderate to Mild: Lesion Volume vs. Moderate to Mild: MRI [Segmented] | 11.98 | 10.73 to 13.22 | <0.0001 |
| Moderate to Mild: Lesion Volume vs. Moderate: Lesion Volume | 17.38 | 16.13 to 18.63 | <0.0001 |
| Moderate to Mild: Lesion Volume vs. Moderate: MRI [Segmented] | 24.89 | 23.64 to 26.14 | <0.0001 |
| Moderate to Mild: Lesion Volume vs. Moderate to Severe: Lesion Volume | 28.85 | 27.61 to 30.10 | <0.0001 |
| Moderate to Mild: Lesion Volume vs. Moderate to Severe: MRI [Segmented] | 30.01 | 28.76 to 31.26 | <0.0001 |
| Moderate to Mild: Lesion Volume vs. Severe: Lesion Volume | 37.9 | 36.65 to 39.15 | <0.0001 |
| Moderate to Mild: Lesion Volume vs. Severe: MRI [Segmented] | 27.38 | 26.13 to 28.62 | <0.0001 |
| Moderate to Mild: MRI [Segmented] vs. Moderate: Lesion Volume | 5.406 | 4.159 to 6.653 | <0.0001 |
| Moderate to Mild: MRI [Segmented] vs. Moderate: MRI [Segmented] | 12.91 | 11.67 to 14.16 | <0.0001 |
| Moderate to Mild: MRI [Segmented] vs. Moderate to Severe: Lesion Volume | 16.88 | 15.63 to 18.13 | <0.0001 |

Continued Table S8

| | | | |
|---|---|---|---|
| Moderate to Mild: MRI [Segmented] vs. Moderate to Severe: MRI [Segmented] | 18.04 | 16.79 to 19.28 | <0.0001 |
| Moderate to Mild: MRI [Segmented] vs. Severe: Lesion Volume | 25.92 | 24.68 to 27.17 | <0.0001 |
| Moderate to Mild: MRI [Segmented] vs. Severe: MRI [Segmented] | 15.4 | 14.15 to 16.65 | <0.0001 |
| Moderate: Lesion Volume vs. Moderate: MRI [Segmented] | 7.507 | 6.259 to 8.754 | <0.0001 |
| Moderate: Lesion Volume vs. Moderate to Severe: Lesion Volume | 11.47 | 10.23 to 12.72 | <0.0001 |
| Moderate: Lesion Volume vs. Moderate to Severe: MRI [Segmented] | 12.63 | 11.38 to 13.88 | <0.0001 |
| Moderate: Lesion Volume vs. Severe: Lesion Volume | 20.52 | 19.27 to 21.76 | <0.0001 |
| Moderate: Lesion Volume vs. Severe: MRI [Segmented] | 9.994 | 8.747 to 11.24 | <0.0001 |
| Moderate: MRI [Segmented] vs. Moderate to Severe: Lesion Volume | 3.967 | 2.719 to 5.214 | <0.0001 |
| Moderate: MRI [Segmented] vs. Moderate to Severe: MRI [Segmented] | 5.124 | 3.876 to 6.371 | <0.0001 |
| Moderate: MRI [Segmented] vs. Severe: Lesion Volume | 13.01 | 11.76 to 14.26 | <0.0001 |
| Moderate: MRI [Segmented] vs. Severe: MRI [Segmented] | 2.487 | 1.240 to 3.735 | <0.0001 |
| Moderate to Severe: Lesion Volume vs. Moderate to Severe: MRI [Segmented] | 1.157 | -0.09038 to 2.404 | 0.1076 |
| Moderate to Severe: Lesion Volume vs. Severe: Lesion Volume | 9.043 | 7.796 to 10.29 | <0.0001 |
| Moderate to Severe: Lesion Volume vs. Severe: MRI [Segmented] | -1.479 | -2.727 to -0.2317 | 0.0053 |
| Moderate to Severe: MRI [Segmented] vs. Severe: Lesion Volume | 7.886 | 6.639 to 9.134 | <0.0001 |
| Moderate to Severe: MRI [Segmented] vs. Severe: MRI [Segmented] | -2.636 | -3.884 to -1.389 | <0.0001 |
| Severe: Lesion Volume vs. Severe: MRI [Segmented] | -10.52 | -11.77 to -9.275 | <0.0001 |

| **Test details** | **Mean 1** | **Mean 2** | **Mean Diff.** | **SE of diff.** | **N1** | **N2** | **t** | **DF** |
|---|---|---|---|---|---|---|---|---|
| Mild: Lesion Volume vs. Mild: MRI [Segmented] | 18.86 | 39.5 | -20.63 | 0.3811 | 50 | 50 | 54.14 | 490 |
| Mild: Lesion Volume vs. Moderate to Mild: Lesion Volume | 18.86 | 44.1 | -25.24 | 0.3811 | 50 | 50 | 66.22 | 490 |
| Mild: Lesion Volume vs. Moderate to Mild: MRI [Segmented] | 18.86 | 32.13 | -13.26 | 0.3811 | 50 | 50 | 34.8 | 490 |
| Mild: Lesion Volume vs. Moderate: Lesion Volume | 18.86 | 26.72 | -7.857 | 0.3811 | 50 | 50 | 20.62 | 490 |
| Mild: Lesion Volume vs. Moderate: MRI [Segmented] | 18.86 | 19.21 | -0.3505 | 0.3811 | 50 | 50 | 0.9195 | 490 |
| Mild: Lesion Volume vs. Moderate to Severe: Lesion Volume | 18.86 | 15.25 | 3.616 | 0.3811 | 50 | 50 | 9.488 | 490 |

Continued Table S8

| | | | | | | | | |
|---|---|---|---|---|---|---|---|---|
| Mild: Lesion Volume vs. Moderate to Severe: MRI [Segmented] | 18.86 | 14.09 | 4.773 | 0.3811 | 50 | 50 | 12.52 | 490 |
| Mild: Lesion Volume vs. Severe: Lesion Volume | 18.86 | 6.205 | 12.66 | 0.3811 | 50 | 50 | 33.22 | 490 |
| Mild: Lesion Volume vs. Severe: MRI [Segmented] | 18.86 | 16.73 | 2.137 | 0.3811 | 50 | 50 | 5.607 | 490 |
| Mild: MRI [Segmented] vs. Moderate to Mild: Lesion Volume | 39.5 | 44.1 | -4.604 | 0.3811 | 50 | 50 | 12.08 | 490 |
| Mild: MRI [Segmented] vs. Moderate to Mild: MRI [Segmented] | 39.5 | 32.13 | 7.371 | 0.3811 | 50 | 50 | 19.34 | 490 |
| Mild: MRI [Segmented] vs. Moderate: Lesion Volume | 39.5 | 26.72 | 12.78 | 0.3811 | 50 | 50 | 33.53 | 490 |
| Mild: MRI [Segmented] vs. Moderate: MRI [Segmented] | 39.5 | 19.21 | 20.28 | 0.3811 | 50 | 50 | 53.22 | 490 |
| Mild: MRI [Segmented] vs. Moderate to Severe: Lesion Volume | 39.5 | 15.25 | 24.25 | 0.3811 | 50 | 50 | 63.63 | 490 |
| Mild: MRI [Segmented] vs. Moderate to Severe: MRI [Segmented] | 39.5 | 14.09 | 25.41 | 0.3811 | 50 | 50 | 66.67 | 490 |
| Mild: MRI [Segmented] vs. Severe: Lesion Volume | 39.5 | 6.205 | 33.29 | 0.3811 | 50 | 50 | 87.36 | 490 |
| Mild: MRI [Segmented] vs. Severe: MRI [Segmented] | 39.5 | 16.73 | 22.77 | 0.3811 | 50 | 50 | 59.75 | 490 |
| Moderate to Mild: Lesion Volume vs. Moderate to Mild: MRI [Segmented] | 44.1 | 32.13 | 11.98 | 0.3811 | 50 | 50 | 31.42 | 490 |
| Moderate to Mild: Lesion Volume vs. Moderate: Lesion Volume | 44.1 | 26.72 | 17.38 | 0.3811 | 50 | 50 | 45.61 | 490 |
| Moderate to Mild: Lesion Volume vs. Moderate: MRI [Segmented] | 44.1 | 19.21 | 24.89 | 0.3811 | 50 | 50 | 65.3 | 490 |
| Moderate to Mild: Lesion Volume vs. Moderate to Severe: Lesion Volume | 44.1 | 15.25 | 28.85 | 0.3811 | 50 | 50 | 75.71 | 490 |
| Moderate to Mild: Lesion Volume vs. Moderate to Severe: MRI [Segmented] | 44.1 | 14.09 | 30.01 | 0.3811 | 50 | 50 | 78.75 | 490 |
| Moderate to Mild: Lesion Volume vs. Severe: Lesion Volume | 44.1 | 6.205 | 37.9 | 0.3811 | 50 | 50 | 99.44 | 490 |
| Moderate to Mild: Lesion Volume vs. Severe: MRI [Segmented] | 44.1 | 16.73 | 27.38 | 0.3811 | 50 | 50 | 71.83 | 490 |
| Moderate to Mild: MRI [Segmented] vs. Moderate: Lesion Volume | 32.13 | 26.72 | 5.406 | 0.3811 | 50 | 50 | 14.18 | 490 |
| Moderate to Mild: MRI [Segmented] vs. Moderate: MRI [Segmented] | 32.13 | 19.21 | 12.91 | 0.3811 | 50 | 50 | 33.88 | 490 |

Continued Table S8

| | | | | | | | | |
|---|---|---|---|---|---|---|---|---|
| Moderate to Mild: MRI [Segmented] vs. Moderate to Severe: Lesion Volume | 32.13 | 15.25 | 16.88 | 0.3811 | 50 | 50 | 44.29 | 490 |
| Moderate to Mild: MRI [Segmented] vs. Moderate to Severe: MRI [Segmented] | 32.13 | 14.09 | 18.04 | 0.3811 | 50 | 50 | 47.32 | 490 |
| Moderate to Mild: MRI [Segmented] vs. Severe: Lesion Volume | 32.13 | 6.205 | 25.92 | 0.3811 | 50 | 50 | 68.02 | 490 |
| Moderate to Mild: MRI [Segmented] vs. Severe: MRI [Segmented] | 32.13 | 16.73 | 15.4 | 0.3811 | 50 | 50 | 40.41 | 490 |
| Moderate: Lesion Volume vs. Moderate: MRI [Segmented] | 26.72 | 19.21 | 7.507 | 0.3811 | 50 | 50 | 19.7 | 490 |
| Moderate: Lesion Volume vs. Moderate to Severe: Lesion Volume | 26.72 | 15.25 | 11.47 | 0.3811 | 50 | 50 | 30.1 | 490 |
| Moderate: Lesion Volume vs. Moderate to Severe: MRI [Segmented] | 26.72 | 14.09 | 12.63 | 0.3811 | 50 | 50 | 33.14 | 490 |
| Moderate: Lesion Volume vs. Severe: Lesion Volume | 26.72 | 6.205 | 20.52 | 0.3811 | 50 | 50 | 53.83 | 490 |
| Moderate: Lesion Volume vs. Severe: MRI [Segmented] | 26.72 | 16.73 | 9.994 | 0.3811 | 50 | 50 | 26.22 | 490 |
| Moderate: MRI [Segmented] vs. Moderate to Severe: Lesion Volume | 19.21 | 15.25 | 3.967 | 0.3811 | 50 | 50 | 10.41 | 490 |
| Moderate: MRI [Segmented] vs. Moderate to Severe: MRI [Segmented] | 19.21 | 14.09 | 5.124 | 0.3811 | 50 | 50 | 13.44 | 490 |
| Moderate: MRI [Segmented] vs. Severe: Lesion Volume | 19.21 | 6.205 | 13.01 | 0.3811 | 50 | 50 | 34.14 | 490 |
| Moderate: MRI [Segmented] vs. Severe: MRI [Segmented] | 19.21 | 16.73 | 2.487 | 0.3811 | 50 | 50 | 6.527 | 490 |
| Moderate to Severe: Lesion Volume vs. Moderate to Severe: MRI [Segmented] | 15.25 | 14.09 | 1.157 | 0.3811 | 50 | 50 | 3.036 | 490 |
| Moderate to Severe: Lesion Volume vs. Severe: Lesion Volume | 15.25 | 6.205 | 9.043 | 0.3811 | 50 | 50 | 23.73 | 490 |
| Moderate to Severe: Lesion Volume vs. Severe: MRI [Segmented] | 15.25 | 16.73 | -1.479 | 0.3811 | 50 | 50 | 3.881 | 490 |
| Moderate to Severe: MRI [Segmented] vs. Severe: Lesion Volume | 14.09 | 6.205 | 7.886 | 0.3811 | 50 | 50 | 20.69 | 490 |
| Moderate to Severe: MRI [Segmented] vs. Severe: MRI [Segmented] | 14.09 | 16.73 | -2.636 | 0.3811 | 50 | 50 | 6.917 | 490 |
| Severe: Lesion Volume vs. Severe: MRI [Segmented] | 6.205 | 16.73 | -10.52 | 0.3811 | 50 | 50 | 27.61 | 490 |

**Table S8 Full statistical report on the analysis of prediction accuracy and severity grade for the subacute deficit in the replication cohort.**

| Šídák's multiple comparisons test | Mean Diff. | 95% CI of Diff. | Adjusted P Value |
|---|---|---|---|
| Mild: MRI [Segmented] vs. Mild: Lesion Volume | -1.146 | -3.049 to 0.7569 | 0.9769 |
| Mild: MRI [Segmented] vs. Mild: Initial Deficit [Day 2-6] | 19.25 | 17.35 to 21.16 | <0.0001 |
| Mild: MRI [Segmented] vs. Moderate to Mild: MRI [Segmented] | 12.36 | 10.46 to 14.27 | <0.0001 |
| Mild: MRI [Segmented] vs. Moderate to Mild: Lesion Volume | 8.36 | 6.457 to 10.26 | <0.0001 |
| Mild: MRI [Segmented] vs. Moderate to Mild: Initial Deficit [Day 2-6] | 28.03 | 26.12 to 29.93 | <0.0001 |
| Mild: MRI [Segmented] vs. Moderate: MRI [Segmented] | 26.09 | 24.18 to 27.99 | <0.0001 |
| Mild: MRI [Segmented] vs. Moderate: Lesion Volume | 28.09 | 26.19 to 29.99 | <0.0001 |
| Mild: MRI [Segmented] vs. Moderate: Initial Deficit [Day 2-6] | 27.39 | 25.48 to 29.29 | <0.0001 |
| Mild: MRI [Segmented] vs. Moderate to Severe: MRI [Segmented] | 19 | 17.09 to 20.90 | <0.0001 |
| Mild: MRI [Segmented] vs. Moderate to Severe: Lesion Volume | 17.94 | 16.04 to 19.85 | <0.0001 |
| Mild: MRI [Segmented] vs. Moderate to Severe: Initial Deficit [Day 2-6] | 18.97 | 17.06 to 20.87 | <0.0001 |
| Mild: MRI [Segmented] vs. Severe: MRI [Segmented] | -1.716 | -3.618 to 0.1870 | 0.1593 |
| Mild: MRI [Segmented] vs. Severe: Lesion Volume | 15.16 | 13.25 to 17.06 | <0.0001 |
| Mild: MRI [Segmented] vs. Severe: Initial Deficit [Day 2-6] | 3.641 | 1.738 to 5.543 | <0.0001 |
| Mild: Lesion Volume vs. Mild: Initial Deficit [Day 2-6] | 20.4 | 18.50 to 22.30 | <0.0001 |
| Mild: Lesion Volume vs. Moderate to Mild: MRI [Segmented] | 13.51 | 11.61 to 15.41 | <0.0001 |
| Mild: Lesion Volume vs. Moderate to Mild: Lesion Volume | 9.506 | 7.603 to 11.41 | <0.0001 |
| Mild: Lesion Volume vs. Moderate to Mild: Initial Deficit [Day 2-6] | 29.17 | 27.27 to 31.07 | <0.0001 |
| Mild: Lesion Volume vs. Moderate: MRI [Segmented] | 27.23 | 25.33 to 29.13 | <0.0001 |
| Mild: Lesion Volume vs. Moderate: Lesion Volume | 29.24 | 27.33 to 31.14 | <0.0001 |
| Mild: Lesion Volume vs. Moderate: Initial Deficit [Day 2-6] | 28.53 | 26.63 to 30.44 | <0.0001 |
| Mild: Lesion Volume vs. Moderate to Severe: MRI [Segmented] | 20.14 | 18.24 to 22.05 | <0.0001 |

Continued Table S9

| | | | |
|---|---|---|---|
| Mild: Lesion Volume vs. Moderate to Severe: Lesion Volume | 19.09 | 17.19 to 20.99 | <0.0001 |
| Mild: Lesion Volume vs. Moderate to Severe: Initial Deficit [Day 2-6] | 20.11 | 18.21 to 22.02 | <0.0001 |
| Mild: Lesion Volume vs. Severe: MRI [Segmented] | -0.5699 | -2.473 to 1.333 | >0.9999 |
| Mild: Lesion Volume vs. Severe: Lesion Volume | 16.3 | 14.40 to 18.20 | <0.0001 |
| Mild: Lesion Volume vs. Severe: Initial Deficit [Day 2-6] | 4.786 | 2.884 to 6.689 | <0.0001 |
| Mild: Initial Deficit [Day 2-6] vs. Moderate to Mild: MRI [Segmented] | -6.889 | -8.791 to -4.986 | <0.0001 |
| Mild: Initial Deficit [Day 2-6] vs. Moderate to Mild: Lesion Volume | -10.89 | -12.80 to -8.990 | <0.0001 |
| Mild: Initial Deficit [Day 2-6] vs. Moderate to Mild: Initial Deficit [Day 2-6] | 8.773 | 6.870 to 10.68 | <0.0001 |
| Mild: Initial Deficit [Day 2-6] vs. Moderate: MRI [Segmented] | 6.833 | 4.930 to 8.736 | <0.0001 |
| Mild: Initial Deficit [Day 2-6] vs. Moderate: Lesion Volume | 8.838 | 6.935 to 10.74 | <0.0001 |
| Mild: Initial Deficit [Day 2-6] vs. Moderate: Initial Deficit [Day 2-6] | 8.135 | 6.232 to 10.04 | <0.0001 |
| Mild: Initial Deficit [Day 2-6] vs. Moderate to Severe: MRI [Segmented] | -0.2558 | -2.159 to 1.647 | >0.9999 |
| Mild: Initial Deficit [Day 2-6] vs. Moderate to Severe: Lesion Volume | -1.308 | -3.211 to 0.5948 | 0.8219 |
| Mild: Initial Deficit [Day 2-6] vs. Moderate to Severe: Initial Deficit [Day 2-6] | -0.2855 | -2.188 to 1.617 | >0.9999 |
| Mild: Initial Deficit [Day 2-6] vs. Severe: MRI [Segmented] | -20.97 | -22.87 to -19.07 | <0.0001 |
| Mild: Initial Deficit [Day 2-6] vs. Severe: Lesion Volume | -4.098 | -6.000 to -2.195 | <0.0001 |
| Mild: Initial Deficit [Day 2-6] vs. Severe: Initial Deficit [Day 2-6] | -15.61 | -17.51 to -13.71 | <0.0001 |
| Moderate to Mild: MRI [Segmented] vs. Moderate to Mild: Lesion Volume | -4.004 | -5.907 to -2.101 | <0.0001 |
| Moderate to Mild: MRI [Segmented] vs. Moderate to Mild: Initial Deficit [Day 2-6] | 15.66 | 13.76 to 17.56 | <0.0001 |
| Moderate to Mild: MRI [Segmented] vs. Moderate: MRI [Segmented] | 13.72 | 11.82 to 15.62 | <0.0001 |
| Moderate to Mild: MRI [Segmented] vs. Moderate: Lesion Volume | 15.73 | 13.82 to 17.63 | <0.0001 |

Continued Table S9

| | | | |
|---|---|---|---|
| Moderate to Mild: MRI [Segmented] vs. Moderate: Initial Deficit [Day 2-6] | 15.02 | 13.12 to 16.93 | <0.0001 |
| Moderate to Mild: MRI [Segmented] vs. Moderate to Severe: MRI [Segmented] | 6.633 | 4.730 to 8.536 | <0.0001 |
| Moderate to Mild: MRI [Segmented] vs. Moderate to Severe: Lesion Volume | 5.581 | 3.678 to 7.484 | <0.0001 |
| Moderate to Mild: MRI [Segmented] vs. Moderate to Severe: Initial Deficit [Day 2-6] | 6.603 | 4.701 to 8.506 | <0.0001 |
| Moderate to Mild: MRI [Segmented] vs. Severe: MRI [Segmented] | -14.08 | -15.98 to -12.18 | <0.0001 |
| Moderate to Mild: MRI [Segmented] vs. Severe: Lesion Volume | 2.791 | 0.8884 to 4.694 | <0.0001 |
| Moderate to Mild: MRI [Segmented] vs. Severe: Initial Deficit [Day 2-6] | -8.723 | -10.63 to -6.821 | <0.0001 |
| Moderate to Mild: Lesion Volume vs. Moderate to Mild: Initial Deficit [Day 2-6] | 19.67 | 17.76 to 21.57 | <0.0001 |
| Moderate to Mild: Lesion Volume vs. Moderate: MRI [Segmented] | 17.73 | 15.82 to 19.63 | <0.0001 |
| Moderate to Mild: Lesion Volume vs. Moderate: Lesion Volume | 19.73 | 17.83 to 21.63 | <0.0001 |
| Moderate to Mild: Lesion Volume vs. Moderate: Initial Deficit [Day 2-6] | 19.03 | 17.12 to 20.93 | <0.0001 |
| Moderate to Mild: Lesion Volume vs. Moderate to Severe: MRI [Segmented] | 10.64 | 8.734 to 12.54 | <0.0001 |
| Moderate to Mild: Lesion Volume vs. Moderate to Severe: Lesion Volume | 9.585 | 7.682 to 11.49 | <0.0001 |
| Moderate to Mild: Lesion Volume vs. Moderate to Severe: Initial Deficit [Day 2-6] | 10.61 | 8.704 to 12.51 | <0.0001 |
| Moderate to Mild: Lesion Volume vs. Severe: MRI [Segmented] | -10.08 | -11.98 to -8.173 | <0.0001 |
| Moderate to Mild: Lesion Volume vs. Severe: Lesion Volume | 6.795 | 4.892 to 8.698 | <0.0001 |
| Moderate to Mild: Lesion Volume vs. Severe: Initial Deficit [Day 2-6] | -4.72 | -6.622 to -2.817 | <0.0001 |
| Moderate to Mild: Initial Deficit [Day 2-6] vs. Moderate: MRI [Segmented] | -1.94 | -3.842 to -0.03692 | 0.039 |
| Moderate to Mild: Initial Deficit [Day 2-6] vs. Moderate: Lesion Volume | 0.06519 | -1.838 to 1.968 | >0.9999 |

Continued Table S9

| | | | |
|---|---|---|---|
| Moderate to Mild: Initial Deficit [Day 2-6] vs. Moderate: Initial Deficit [Day 2-6] | -0.6379 | -2.541 to 1.265 | >0.9999 |
| Moderate to Mild: Initial Deficit [Day 2-6] vs. Moderate to Severe: MRI [Segmented] | -9.028 | -10.93 to -7.126 | <0.0001 |
| Moderate to Mild: Initial Deficit [Day 2-6] vs. Moderate to Severe: Lesion Volume | -10.08 | -11.98 to -8.178 | <0.0001 |
| Moderate to Mild: Initial Deficit [Day 2-6] vs. Moderate to Severe: Initial Deficit [Day 2-6] | -9.058 | -10.96 to -7.156 | <0.0001 |
| Moderate to Mild: Initial Deficit [Day 2-6] vs. Severe: MRI [Segmented] | -29.74 | -31.64 to -27.84 | <0.0001 |
| Moderate to Mild: Initial Deficit [Day 2-6] vs. Severe: Lesion Volume | -12.87 | -14.77 to -10.97 | <0.0001 |
| Moderate to Mild: Initial Deficit [Day 2-6] vs. Severe: Initial Deficit [Day 2-6] | -24.38 | -26.29 to -22.48 | <0.0001 |
| Moderate: MRI [Segmented] vs. Moderate: Lesion Volume | 2.005 | 0.1021 to 3.908 | 0.0249 |
| Moderate: MRI [Segmented] vs. Moderate: Initial Deficit [Day 2-6] | 1.302 | -0.6009 to 3.204 | 0.8313 |
| Moderate: MRI [Segmented] vs. Moderate to Severe: MRI [Segmented] | -7.089 | -8.992 to -5.186 | <0.0001 |
| Moderate: MRI [Segmented] vs. Moderate to Severe: Lesion Volume | -8.141 | -10.04 to -6.238 | <0.0001 |
| Moderate: MRI [Segmented] vs. Moderate to Severe: Initial Deficit [Day 2-6] | -7.119 | -9.021 to -5.216 | <0.0001 |
| Moderate: MRI [Segmented] vs. Severe: MRI [Segmented] | -27.8 | -29.70 to -25.90 | <0.0001 |
| Moderate: MRI [Segmented] vs. Severe: Lesion Volume | -10.93 | -12.83 to -9.028 | <0.0001 |
| Moderate: MRI [Segmented] vs. Severe: Initial Deficit [Day 2-6] | -22.45 | -24.35 to -20.54 | <0.0001 |
| Moderate: Lesion Volume vs. Moderate: Initial Deficit [Day 2-6] | -0.7031 | -2.606 to 1.200 | >0.9999 |
| Moderate: Lesion Volume vs. Moderate to Severe: MRI [Segmented] | -9.094 | -11.00 to -7.191 | <0.0001 |
| Moderate: Lesion Volume vs. Moderate to Severe: Lesion Volume | -10.15 | -12.05 to -8.243 | <0.0001 |
| Moderate: Lesion Volume vs. Moderate to Severe: Initial Deficit [Day 2-6] | -9.123 | -11.03 to -7.221 | <0.0001 |

Continued Table S9

| | | | |
|---|---|---|---|
| Moderate: Lesion Volume vs. Severe: MRI [Segmented] | -29.81 | -31.71 to -27.90 | <0.0001 |
| Moderate: Lesion Volume vs. Severe: Lesion Volume | -12.94 | -14.84 to -11.03 | <0.0001 |
| Moderate: Lesion Volume vs. Severe: Initial Deficit [Day 2-6] | -24.45 | -26.35 to -22.55 | <0.0001 |
| Moderate: Initial Deficit [Day 2-6] vs. Moderate to Severe: MRI [Segmented] | -8.391 | -10.29 to -6.488 | <0.0001 |
| Moderate: Initial Deficit [Day 2-6] vs. Moderate to Severe: Lesion Volume | -9.443 | -11.35 to -7.540 | <0.0001 |
| Moderate: Initial Deficit [Day 2-6] vs. Moderate to Severe: Initial Deficit [Day 2-6] | -8.42 | -10.32 to -6.518 | <0.0001 |
| Moderate: Initial Deficit [Day 2-6] vs. Severe: MRI [Segmented] | -29.1 | -31.01 to -27.20 | <0.0001 |
| Moderate: Initial Deficit [Day 2-6] vs. Severe: Lesion Volume | -12.23 | -14.14 to -10.33 | <0.0001 |
| Moderate: Initial Deficit [Day 2-6] vs. Severe: Initial Deficit [Day 2-6] | -23.75 | -25.65 to -21.84 | <0.0001 |
| Moderate to Severe: MRI [Segmented] vs. Moderate to Severe: Lesion Volume | -1.052 | -2.955 to 0.8506 | 0.9968 |
| Moderate to Severe: MRI [Segmented] vs. Moderate to Severe: Initial Deficit [Day 2-6] | -0.02972 | -1.932 to 1.873 | >0.9999 |
| Moderate to Severe: MRI [Segmented] vs. Severe: MRI [Segmented] | -20.71 | -22.62 to -18.81 | <0.0001 |
| Moderate to Severe: MRI [Segmented] vs. Severe: Lesion Volume | -3.842 | -5.745 to -1.939 | <0.0001 |
| Moderate to Severe: MRI [Segmented] vs. Severe: Initial Deficit [Day 2-6] | -15.36 | -17.26 to -13.45 | <0.0001 |
| Moderate to Severe: Lesion Volume vs. Moderate to Severe: Initial Deficit [Day 2-6] | 1.022 | -0.8803 to 2.925 | 0.9985 |
| Moderate to Severe: Lesion Volume vs. Severe: MRI [Segmented] | -19.66 | -21.56 to -17.76 | <0.0001 |
| Moderate to Severe: Lesion Volume vs. Severe: Lesion Volume | -2.79 | -4.692 to -0.8870 | <0.0001 |
| Moderate to Severe: Lesion Volume vs. Severe: Initial Deficit [Day 2-6] | -14.3 | -16.21 to -12.40 | <0.0001 |
| Moderate to Severe: Initial Deficit [Day 2-6] vs. Severe: MRI [Segmented] | -20.68 | -22.59 to -18.78 | <0.0001 |

Continued Table S9

| | | | |
|---|---|---|---|
| Moderate to Severe: Initial Deficit [Day 2-6] vs. Severe: Lesion Volume | -3.812 | -5.715 to -1.909 | <0.0001 |
| Moderate to Severe: Initial Deficit [Day 2-6] vs. Severe: Initial Deficit [Day 2-6] | -15.33 | -17.23 to -13.42 | <0.0001 |
| Severe: MRI [Segmented] vs. Severe: Lesion Volume | 16.87 | 14.97 to 18.77 | <0.0001 |
| Severe: MRI [Segmented] vs. Severe: Initial Deficit [Day 2-6] | 5.356 | 3.454 to 7.259 | <0.0001 |
| Severe: Lesion Volume vs. Severe: Initial Deficit [Day 2-6] | -11.51 | -13.42 to -9.612 | <0.0001 |

| **Test details** | **Mean 1** | **Mean 2** | **Mean Diff.** | **SE of Diff.** | **N1** | **N2** | **t** | **DF** |
|---|---|---|---|---|---|---|---|---|
| Mild: Subacute Deficit vs. Mild: MRI [Segmented] | 15.38 | 34.64 | -19.25 | 0.5432 | 50 | 50 | 35.44 | 735 |
| Mild: Subacute Deficit vs. Mild: Lesion Volume | 15.38 | 35.78 | -20.4 | 0.5432 | 50 | 50 | 37.55 | 735 |
| Mild: Subacute Deficit vs. Moderate to Mild: Subacute Deficit | 15.38 | 6.611 | 8.773 | 0.5432 | 50 | 50 | 16.15 | 735 |
| Mild: Subacute Deficit vs. Moderate to Mild: MRI [Segmented] | 15.38 | 22.27 | -6.889 | 0.5432 | 50 | 50 | 12.68 | 735 |
| Mild: Subacute Deficit vs. Moderate to Mild: Lesion Volume | 15.38 | 26.28 | -10.89 | 0.5432 | 50 | 50 | 20.05 | 735 |
| Mild: Subacute Deficit vs. Moderate: Subacute Deficit | 15.38 | 7.248 | 8.135 | 0.5432 | 50 | 50 | 14.98 | 735 |
| Mild: Subacute Deficit vs. Moderate: MRI [Segmented] | 15.38 | 8.55 | 6.833 | 0.5432 | 50 | 50 | 12.58 | 735 |
| Mild: Subacute Deficit vs. Moderate: Lesion Volume | 15.38 | 6.545 | 8.838 | 0.5432 | 50 | 50 | 16.27 | 735 |
| Mild: Subacute Deficit vs. Moderate to Severe: Subacute Deficit | 15.38 | 15.67 | -0.2855 | 0.5432 | 50 | 50 | 0.5256 | 735 |
| Mild: Subacute Deficit vs. Moderate to Severe: MRI [Segmented] | 15.38 | 15.64 | -0.2558 | 0.5432 | 50 | 50 | 0.4709 | 735 |
| Mild: Subacute Deficit vs. Moderate to Severe: Lesion Volume | 15.38 | 16.69 | -1.308 | 0.5432 | 50 | 50 | 2.408 | 735 |
| Mild: Subacute Deficit vs. Severe: Subacute Deficit | 15.38 | 31 | -15.61 | 0.5432 | 50 | 50 | 28.74 | 735 |
| Mild: Subacute Deficit vs. Severe: MRI [Segmented] | 15.38 | 36.35 | -20.97 | 0.5432 | 50 | 50 | 38.6 | 735 |
| Mild: Subacute Deficit vs. Severe: Lesion Volume | 15.38 | 19.48 | -4.098 | 0.5432 | 50 | 50 | 7.543 | 735 |

Continued Table S9

| | | | | | | | | |
|---|---|---|---|---|---|---|---|---|
| Mild: MRI [Segmented] vs. Mild: Lesion Volume | 34.64 | 35.78 | -1.146 | 0.5432 | 50 | 50 | 2.109 | 735 |
| Mild: MRI [Segmented] vs. Moderate to Mild: Subacute Deficit | 34.64 | 6.611 | 28.03 | 0.5432 | 50 | 50 | 51.59 | 735 |
| Mild: MRI [Segmented] vs. Moderate to Mild: MRI [Segmented] | 34.64 | 22.27 | 12.36 | 0.5432 | 50 | 50 | 22.76 | 735 |
| Mild: MRI [Segmented] vs. Moderate to Mild: Lesion Volume | 34.64 | 26.28 | 8.36 | 0.5432 | 50 | 50 | 15.39 | 735 |
| Mild: MRI [Segmented] vs. Moderate: Subacute Deficit | 34.64 | 7.248 | 27.39 | 0.5432 | 50 | 50 | 50.42 | 735 |
| Mild: MRI [Segmented] vs. Moderate: MRI [Segmented] | 34.64 | 8.55 | 26.09 | 0.5432 | 50 | 50 | 48.02 | 735 |
| Mild: MRI [Segmented] vs. Moderate: Lesion Volume | 34.64 | 6.545 | 28.09 | 0.5432 | 50 | 50 | 51.71 | 735 |
| Mild: MRI [Segmented] vs. Moderate to Severe: Subacute Deficit | 34.64 | 15.67 | 18.97 | 0.5432 | 50 | 50 | 34.92 | 735 |
| Mild: MRI [Segmented] vs. Moderate to Severe: MRI [Segmented] | 34.64 | 15.64 | 19 | 0.5432 | 50 | 50 | 34.97 | 735 |
| Mild: MRI [Segmented] vs. Moderate to Severe: Lesion Volume | 34.64 | 16.69 | 17.94 | 0.5432 | 50 | 50 | 33.03 | 735 |
| Mild: MRI [Segmented] vs. Severe: Subacute Deficit | 34.64 | 31 | 3.641 | 0.5432 | 50 | 50 | 6.702 | 735 |
| Mild: MRI [Segmented] vs. Severe: MRI [Segmented] | 34.64 | 36.35 | -1.716 | 0.5432 | 50 | 50 | 3.158 | 735 |
| Mild: MRI [Segmented] vs. Severe: Lesion Volume | 34.64 | 19.48 | 15.16 | 0.5432 | 50 | 50 | 27.9 | 735 |
| Mild: Lesion Volume vs. Moderate to Mild: Subacute Deficit | 35.78 | 6.611 | 29.17 | 0.5432 | 50 | 50 | 53.7 | 735 |
| Mild: Lesion Volume vs. Moderate to Mild: MRI [Segmented] | 35.78 | 22.27 | 13.51 | 0.5432 | 50 | 50 | 24.87 | 735 |
| Mild: Lesion Volume vs. Moderate to Mild: Lesion Volume | 35.78 | 26.28 | 9.506 | 0.5432 | 50 | 50 | 17.5 | 735 |
| Mild: Lesion Volume vs. Moderate: Subacute Deficit | 35.78 | 7.248 | 28.53 | 0.5432 | 50 | 50 | 52.53 | 735 |
| Mild: Lesion Volume vs. Moderate: MRI [Segmented] | 35.78 | 8.55 | 27.23 | 0.5432 | 50 | 50 | 50.13 | 735 |
| Mild: Lesion Volume vs. Moderate: Lesion Volume | 35.78 | 6.545 | 29.24 | 0.5432 | 50 | 50 | 53.82 | 735 |

Continued Table S9

| | | | | | | | | |
|---|---|---|---|---|---|---|---|---|
| Mild: Lesion Volume vs. Moderate to Severe: Subacute Deficit | 35.78 | 15.67 | 20.11 | 0.5432 | 50 | 50 | 37.03 | 735 |
| Mild: Lesion Volume vs. Moderate to Severe: MRI [Segmented] | 35.78 | 15.64 | 20.14 | 0.5432 | 50 | 50 | 37.08 | 735 |
| Mild: Lesion Volume vs. Moderate to Severe: Lesion Volume | 35.78 | 16.69 | 19.09 | 0.5432 | 50 | 50 | 35.14 | 735 |
| Mild: Lesion Volume vs. Severe: Subacute Deficit | 35.78 | 31 | 4.786 | 0.5432 | 50 | 50 | 8.811 | 735 |
| Mild: Lesion Volume vs. Severe: MRI [Segmented] | 35.78 | 36.35 | -0.5699 | 0.5432 | 50 | 50 | 1.049 | 735 |
| Mild: Lesion Volume vs. Severe: Lesion Volume | 35.78 | 19.48 | 16.3 | 0.5432 | 50 | 50 | 30.01 | 735 |
| Moderate to Mild: Subacute Deficit vs. Moderate to Mild: MRI [Segmented] | 6.611 | 22.27 | -15.66 | 0.5432 | 50 | 50 | 28.83 | 735 |
| Moderate to Mild: Subacute Deficit vs. Moderate to Mild: Lesion Volume | 6.611 | 26.28 | -19.67 | 0.5432 | 50 | 50 | 36.2 | 735 |
| Moderate to Mild: Subacute Deficit vs. Moderate: Subacute Deficit | 6.611 | 7.248 | -0.6379 | 0.5432 | 50 | 50 | 1.174 | 735 |
| Moderate to Mild: Subacute Deficit vs. Moderate: MRI [Segmented] | 6.611 | 8.55 | -1.94 | 0.5432 | 50 | 50 | 3.571 | 735 |
| Moderate to Mild: Subacute Deficit vs. Moderate: Lesion Volume | 6.611 | 6.545 | 0.06519 | 0.5432 | 50 | 50 | 0.12 | 735 |
| Moderate to Mild: Subacute Deficit vs. Moderate to Severe: Subacute Deficit | 6.611 | 15.67 | -9.058 | 0.5432 | 50 | 50 | 16.68 | 735 |

**Table S9 Full statistical report on the analysis of prediction accuracy and severity grade for the residual deficit in the replication cohort.**
